# ADAMTS-family protease MIG-17 regulates synaptic allometry by modifying the extracellular matrix and modulating glia morphology during growth

**DOI:** 10.1101/734830

**Authors:** Tingting Ji, Kai Wang, Jiale Fan, Jichang Huang, Mengqing Wang, Xiaohua Dong, Yanjun Shi, Laura Manning, Xumin Zhang, Zhiyong Shao, Daniel A. Colón-Ramos

**Author notes:** These authors contribute equally (Tingting Ji, Kai Wang, Jiale Fan). **Corresponding authors:** Zhiyong Shao, Institute of Brain Science, Fudan University, 131 DongAn Road, Mingdao Building 1016, Shanghai, Shanghai, 200032, China, Daniel A. Colón-Ramos, Program in Cellular Neuroscience, Neurodegeneration and Repair, Department of Neuroscience and Department of Cell Biology, Yale University, School of Medicine, 333 Cedar Street, SHM B 163D, New Haven, CT 06510.

## Abstract

Synapses are largely established during embryogenesis and maintained during growth. The mechanisms that regulate synaptic allometry—the maintenance of synaptic positions during growth—are largely unknown. We performed forward genetic screens in *C. elegans* for synaptic allometry mutants and identified *mig-17*, a secreted metalloprotease of the conserved ADAMTS family. Through proteomic mass spectrometry analyses, cell biological and genetic studies we determined that MIG-17 is expressed by muscle cells to modulate glia location and morphology. Glia are proximal to synapses, and the glial location and morphology determine synaptic position during growth. *Mig-17* regulates synapse allometry by influencing epidermal-glia crosstalk through the regulation of basement membrane proteins, including collagen type IV, SPARC and fibulin. Our findings underscore the importance of glia location in the maintenance of synaptic allometry, and uncover a muscle-epidermal-glia signaling axis, mediated through the extracellular matrix, in the regulation of glia morphology and synaptic positions during growth.

## INTRODUCTION

Nervous system architecture and function depend on precise connectivity between pre- and post-synaptic partners. Circuit architecture can also be maintained during the lifetime of the organism (Benard and Hobert, 2009). The mechanisms that preserve synaptic specificity during post-embryonic growth remain largely unknown.

Most of our understanding of synaptic specificity comes from developmental studies examining precise positioning of synapses during their biogenesis (Rawson et al., 2017;Park et al., 2018;Kurshan and Shen, 2019). From these studies we know that precise connectivity during development occurs through orchestrated signaling across multiple tissues. For example, *in vivo* studies have revealed that while cell-cell recognition and signaling between synaptic partners is important for synaptogenesis, non-neuronal cells also guide synaptic specificity (Colon-Ramos, 2009;Sanes and Yamagata, 2009;Margeta and Shen, 2010;Shimozono et al., 2019). During development, synaptic specificity can be instructed by guidepost cells through the secretion of positional cues in the form of morphogenic extracellular signaling molecules (Ullian et al., 2001;Shen and Bargmann, 2003;Colon-Ramos et al., 2007;Ango et al., 2008;Eroglu and Barres, 2010;Tsai et al., 2012;Molofsky et al., 2014).

Post-embryonic maintenance of synapses has been mainly examined at the level of identifying molecular factors necessary for maintaining synaptic stability, density and morphology (Lin and Koleske, 2010;Luo et al., 2014;Cherra and Jin, 2016;Sytnyk et al., 2017;Burden et al., 2018;Hasan and Singh, 2019). Less is known about factors required for maintaining the position of synapses, particularly during post-embryonic growth. As an animal grows, organs scale in different proportions relative to body size, a fact that has long been recognized by biologists and is termed “allometry” (Huxley, 1924;Huxley J, 1936). For example, brain neocortical white matter and neo-cortical grey matter scale different from each other, indicating that specific sub-structures of the brain scale allometrically to total brain size (de Jong et al., 2017). It remains largely unknown, as different tissues disproportionately scale in size during organismal growth, how embryonically-derived synaptic distribution is retained to maintain circuit architecture.

Like synapses, axon positions are also maintained during growth, and genetic studies in *C. elegans* have identified molecules specifically required for this process, such as L1-CAM, F-spondin and the ecto-domain of the FGF Receptor (Aurelio et al., 2002;Aurelio et al., 2003;Hobert and Bulow, 2003;Bulow et al., 2004;Benard et al., 2006;Pocock et al., 2008;Woo et al., 2008;Zhou et al., 2008;Benard et al., 2009;Benard et al., 2012;Noblett et al., 2019;Ramirez-Suarez et al., 2019). These studies uncover two important features regarding maintenance of axon positions during growth. First, the signaling pathways required for maintaining axon positions are different from those required for establishing axon positions during development. Second, these studies suggest that secreted factors in the extracellular matrix (ECM) play important roles in axon position maintenance.

The ECM is a network of macromolecules important for cell-cell interactions, signaling and maintenance of tissue morphogenesis (Jayadev and Sherwood, 2017;Song and Dityatev, 2018). The ECM is a dynamic structure that remodels in part through the activity of ADAMTS metalloproteases (Kelwick et al., 2015). Genetic studies in *C. elegans*, *Drosophila* and humans highlight the importance of ADAMTS metalloproteases for post-embryonic development (Jafari et al., 2010;Meyer et al., 2014;Skeath et al., 2017;Mead and Apte, 2018). Human genetic disorders that affect ADAMTS metalloproteases result in neurodegenerative disorders (Gottschall and Howell, 2015;Rivera et al., 2019), vascular diseases (Zhong and Khalil, 2019), birth defects and short stature, among other diseases (Binder et al., 2017;Mead and Apte, 2018). The ECM is also an important structure for the development and maintenance of neuromuscular junctions (NMJ), and disruption of ECM components, including ADAMTS metalloproteases, affects post-embryonic maintenance of NMJ morphology (Singhal and Martin, 2011;Kurshan et al., 2014;Qin et al., 2014;Dear et al., 2016;Cescon et al., 2018;Heikkinen et al., 2019) The role of ECM in the maintenance of CNS neuron-neuron synapses remains less understood (Heikkinen et al., 2014;Krishnaswamy et al., 2019), particularly during brain allometric growth.

The nematode *C. elegans* provides a tractable genetic model for examining questions related to synaptic allometry—how maintenance of correct synaptic contact, and prevention of formation of inappropriate contacts, are regulated during growth to preserve circuit architecture (Shao et al., 2013). *C. elegans* hatches from its egg as a miniature version of the adult, and grows two orders of magnitude in volume during post-embryonic growth (Knight et al., 2002). The architecture of the nervous system, which is laid out in embryogenesis, is largely preserved during this process (Benard and Hobert, 2009). Single neurons of known identity can be tracked during the lifetime of the organism using cell-specific promoters, along with *in vivo* probes for visualization of their synaptic positions (Nonet, 1999;Colon-Ramos et al., 2007).

We established a system in *C. elegans* to study synaptic allometry *in vivo*, and from forward genetic screens identified *cima-1* as a gene required for synaptic allometry (Shao et al., 2013). In *cima-1* mutants, synaptic contacts are correctly established during embryogenesis, but ectopic synapses emerge as the animals grow. *cima-1* encodes a novel solute carrier in the SLC17 family of transporters that includes Sialin, a protein that when mutated in humans results in neurological disorders (Verheijen et al., 1999). Rather than functioning in neurons, *cima-1* functions in the nearby epidermal cells to antagonize the FGF Receptor, most likely by inhibiting its role in epidermal-glia adhesion. Therefore, *cima-1* functions in non-neuronal cells during post-embryonic growth to preserve synaptic positions through glia (Shao et al., 2013).

To further determine the cellular and molecular mechanisms that regulate synaptic allometry, we performed suppressor forward genetic screens in the *cima-1* mutant background, and identified *mig-17*, encoding a secreted ADAMTS metalloprotease (Nishiwaki et al., 2000). We find that muscle-derived *mig-17* modulates basement membrane proteins. The basement membrane is not in direct contact with the affected synapses. Instead, muscle-derived basement membrane coats the apical side of glia, while glia contact synapses on their basal side. MIG-17 is regulated during growth, and remodels the basement membrane to modulate glia morphology, which in turn modulates synaptic positions during growth. Our findings underscore the *in vivo* importance of non-neuronal cells in the maintenance of synaptic allometry. Our findings also uncover a muscle-epidermal-glia signaling axis, modulated by *mig-17* and the ECM, in regulating synaptic allometry during growth.

## MATERIALS AND METHODS

### Strains

All strains were grown at 22°C on NGM agar plates seeded with *Escherichia coli* OP50 (Brenner, 1974) unless specified. *C. elegans* N2 bristol was used as the wild-type strain.

The following mutant alleles were utilized in this study: LGI: *cle-1(cg120)*

LGII: *unc-52(gk3)*

LGIII: *emb-9(tk75), emb-9(xd51), ina-1(gm144)*

LGIV: *cima-1(wy84)*, *fbl-1(k201), ost-1(gk193465), ost-1(gk786697)*

LGV: *mig-17(ola226)*, *mig-17(k113)*, *mig-17(shc8), mig-17(shc19)*, *nid-1(cg118), nid-1(cg119)*

LGX: *let-2(k193)*, *let-2(b246)*, *egl-15(n484)*, *sdn-1(zh20)*

The following transgenic lines were used in this study:

*shcEx1126, shcEx1127* and *shcEx1128[Pttx-3::syd-1::GFP;Pttx-3::rab-3::mCherry;Punc-122::RFP], shcEx1146* and *shcEx1147[Pmig-17::mig-17 genomics;Phlh-17::mCherry], shcEx1129[Pmig-17::mig-17::SL2::GFP;Pdpy-4::mCherry], shcEx1130[Pmig-17::mig-17::SL2::GFP;Pmyo-3::mCherry], shcEx1131[Pmig-17::mig-17::SL2::GFP;Phlh-17::mCherry], shcEx1410[Pmig-17::mig-17::SL2::GFP;Prab-3::mCherry], shcEx845[Phlh-17::mCherry], shcEx1145[Pdpy-4::mCherry], shcEx1402[Pmyo-3::mCherry], shcEx1403[Prab-3::mCherry], shcEx1414* and *shcEx1415 [Pmig-17::mig-17(E303A); Phlh-17::mCherry], shcEx1133, shcEx1134* and *shcEx1135[Pmyo-3::mig-17;Phlh-17::mCherry], shcEx1136* and *shcEx1137[Punc-14::mig-17;Phlh-17::mCherry], shcEx1139* and *shcEx1140[Phlh-17::mig-17;Phlh-17::mCherry], shcEx1142* and *shcEx1143[Pdpy-7::mig-17;Punc-122::GFP], qyIs46[unc119;emb-9::mCherry], shcEx776, shcEx777, shcEx778, shcEx780* and *shcEx781[Phlh-17::mCherry;Pttx-3::GFP::rab-3], shcEx424, shcEx425, shcEx536, shcEx537* and *shcEx538[Pdpy-7::egl-15(5A);Phlh-17::mCherry;Pttx-3::GFP:: rab-3], shcEx1252* and *shcEx1253 [Pmig-17::mig-17(genomic);Phlh-17::mCherry].*

Details on strains used in this study are listed in Table S1.

### EMS Screen and mutant identification

To identify *cima-1* suppressors, animals that exhibited normal presynaptic distribution were isolated from a forward Ethyl Methane-Sulphonate (EMS) screen performed on the *cima-1(wy84)* mutants. The suppressor *ola226* was isolated from this screen. The causative genetic lesion was identified through SNP mapping and whole genome sequencing (Minevich et al., 2012) to be a G to A point mutation in the first exon of *mig-17*, turning E19 into K in the protein. Fosmid WRM0616aB07, which includes the *mig-17* gene, rescues the observed suppression of the AIY presynaptic distribution in *cima-1(wy84); ola226*.

### Germline Transformation

Transformations were carried out by microinjection of plasmid DNA into the gonad of adult hermaphrodites (Mello et al., 1991). Plasmids were injected with 5-20ng/μl.

### Plasmids

The following constructs were created by Gateway cloning (Invitrogen): P*mig-17::SL2::GFP;* P*mig-17::mig-17(E303A)::GFP;* P*hlh-17::mig-17;* P*unc-14::mig-17;* P*dpy-7::mig-17;* P*myo-3::mig-17.* The *mig-17* promoter is 1.7kb sequence upstream from the start codon. The remaining constructs are listed in Table S2. Detailed cloning information is available upon request.

We constructed two Cas9-sgRNAs with pDD162 for each strain according to the method in (Dickinson et al., 2015). The repair template of *mig-17::mNeonGreen* was modified from pDD268 and is illustrated in Figure S6A. Briefly, *mNeonGreen* was flanked by 1.2kb genomic sequence upstream or downstream of *mig-17* stop codon. To prevent Cas9 from cutting the donor template, we also introduced one synonymous mutation in the protospacer adjacent motif (PAM). The repair template of *mig-17(E303A)* includes 1.2 kb upstream and 1.2 kb downstream of *mig-17* genomic sequence, which flank the Glutamic acid at 303 site. We mutated the Glutamic acid (GAA) to Alanine (GCA) and introduced 8 synonymous mutations to prevent Cas9 from cutting the donor template (Figure S6B). *mig-17(E303A)* point mutatio*n* or *mig-17::mNeonGreen* knock-in animals were generated by microinjection of 50 ng/μl Cas9-sgRNA plasmids, 20ng/μl repair template, and 5ng/μl *Pmyo-3::mCherry* as a co-injection marker. The engineered strains were screened by PCR and verified by Sanger sequencing.

### Protein extraction, digestion, and labeling

The samples were lysed in buffer (8 M guanidine hydrochloride, 100 mM TEAB) and sonicated. Samples were then centrifuged at 20,000g for 30 min at 4°C, and the supernatant collected. Proteins were submitted to reduction by incubation with 10 mM DTT at 37 °C for 45 min, followed by alkylation using 100 mM acrylamide for 1 h at room temperature and digestion with Lys-C and trypsin using the FASP method (Wisniewski et al., 2009). After stable isotope dimethyl labeling in 100 mM TEAB, peptides were mixed with light, intermediate and heavy (formaldehyde and NaBH3CN) isotopic reagents (1:1:1), respectively (Boersema et al., 2009). The peptide mixtures were desalted on a Poros R3 microcolumn according to the previous method (Huang et al., 2018).

### Liquid chromatography–tandem mass spectrometry (LC-MS/MS)

LC-ESI-MS/MS analyses were performed using an LTQ Orbitrap Elite mass spectrometer (Thermo Fisher Scientific, Bremen, Germany) coupled with a nanoflow EASY-nLC 1000 system (Thermo Fisher Scientific, Odense, Denmark). A two-column system was adopted for proteomic analysis. The mobile phases were in Solvent A (0.1% formic acid in H2O) and Solvent B (0.1% formic acid in ACN). The derivatized peptides were eluted using the following gradients: 2-5% B in 2 min, 5-28% B in 98 min, 28-35% B in 5 min, 35-90% B in 2 min, 90% B for 13 min at a flow rate of 200 nl/min. Data-dependent analyses were used in MS analyses. The top 15 abundant ions in each MS scan were selected and fragmented in HCD mode.

Raw data was processed by Proteome Discover (Version 1.4, Thermo Fisher Scientific, Germany) and matched to the *C. elegans* database (20161228, 17,392 sequences) through the Mascot server (Version 2.3, Matrix Science, London, UK). Data was searched using the following parameters: 10 ppm mass tolerance for MS and 0.05 Da for MS/MS fragment ions; up to two missed cleavage sites were allowed; carbamidomethylation on cysteine, dimethyl labeling as fixed modifications; oxidation on methionine as variable modifications. The incorporated Target Decoy PSM Validator in Proteome Discoverer was used to validate the search results with only the hits with FDR≤ 0.01.

### Microscopy and image analyses

Animals were anaesthetized with 50mM Muscimol (Tocris) on 2% agarose pads (Biowest, Lot No.: 111860), and examined with either with Perkin Elmer or Andor Dragonfly Spinning-Disk Confocal Microscope Systems. Image processing was performed by using Image J, Adobe Photoshop CS6 or Imaris software (Andor).

### Quantification

To quantify the percentage of animals with ectopic pre-synapses of AIY Zone 1 and posterior extension of glia, animals were synchronized by being selected at larva stage 4 (L4), and then examined 24 hours later using a Nikon Ni-U fluorescent microscope. Each dataset was collected from at least three biological replicates. At least 20 animals were scored for each group. For each germline transformation, multiple transgenic lines were examined. For synaptic allometric quantification, the ectopic synapses were defined as the presence of synaptic fluorescent markers the AIY Zone 1 region, an asynaptic area in wild type AIY neurons (Colon-Ramos et al., 2007;Shao et al., 2013). We also quantified the ratio of ventral length to total synaptic length (Shao et al. 2013). The overlap of VCSC glia and ectopic synapses was defined as the VCSC glia and synaptic area of overlap at the Zone 1 and Zone 2 regions. The length of VCSC glial cilia and ventral process (a and b in Figure 2A) were measured from confocal images taken in synchronized one-day old adults. The length of the pharynx and the body length were measured via DIC microscopy performed in synchronized one-day old adults.

**Figure 1.**
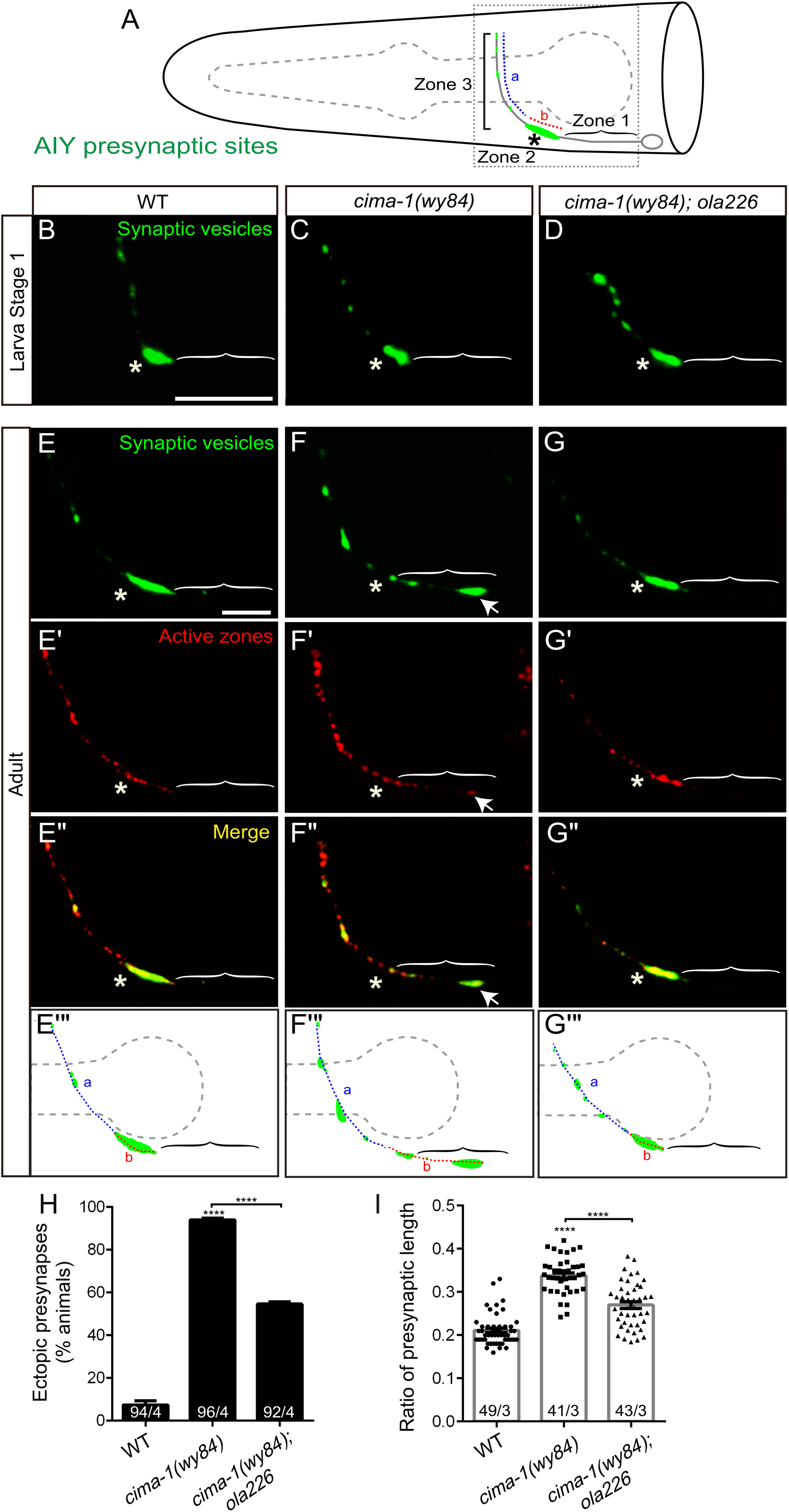
*ola226* suppresses *cima-1 (wy84)* synaptic allometry defects. **(A)** Cartoon diagram of the distribution of presynaptic sites in the AIY interneurons of the nematode *C. elegans*. The head of *C. elegans* (solid black lines), and the pharynx (dashed grey line) are outlined. A single AIY interneuron is depicted in gray, with an oval representing the cell body and a solid gray line representing the neurite. Presynaptic puncta are green. The AIY neurites can be subdivided to three zones: an asynaptic region proximal to the cell body called Zone 1, a synapse-rich region called Zone 2 (asterisk) and a region with sparse synapses, called Zone 3. The red (b) and blue (a) dashed lines represent synaptic distribution, and correspond to Zone 2 and 3 (respectively) in wild type animals. The dotted box represents the region of the head imaged in B-G”. **(B-G’’)** Confocal micrograph images of the AIY presynaptic sites labeled with the synaptic vesicle marker mCherry::RAB-3 (pseudo-colored green, B-G) and active zone protein GFP::SYD-1 (pseudo-colored red, E’-G’) in larva stage 1 animals (B-D) or adult animals (E-G”) for wild type (B, E, E’, E”), *cima-1(wy84)* mutants (C, F, F’, F”) or *cima-1(wy84); ola226* (D, G, G’, G”). Merged images displaying co-localization of synaptic vesicle marker mCherry::RAB-3 and active zone protein GFP::SYD-1 in (E”-G”). Schematic diagrams of the observations are depicted in (E’’’-G’’’). Scale bars, 10μm. Note that the size of the L1 animal is ∼4x smaller than the adult, but the synaptic pattern is similar. Brackets: Zone 1 region; Asterisk: Zone 2 region; Arrows: ectopic synapses in Zone 1 region. **(H)** Quantification of the percentage of animals displaying ectopic AIY presynaptic sites in the Zone 1 region for indicated genotypes. **(I)** Quantification of the ratio of ventral synaptic length (see red (b) in schematic in (A and E’’’-G’’’) to total synaptic region (sum of the length of blue (a) and red (b) in schematic in (A and E’’’-G’’’)). The total number of animals (N) and the number of times scored (n) are indicated in each bar for each genotype as N/n. Error bars represent SEM. Statistical analyses are based on two-tailed student’s t-test, **** p<0.0001 as compared to wild type (if on top of bar graph), unless brackets are used between two compared genotypes.

**Figure 2.**
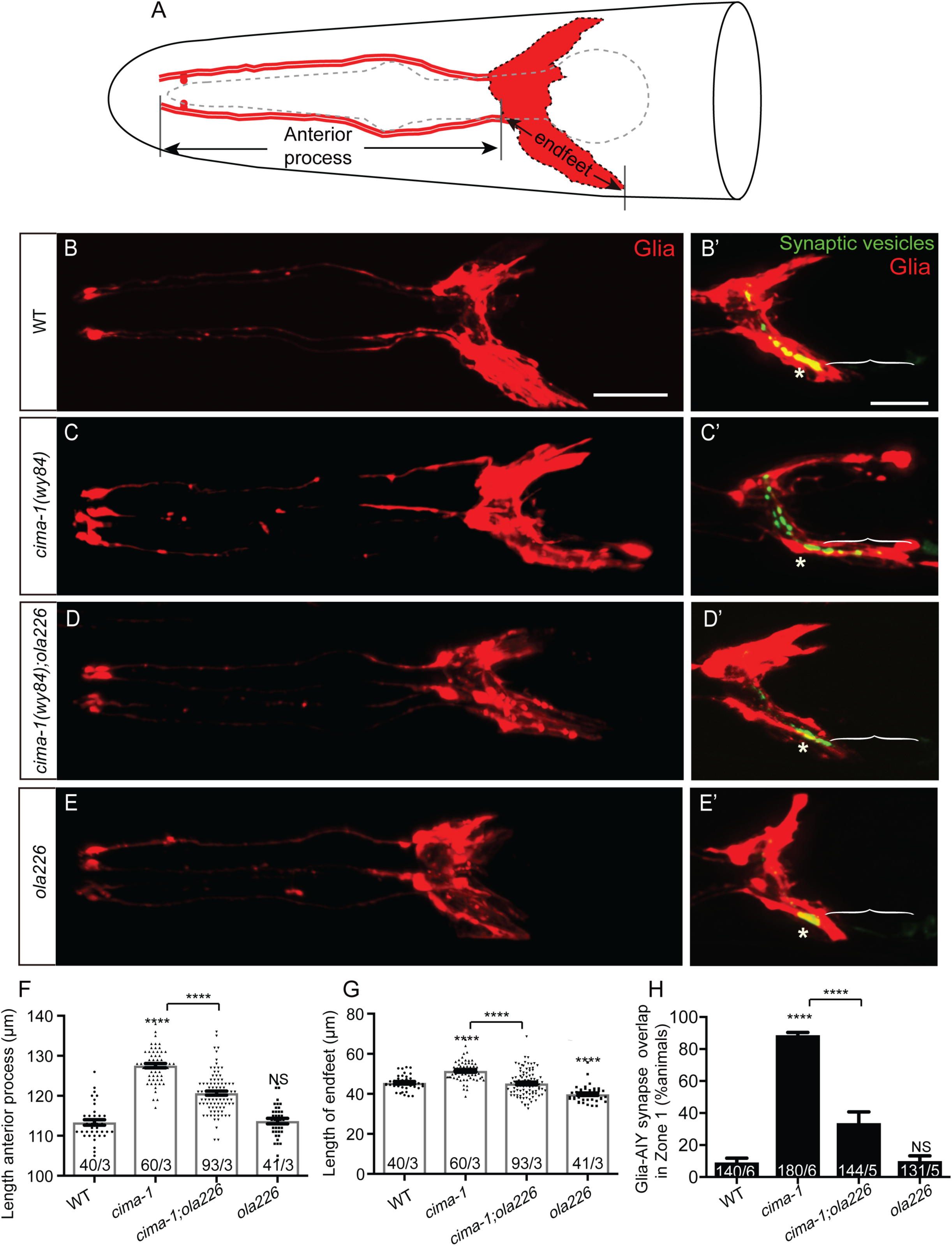
Glia morphology is affected in *ola226* mutants. **(A)** Cartoon diagram of the ventral and dorsal cephalic sheath cell glia (red) in the *C. elegans* head. The ventral cephalic sheath cell (VCSC) glia, which is the lower one labeled in the schematic, contacts the AIY synapses. **(B-E’)** Confocal micrographs of the morphology of VCSC glia and the anterior process (red, labeled with *Phlh-17::mCherry*, B-E), or of VCSC glia cell body and endfeet (red) with the AIY presynaptic marker (green, GFP::RAB-3, B’-E’) in adult wild type (B, B’), *cima-1(wy84)* mutants (C, C’), *cima-1(wy84);ola226* mutants (D, D’), and *ola226* mutants (E, E’). Brackets indicate the AIY Zone 1 region, and asterisks mark the AIY Zone 2 region (see Figure 1A). The animals imaged in B-E are not the same as B’-E’. **(F-H)** Quantification of phenotypes, including the length of glia anterior process (F, indicated in schematic A), the length of ventral endfeet (G, indicated in schematic A) and the percentage of animals displaying overlap between the AIY synapses and the VCSC glia in Zone 1 (H). The total number of animals (N) and the number of times scored (n) are indicated in each bar for each genotype as N/n. Statistical analyses are based on two-tailed student’s t-test. Error bars represent SEM, NS: not significant as compared to wild type, ****p< 0.0001 as compared to wild type (if on top of bar graph), unless brackets are used between two compared genotypes.

Fluorescent intensity of MIG-17::mNeonGreen and EMB-9::mCherry was quantified with Image J from confocal images at the specified developmental stages.

### Statistical analysis

Specified statistical analyses were based on student’s T-test and performed with Prism 6. For comparisons of mean fluorescence intensities, ectopic synapse ratio, length or length ratio, we used an unpaired two-tailed Student’s t test.

## RESULTS

### *ola226* suppresses defects of *cima-1 (wy84)* synaptic allometry

The AIY interneurons are a pair of bilaterally symmetric neurons in the *C. elegans* brain (nerve ring). AIYs display a stereotyped and specific pattern of presynaptic specializations (White et al., 1986;Colon-Ramos et al., 2007). This pattern is established during embryogenesis and, although the animals grow an order of magnitude in length from early embryogenesis to adulthood (from ∼100 μm to ∼1mm) (Knight et al., 2002;Shibata et al., 2016), the AIY synaptic pattern is maintained during growth (Figure 1A-1B, 1E,-1E’” and (Shao et al., 2013)). We term this process of correct maintenance of synaptic positions during growth “synaptic allometry”.

From forward genetic screens we had identified *cima-1,* a gene required for synaptic allometry (Shao et al., 2013). In *cima-1* mutants, the AIY synaptic pattern develops correctly. However, as animals grow, ectopic synapses emerge in the Zone 1 region, a region of the AIY neuron which is normally asynaptic (Figure 1C, 1F-1F”’ and (Shao et al., 2013)). *cima-1* encodes a solute carrier transporter which does not function in neurons, but is rather required in epidermal cells to antagonize the FGF receptor and likely modulate epidermal-glia adhesion ((Shao et al., 2013) and Figure S1 for model). In *cima-1* mutants, crosstalk between the epidermal cell and the neighboring ventral cephalic sheath cell glia (VCSC glia) is affected, resulting in defects in VCSC glia position during growth. Abnormal VCSC glia ectopically ensheath the normally asynaptic Zone 1 region of AIY, resulting in ectopic synapses in Zone 1 (Figure S1). To identify molecules which cooperate with *cima-1* in regulating synaptic allometry, we performed an unbiased EMS screen in *cima-1(wy84)* mutants for suppressors of ectopic synapses, and isolated allele *ola226*.

Although the animal morphology and the guidance of AIY neurites are largely unaffected in *cima-1(wy84);ola226* double mutants (Figure S2 and data not shown), the newly isolated *ola226* allele robustly suppresses the ectopic presynaptic structures of AIY Zone 1 observed in *cima-1(wy84)* mutants (93.9% of animals displayed ectopic synapses in *cima-1(wy84)* vs 54.6% in *cima-1(wy84);ola226* double mutants, p<0.0001, Figure 1D, 1G-1I). Suppression was observed both for the vesicular marker RAB-3 and for the active zone marker SYD-1 (Figure 1E’, 1F’, 1G’), suggesting that the *ola226* allele suppresses ectopic assembly of presynaptic structures, and not just relocalization of synaptic vesicles. Moreover, young *cima-1(wy84);ola226* animals display a wild type pattern of presynaptic specializations (Figure 1D), suggesting that the *ola226* allele does not generally affect synaptogenesis. Instead, these results suggest that *ola226* is specifically required for the suppression of ectopic presynaptic specializations resulting from *cima-1* induced defects in synaptic allometry.

Synaptic allometry is regulated by growth, and the penetrance of the *cima-1* phenotype is affected by the size of the animal (Shao et al., 2013). For example, dumpy (*dpy*) mutants, which are about 25% shorter than wild type animals, suppress synaptic allometry defects in *cima-1* mutants (Figure S2A, S2B, S2D-S2E, S2D’-S2E’ and (Shao et al., 2013)). Conversely, long (*lon*) mutants, which are up to 30% longer than wild type (Brenner, 1974;Morita et al., 2002;Nystrom et al., 2002;Suzuki et al., 2002), enhance the *cima-1* mutant phenotype (Figure S2C, S2F-S2F’ and (Shao et al., 2013)). To determine if *ola226* affects synaptic allometry by regulating body size, we examined animal size in *ola226* single or *cima-1(wy84);ola226* double mutants. We determined that the length of either *ola226* single or *cima-1(wy84);ola226* double mutants is similar to that of wild type or *cima-1(wy84)* mutant animals (Figure S2I). Therefore, the suppression of the AIY ectopic synaptic positions by *ola226* (Figure S2G-S2H’) is not due to an effect of *ola226* on the size of the animal.

Together, our findings indicate that *ola226* represents a genetic lesion required for the emergence of ectopic presynaptic sites during growth in *cima-1* mutants.

### Glia morphology is affected in *ola226* mutants

*Cima-1* affects synaptic allometry by repositioning ventral cephalic sheath cell (VCSC) glia during growth (Shao et al., 2013). To test if *ola226* also affects VCSC glia position, we labeled the VCSC glia with mCherry in wild type and indicated mutants, and quantified VCSC glia position and morphology (Figure 2A). Consistent and extending our previous observations, we determined that the VCSC glia defects in *cima-1(wy84)* mutants result from defects in both position and morphology of the VCSC glia during growth. As *cima-1* mutant animals grow, VCSC glia cell bodies are posteriorly displaced, resulting in longer VCSC glia anterior processes (length of the VCSC glia anterior process: 113.35μm in wild type, 127.53μm in *cima-1(wy84)* mutants, p<0.0001. Figure 2B, 2C, 2F). VCSC glia morphology is also altered in *cima-1* mutants, with endfeet abnormally extending posteriorly (length of VCSC glia endfeet: 45.52μm in wild type and 51.47μm in *cima-1(wy84)* mutants, p<0.0001. Figure 2B-2C, 2G). These two defects change the relative positions of VCSC glia and the AIY neurite, resulting in ectopic contact of the VCSC glia with the asynaptic Zone 1 region, and concomitant emergence of ectopic presynaptic sites in Zone 1 (Figure 2B’-C’, 2H). Ablation of VCSC glia suppress the ectopic synaptic phenotype in Zone 1 (Shao et al, 2013), indicating the importance of glia in the emergence of these ectopic synapses.

In *cima-1(wy84);ola226* double mutants, VCSC glia cell body position and endfeet morphology phenotypes are suppressed (length of glia anterior process: 127.53μm in *cima-1(wy84)* and 120.68μm in *cima-1(wy84);ola226*, p<0.0001; length of VCSC glia endfeet: 51.47μm in *cima-1(wy84)* and 45.19μm in *cima-17(wy84);ola226*, p<0.0001. Figure 2C, 2D, 2F, 2G). This suppression, in turn, results in *cima-1(wy84);ola226* double mutants having a reduced region of contact between the AIY neurons and VCSC glia (88.70% in *cima-1(wy84)* and 33.67% in *cima-1(wy84);ola226*, p<0.0001. Figure 2H). Consequently, ectopic presynaptic specializations in AIY Zone 1 are suppressed (Figure 2D’). Our findings suggest that *ola226* is a genetic lesion that suppresses *cima-1* ectopic synapses by reverting the *cima-1* phenotypes on glia position and morphology.

To better understand the phenotype from allele *ola226*, we outcrossed *cima-1* and examined the VCSC glia and the AIY synaptic phenotypes for just the *ola226* allele. We determined that animals carrying the *ola226* allele do not display defects in the position of the VCSC glia (length of glia anterior process: 113.35μm in wild type and 113.68μm in *ola226*, p=0.72. Figures 2E and 2F). However, *ola226* mutants do display a modest but significant defect in the VCSC glia morphology, with posterior end-feet being shorter in *ola226* as compared to wild type animals (length of glia end-feet: 45.52μm in wild type, 39.79μm in *ola226* p<0.0001. Figure 2G). *ola226* mutants also display a concomitant defect in the position of AIY, with both the neurite and the soma being anteriorly displaced as compared to wild type animals (Figure S3). Interestingly, while *ola226* mutants have phenotypes for both glia morphology and AIY neurite position, the area of overlap between the glia and AIY is not affected, nor is the distribution of presynaptic specializations as compared to wild type (Figure 2E’ and 2H). These phenotypes demonstrate that it is not just glia morphology, glia position or even the position of the AIY neurite in the animal that regulates synaptic allometry, but rather the relative position between the VCSC glia and the AIY neurons which determines synaptic positions during growth. Our findings indicate that both *cima-1* and our newly identified *ola226* allele affect relative positions of glia and the AIY interneurons during post-embryonic growth, and therefore, affect synaptic allometry by altering the areas of overlap between these two cells. Together our data indicate that allele *ola226* is required for normal VCSC glial morphology, and that it affects synapses by acting in opposition to *cima-1*, thereby reverting the defective interaction between glia and AIY interneurons observed for *cima-1* mutants.

### *ola226* is a lesion in *mig-17,* a gene that encodes an ADAMTS metalloprotease

To identify which gene is affected in the *ola226* allele, we performed SNP mapping, whole genome sequencing and transgenic rescue experiments. The *ola226* allele results from a G to A mutation at the end of first exon of the *mig-17* gene, and alters a conserved glutamic acid residue at position 19 to a lysine (Figure 3A). To then test if *ola226* is a loss-of-function allele of *mig-17*, we examined two additional loss-of-function *mig-17* alleles, *mig-17(k113)* and *mig-17(k174)* (Nishiwaki, 1999;Nishiwaki et al., 2000). *mig-17(k113)* is a point mutation in the first intron of the gene and is predicted to affect correct splicing, while *mig-17(k174)* allele results from a change in Q111 to a premature stop codon, resulting a putative null allele (Figure 3A) (Shibata et al., 2016). We found that both *mig-17(k113)* and *mig-17(k174)* mutants suppress the ectopic synapses in *cima-1(wy84)* mutants (91.9% of animals display ectopic synapses in *cima-1(wy84)*, 62.3% in *cima-1(wy84);mig-17(k113)*, 29.9% in *cima-1(wy84);mig-17(k174)* and 45.7% in *cima-1(wy84);mig-17(ola226)*, p<0.0001 for all double mutants as compared to *cima-1(wy84)*; Figures 3B-3F, 3I). Importantly, introducing a wild type copy of *mig-17* genomic sequence results in robust rescue of the *ola226* phenotype in the *cima-1(wy84);mig-17(ola226)* double mutants (45.70% of animals display ectopic synapses in *cima-1(wy84); mig-17(ola226)* and 78.04% in *cima-1(wy84);mig-17(ola226);Pmig-17::mig-17*(genomic), p<0.0001; Figures 3G and 3I). Together our findings indicate that *ola226* is a recessive loss-of-function allele of *mig-17* which suppresses *cima-1(wy84)* defects in synaptic allometry.

**Figure 3.**
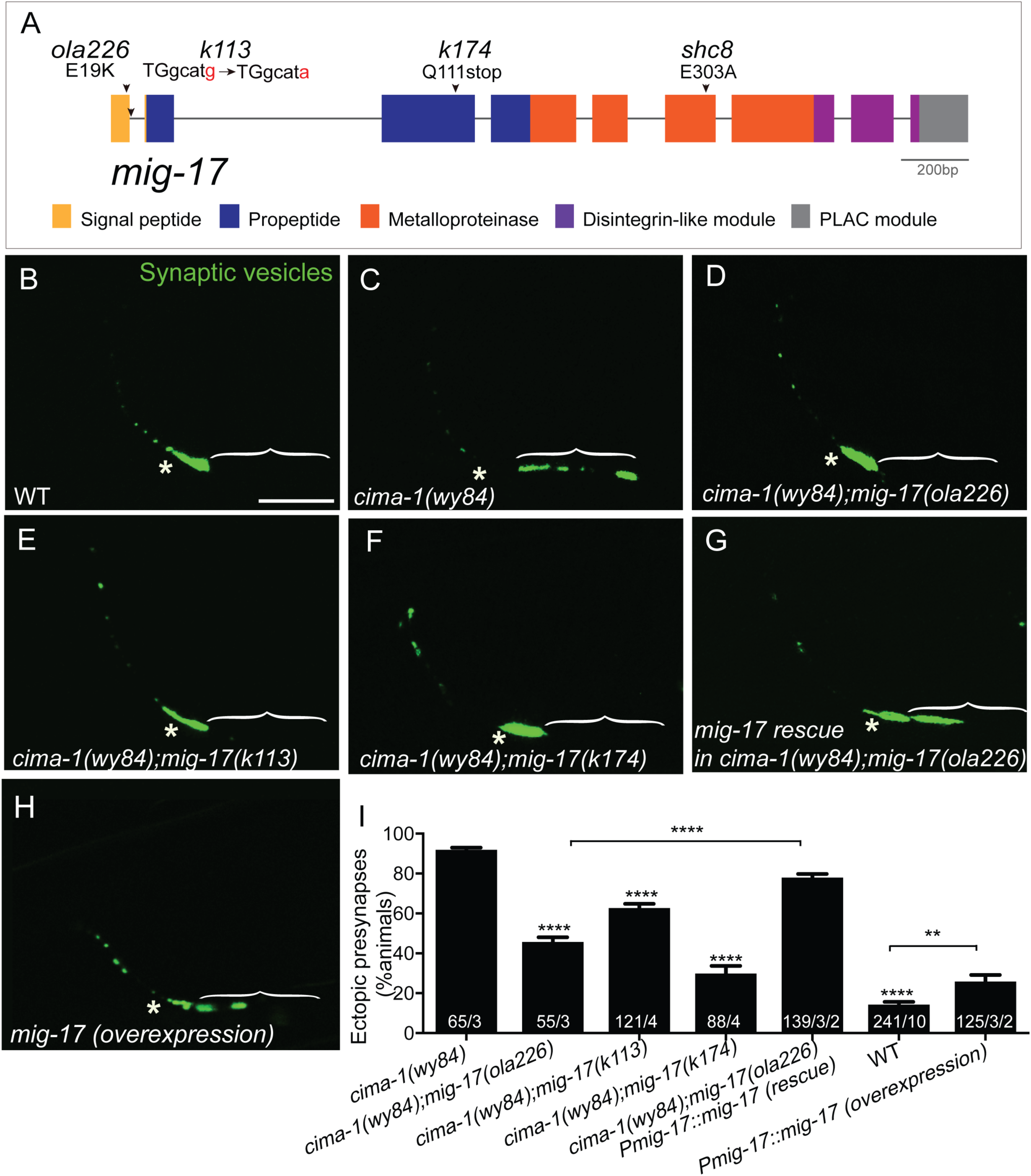
*ola226* is a lesion in the *mig-17* gene. **(A)** Schematic diagram of the *mig-17* gene, corresponding protein domains coded by the exons (colored) and genetic lesions for the alleles used in this study. **(B-H)** Confocal micrographs of the AIY synaptic vesicle marker GFP::RAB-3 (green) in adult wild type (B), *cima-1(wy84)* (C), *cima-1(wy84);mig-17(ola226)* (D), *cima-1(wy84);mig-17(k113)* (E), *cima-1(wy84);mig-17(k174)* (F), *cima-1(wy84);mig-17(ola226)* animals expressing a wild type copy of the *mig-17* gene (P*mig-17::mig-17(genomic)*) (G), and wild type animals over-expressing the *mig-17* gene (P*mig-17::mig-17(genomic)*) (H). Brackets indicate the AIY Zone 1 region; asterisks indicate the Zone 2 region. Scale bar in B applies to all images, 10μm. **(I)** Quantification of the percentage of animals with ectopic synapses in the AIY Zone 1 region for indicated genotypes. The total number of animals (N), the number of times scored (n1) are indicated in each bar for each genotype, as are, for the transgenic lines created, the number of transgenic lines (n2) examined (all using the convention N/n1/n2). Statistical analyses are based on two-tailed student’s t-test. Error bars represent SEM, **p<0.01, ****p< 0.0001 as compared to *cima-1 (wy84)* (if on top of bar graph), unless brackets are used between two compared genotypes.

To further explore how *mig-17* regulates synaptic distribution, we examined the AIY synaptic phenotype in the loss-of-function mutants *mig-17(ola226)* and *mig-17(k113)*, and also over-expressed a genomic construct of *mig-17* in wild type animals (*mig-17(OE)*). We found that *mig-17(k113)* loss-of-function allele displays a normal synaptic distribution in AIY, similar to what we had observed for the *mig-17(ola226)* allele (Figures S4A-S4D). We also observed that over-expressing *mig-17* in wild type animals (*mig-17(OE)*) resulted in ectopic synapses in the Zone 1 region of AIY (Figure 3H and 3I). These data are consistent with our studies using the *mig-17(ola226)* allele, which demonstrate that *mig-17* does not affect AIY synaptic distribution on its own because it does not alter the AIY:VCSC relationship (Figure 2). The data also demonstrate that overexpression of *mig-17* phenocopies *cima-1(wy84)* loss-of-function allele, in support of our model that the *mig-17* acts in opposition to *cima-1*.

MIG-17 is best known for its post-embryonic roles in regulating distal tip cell migration during gonad development (Nishiwaki, 1999), and pharyngeal size and shape during growth (Shibata et al., 2016). Since AIY is present near the *C. elegans* pharynx, we tested if the synaptic positions of AIY are related to pharyngeal length. We found that the length of the pharynx slightly increases in *cima-1(wy84)* mutants as compared to wild type, and that the increase in pharynx length is more robust for *mig-17(ola226)* mutants (pharynx length is 146.6μm in wild type, 150.6μm in *cima-1(wy84)* and 157.9μm in *mig-17(ola226)* animals, p<0.0001 between wild type and *cima-1(wy84)* and *mig-17(ola226)* mutant animals; Figure S5). We also observed the *cima-1(wy84);mig-17(ola226)* double mutants enhance the *cima-1* pharynx length phenotype (pharynx length is 160.8μm for *cima-1(wy84);mig-17(ola226)* double mutants, p<0.0001 when compared with *cima-1(wy84)* mutants; Figure S5). Therefore, while *mig-17* acts in opposition to *cima-1* in synaptic allometry, it enhances *cima-1* for the pharyngeal length phenotype. Our findings suggest that the synaptic allometry phenotypes do not simply result from a defect in pharynx length. Our studies are consistent with previous reports on the role MIG-17 in pharyngeal length regulation (Shibata et al., 2016) and extend them, now indicating that MIG-17 is also required for glia morphology and modulation of synaptic allometry during growth.

### MIG-17 and EGL-15/FGFR work in the same pathway to promote the formation of ectopic synapses in *cima-1(wy84)*

Our previous study showed that CIMA-1 negatively regulates the Fibroblast Growth Factor Receptor (FGFR) EGL-15(isoform 5A) in the epidermal cells to position glia and synapses during growth. Consistent with this interpretation, mutation of *egl-15(5A)* suppressed the ectopic synapses caused by *cima-1(wy84)*, and overexpression of the EGL-15(5A) ectodomain in wild type animals phenocopied *cima-1* mutants (Shao et al., 2013). These genetic findings are also consistent with western-blot data demonstrating regulation of EGL-15(5A) levels by CIMA-1 (Shao et al, 2013). Together, the data support a model whereby *cima-1* modulates epidermal-glia cell adhesion via regulation of EGL-15/FGFR ectodomain which acts, not in its canonical signaling role, but as an extracellular adhesion factor (Bulow et al., 2004;Shao et al., 2013).

To understand how *mig-*17 cross talks with this pathway in the regulation of synaptic allometry, we examined its genetic relationship to EGL-15/FGFR. We generated *cima-1(wy84);egl-15(n484)* double mutants and *cima-1(wy84);mig-17(ola226);egl-15(n484)* triple mutants and observed AIY synaptic distribution. We determined that *cima-1(wy84);egl-15(n484)* double mutants, *cima-1(wy84);mig-17(ola226)* double mutants and *cima-1(wy84);mig-17(ola226);egl-15(n484)* triple mutants similarly suppress *cima-1(wy84)* phenotypes (Figures 4A-4F). We note that while the observed suppression is not a complete reversion to wild type phenotypes, it is consistent with the degree of suppression observed for glia-ablated animals (Shao et al., 2013). Our findings might indicate a ceiling effect at the level of the contribution of glia to the synaptic phenotypes, and suggest that other glia-independent mechanisms also contribute to synaptic allometry through molecular pathways distinct from those regulated by MIG-17 and EGL-15. Importantly, these data suggest that *mig-17* and *egl-15* genetically act in the same pathway to promote the formation of ectopic synapses in *cima-1(wy84)* mutants via regulation of glia morphology and position.

**Figure 4.**
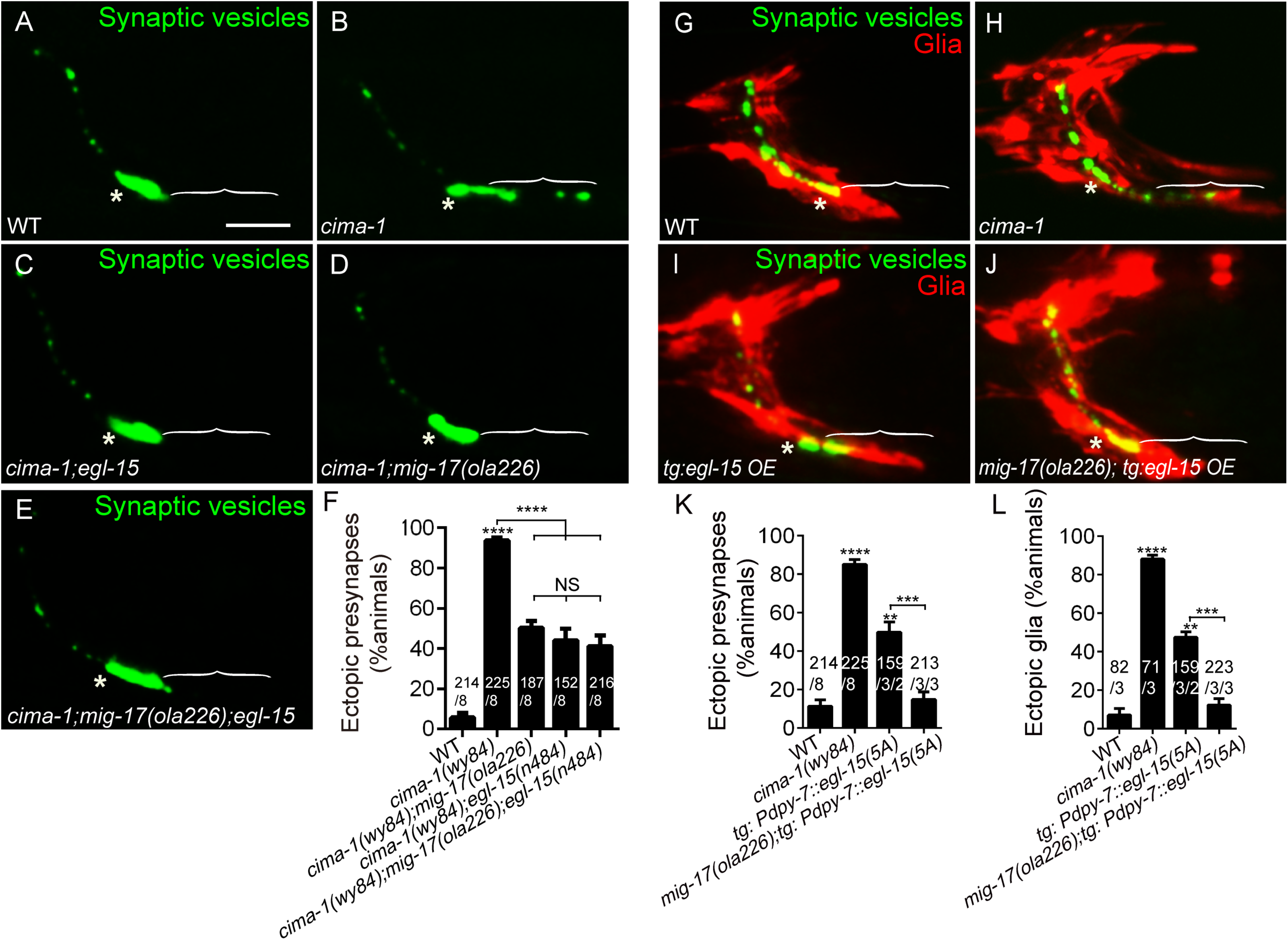
MIG-17 genetically interacts with EGL-15/Fibroblast Growth Factor Receptor to regulate synaptic allometry. **(A-E)** Confocal micrographs of the AIY synaptic vesicle marker GFP::RAB-3 (green) in adult wild type (A), *cima-1(wy84)* (B), *cima-1(wy84);egl-15(n484)* (C), *cima-1(wy84);mig-17(ola226)* (D), *cima-1(wy84);mig-17(ola226);egl-15(n484)* (E) adult animals. In all images (A-E, G-J), brackets indicate the AIY Zone 1 region, asterisks mark the Zone 2 region and scale bar (in (A)), 10μm. **(F)** Quantification of the percentage of animals with ectopic synapses in the AIY Zone 1 region for indicated genotypes. **(G-J)** Confocal micrographs of AIY synaptic vesicle marker GFP::RAB-3 (green) and VCSC glia (red) in adult wild type (G), *cima-1(wy84)* (H), wild type overexpressing EGL-15(isoform 5A) in epidermal cells by using P*dpy-7::egl-15(5A)* (I) and *mig-17(ola226)* overexpressing EGL-15(isoform 5A) in epidermal cells by using P*dpy-7::egl-15(5A)* (J). **(K-L)** Quantification of the percentage of animals with ectopic synapses in the AIY Zone 1 (K) and with distended glia endfeet (L) for indicated genotypes. In the graphs, the total number of animals (N), the number of times scored (n1) are indicated in each bar for each genotype, as are, for the transgenic lines created, the number of transgenic lines (n2) examined (all using the convention N/n1/n2). Statistical analyses are based on two-tailed student’s t-test. Error bars represent SEM, NS: not significant, **p<0.01, ***p<0.001, ****p < 0.0001 as compared to wild type (if on top of bar graph), unless brackets are used between two compared genotypes.

EGL-15(isoform 5A) is required for maintenance of axon positions during movement and growth (Bulow et al., 2004). The overexpression of EGL-15(5A) in epidermal cells also promotes VCSC glia end-feet extension and ectopic synapses in AIY. This result phenocopies *cima-1(wy84)* mutants and is consistent with *cima-1* acting antagonistically to the EGL-15/FGF Receptor (Figures 4G-4I and (Shao et. al. 2013)). To further probe the relationship between EGL-15(5A) and MIG-17, we examined synapses and glia in animals overexpressing EGL-15(5A). Interestingly, and consistent with MIG-17 and EGL-15 acting in the same synaptic allometry pathway, we observed that *mig-17(ola226)* suppresses the VCSC glia extension and ectopic synapses in animals over-expressing EGL-15(5A) (Figure 4I-4L). Together, our genetic findings indicate that EGL-15(5A) and MIG-17 act in the same pathway to position glia and regulate synaptic allometry during growth. Our findings also indicate that MIG-17 is epistatic to EGL-15/FGFR in positioning glia and regulating synaptic allometry, and that the defects observed for EGL-15/FGFR overexpressing-animals require *mig-17*.

### MIG-17 is expressed by muscles to regulate synaptic allometry

CIMA-1 and FGFR EGL-15(5A) are expressed by epidermal cells to position glia and synapses during growth (model in Figure S1 and (Shao et. al. 2013)). To determine where MIG-17 acts, we first analyzed the expression pattern of *mig-17*. We found that a *mig-17* transcriptional GFP reporter is robustly expressed by body wall muscles as colabeled by P*myo-3::mCherry* and consistent with previous reports (Nishiwaki et al., 2000) (Figures 5A-5A’’’). Additionally, we observed that in the head region, the reporter is seen in the nervous system (Figure 5B-5B’’’). We did not detect expression of MIG-17 in VCSC glial cells, or in epidermal cells where the MIG-17 genetic interactors CIMA-1 and FGFR EGL-15(5A) are expressed (Figure 5C-5D’’’).

**Figure 5.**
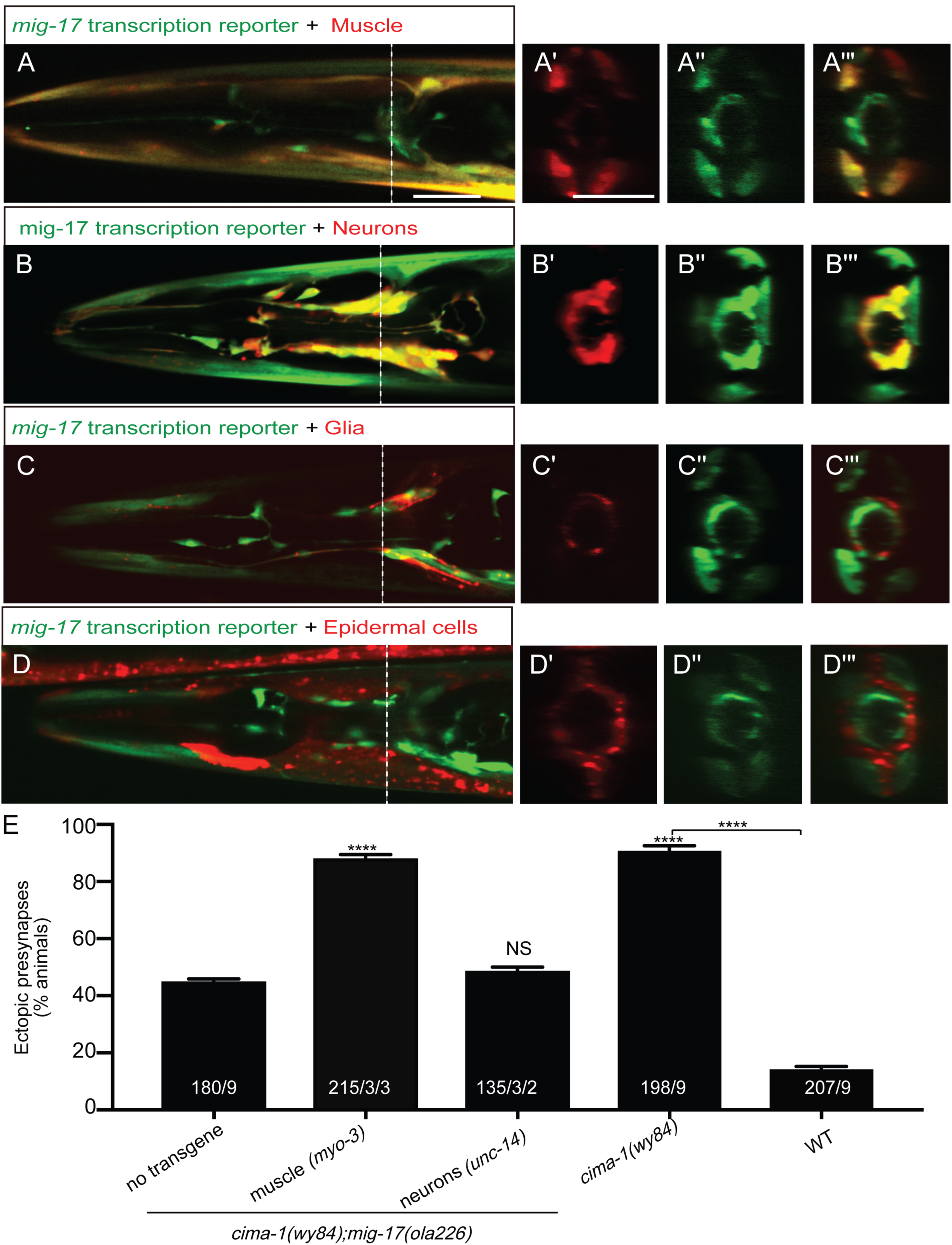
MIG-17 is expressed by body wall muscles to regulate synaptic allometry. **(A-D”’)** Confocal micrographs of adult animals expressing the transcriptional reporter *mig-17(genomic)::SL2::GFP* (green) with reporters that co-label body wall muscles (P*myo-3::mCherry* (A-A”’)), epidermal cells (P*dpy-4::mCherry* (B-B”’)), VCSC glia (P*hlh-17::mCherry* (C-C”’)) and neurons (P*rab-3::mCherry* (D-D”’). Images (A’-D”’) correspond to a transverse cross-section of the confocal micrographs, specifically for the region corresponding to the dashed white line in (A-D). The scale bar (10μm) in A applies to B, C, D, and in A’ applies all transverse cross-section images. **(E)** Quantification of the percentage of adult animals with ectopic synapses in the AIY Zone 1 region of the indicated genotypes and rescue experiments. The total number of animals (N) and the number of times scored (n1) are indicated in each bar for each genotype, as are, for the transgenic lines created, the number of transgenic lines (n2) examined (all using the convention N/n1/n2). Statistical analyses are based on two-tailed student’s t-test. Error bars represent SEM, NS: not significant, ****p< 0.0001 as compared to the no-transgene control (if on top of bar graph), unless brackets are used between two compared genotypes.

To determine the *mig-17* site of action, we expressed *mig-17* in the two tissues in which we observed *mig-17* expression: the nervous system (using the *unc-14* promoter) and the body wall muscles (using the *myo-3* promoter). We found robust rescue of *cima-1(wy84);mig-17(ola226)* phenotype when *mig-17* was expressed in body wall muscles, but not upon expression in the nervous system (Figure 5E). Together, our findings indicate that MIG-17 is expressed by muscle cells to modulate synaptic allometry. Our findings also indicate that muscle-derived MIG-17 acts in the same pathway as epidermally-expressed CIMA-1 and FGFR EGL-15(5A) to modulate glia morphology and synaptic allometry. Our findings suggest that multiple non-neuronal tissues act *in vivo* to convey growth information to the nervous system, and regulate synaptic allometry through glia position.

### MIG-17 localizes to basement membrane

To better understand how MIG-17, expressed by muscles, cooperates with CIMA-1, expressed by epidermal cells, to regulate synaptic positions in neurons, we generated a MIG-17::mNeonGreen knock-in allele (via CRISPR-Cas9 strategies) that allows us to visualize endogenous MIG-17 localization throughout development (Dickinson et al., 2013)(Figure S6A). Using this MIG-17::mNeonGreen reporter, we observed that in the head-region, MIG-17 prominently localizes to the extracellular matrix proximal to the pharynx bulb. Furthermore, MIG-17 protein levels are detectable in larva stage 1 through larva stage 4, but becomes undetectable upon reaching the adult stage (Figure 6A-6H). Our findings are consistent with previous studies indicating that MIG-17 accumulates in the extracellular matrix (Ihara and Nishiwaki, 2007;Shibata et al., 2016) and *in situ* and western blot studies demonstrating accumulation of the active form of MIG-17 during larva stage 3 (L3) and larva stage 4 (L4), and downregulation in adults (Ihara and Nishiwaki, 2008).

**Figure 6.**
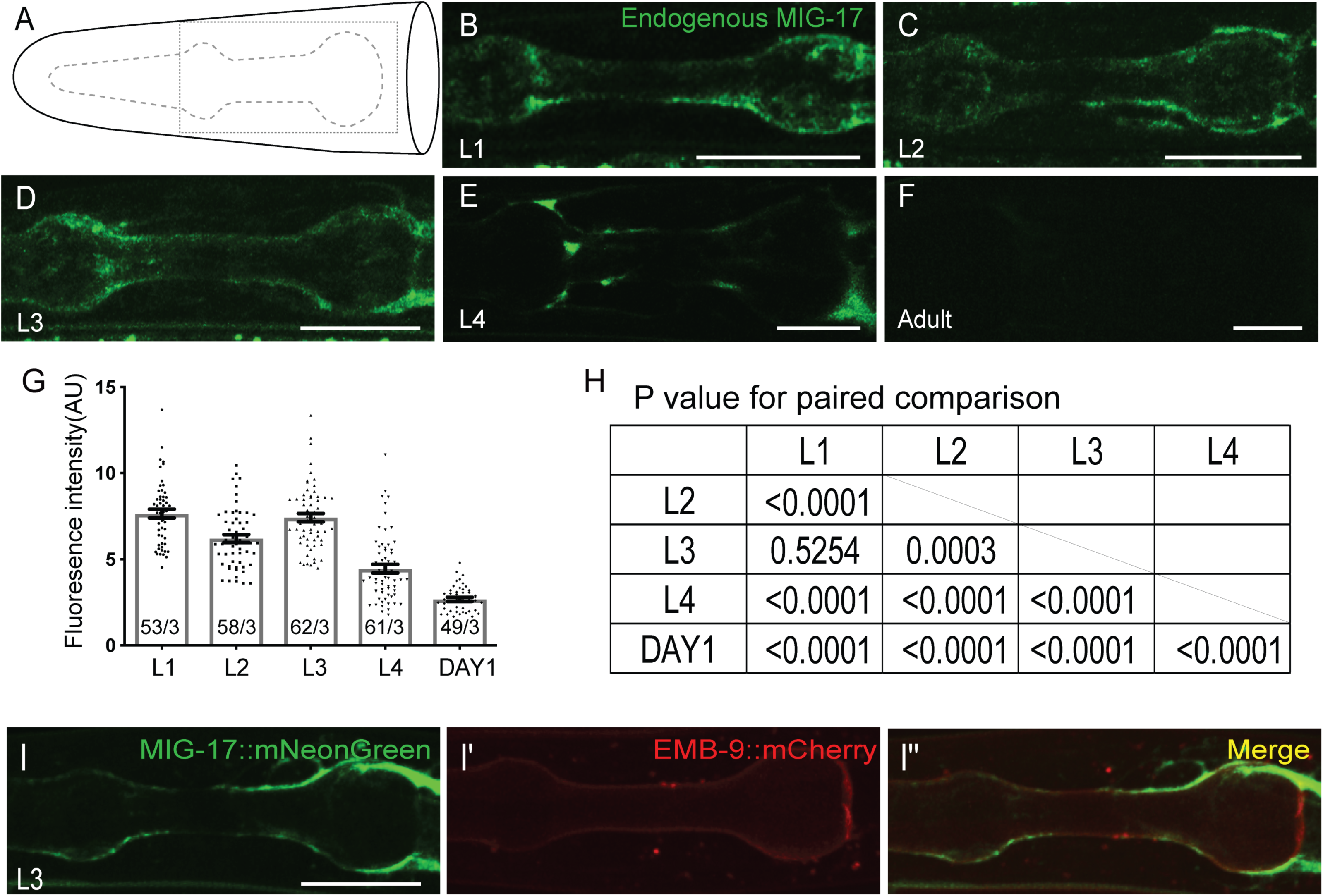
MIG-17 is developmentally regulated and localizes to the basement membrane. **(A)** Cartoon diagram of the head of *C. elegans*, similar to Figure 1A. The dotted box indicates the region imaged in the subsequent micrographs. **(B-F)** Confocal micrographs of animals with a CRISPR-engineered MIG-17::mNeonGreen, imaged at larva stage 1 (L1 in (B)), larva stage 2 (L2 in (C)), larva stage 3 (L3 in (D)), larva stage 4 (L4 in (E)) and 1 day-old adults (F). **(G-H)** Quantification of MIG-17::mNeonGreen intensity at different developmental stages (G) and the p-value for paired comparison based on two-tailed student’s t-test (H). In the graph, the total number of animals (N) and the number of times scored (n) are indicated in each bar for each genotype as N/n. Statistical analyses are based on two-tailed student’s t-test. Error bars represent SEM. **(I-I’’)** Confocal micrographs of adult animals expressing MIG-17::mNeonGreen (I) and EMB-9::mCherry (I’) and merged image of both markers. For all images, scale bars are 10μm.

The localization of MIG-17 is reminiscent of that reported for basement membrane proteins (Ihara et al., 2011). To more carefully determine if the localization of secreted MIG-17 corresponds to the basement membrane, we simultaneously examined MIG-17 localization with the basement membrane protein EMB-9::mCherry (Kramer, 2005). We observed that MIG-17 colocalizes with EMB-9 at the extracellular space (Figure 6I-6I”). Together, our findings indicate that MIG-17 is a muscle-derived secreted metalloprotease whose levels are regulated during development, and that it localizes to the basement membrane to modulate synaptic allometry during growth.

### The metalloprotease activity of MIG-17 is required to promote the formation of ectopic synapses in *cima-1(wy84)* mutant animals

MIG-17 is best known for its role in distal tip cell migration in *C. elegans* (Nishiwaki, 1999;Jafari et al., 2010;Shibata et al., 2016). This role depends on the remodeling of gonadal basement membrane through MIG-17 metalloprotease enzymatic activity (Nishiwaki et al., 2000). To determine if MIG-17 metalloprotease enzymatic activity is also required for promoting the formation of ectopic synapses in *cima-1(wy84)*, we engineered an E303A point mutation at the metalloprotease catalytic site via CRISPR/cas-9 to generate the *mig-17(shc8)* allele (Figures 7A and S6B) (Nishiwaki et al., 2000;Dickinson et al., 2013). We observed that our engineered *mig-17(shc8)* allele behaves like other loss-of-function alleles of *mig-17*, suppressing the ectopic synapses of *cima-1(wy84)* mutant animals (91.91% of animals displayed ectopic synapses in *cima-1(wy84)* vs 57.49% in *cima-1(wy84); mig-17(shc8)*, p<0.0001, Figures 7B-7E, 7H). Consistent with this result, we also determined that a transgene with the E303A (*mig-17(E303A)*) lesion is incapable of rescuing the *mig-17* suppression in *mig-17(ola226);cima-1(wy84)* mutants (78.04% of animals displaying ectopic synapses in *cima-1(wy84);mig-17(ola226)* animals rescued with the wild type genomic fragment of *mig-17 (tg:Pmig-17::mig-17(genomic),* vs 45.78% in *cima-1(wy84);mig-17(ola226)* with the genomic fragment with the point mutation in the protease active site (*tg:Pmig-17::mig-17(E303A),* p<0.0001; Figures 7F-7H). Our data are consistent with structure-function studies of MIG-17 which have underscored the importance of its metalloprotease domain (Nishiwaki et al., 2000), and indicate that MIG-17 protease activity is also required for promoting AIY ectopic synapse formation in *cima-1(wy84)* mutants during growth.

**Figure 7.**
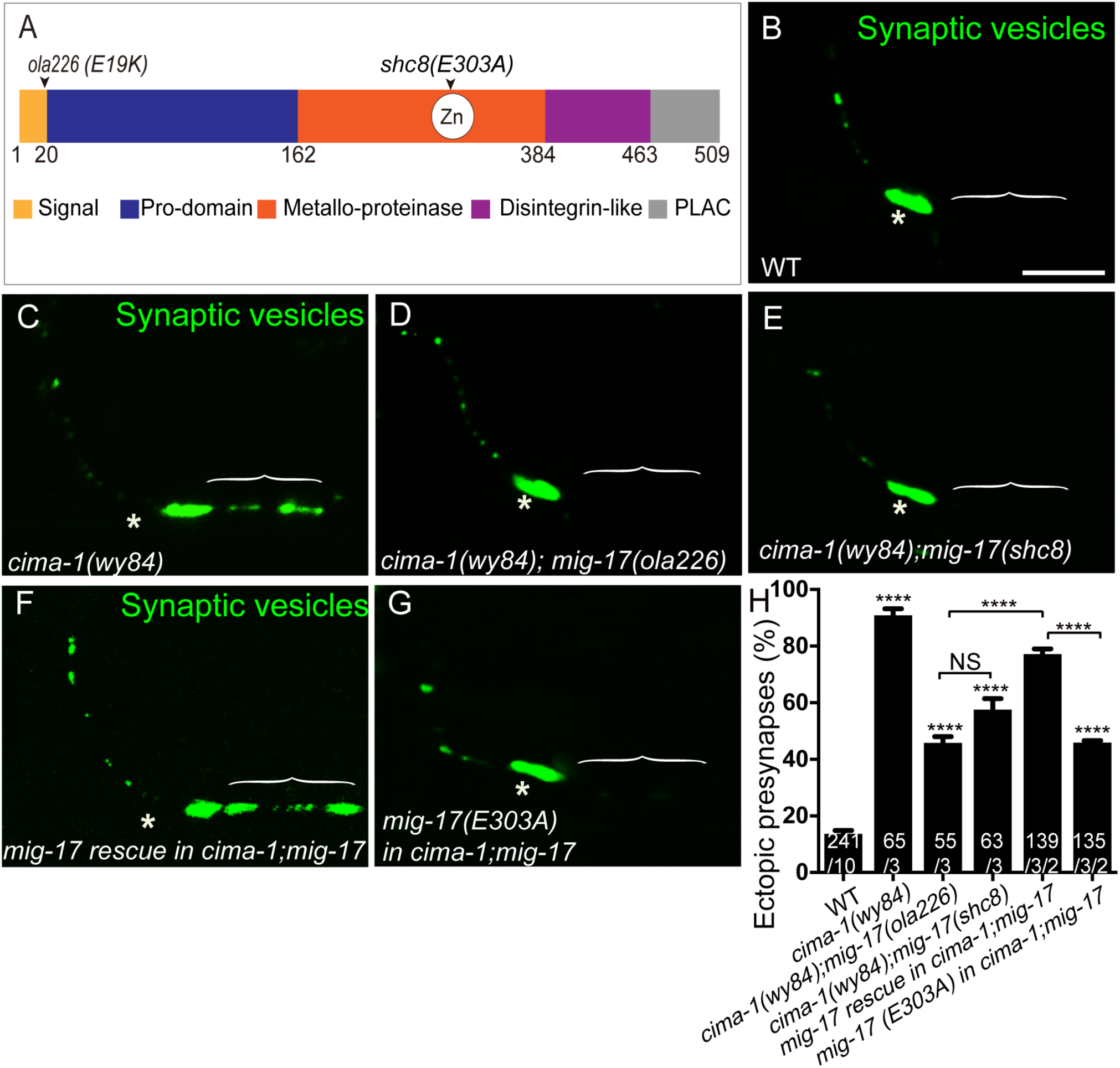
The metalloprotease activity of MIG-17 is required to suppress the formation of ectopic synapses in *cima-1(wy84)* mutant animals. **(A)** Schematic diagram of the MIG-17 protein, corresponding conserved protein domains (colored) and genetic lesions for the alleles used in this study. **(B-G)** Confocal micrographs of the AIY presynaptic sites labeled with the synaptic vesicle marker GFP::RAB-3 (pseudo-colored green) in adult wild type (B), *cima-1(wy84)* (C), *cima-1(wy84);mig-17(ola226)* (D), *cima-1(wy84);mig-17(shc8)* (E), *cima-1(wy84);mig-17(ola226)* animals expressing a wild type copy of the *mig-17* genomic DNA (P*mig-17::mig-17)* (F), and *cima-1(wy84);mig-17(ola226)* animals expressing a copy of the *mig-17* genomic DNA with a point mutation in the metalloprotease domain (P*mig-17::mig-17(E303A)*) (G). Brackets indicate the AIY Zone 1 region; asterisks indicate the Zone 2 region. The scale bar in B is 10μm and applies to all images. **(H)** Quantification of the percentage of animals with ectopic synapses in the AIY Zone 1 region in indicated genotypes. In the graph, the transgene rescue with wild type copy of the *mig-17* genomic DNA control data is the same as in Figure 3I. The total number of animals (N) and the number of times scored (n1) are indicated in each bar for each genotype, as are, for the transgenic lines created, the number of transgenic lines (n2) examined (all using the convention N/n1/n2). Statistical analyses are based on two-tailed student’s t-test. Error bars represent SEM, NS: not significant, ****p< 0.0001 as compared to wild type (if on top of bar graph), unless brackets are used between two compared genotypes.

### MIG-17 regulates basement membrane proteins to modulate synaptic allometry

To determine how the MIG-17 metalloprotease activity might regulate synaptic allometry, we first examined the proteome through liquid chromatography–tandem mass spectrometry (LC-MS/MS) analyses in wild type and *mig-17(ola226)* mutant animals. Unbiased comparative LC-MS/MS analyses had not been performed for *mig-17(ola226)* mutant animals, and our analyses provided an opportunity to both identify new targets for the MIG-17 metalloprotease, and to characterize the proteome of the basement membrane in the *mig-17* mutant background.

We observed significant and reproducible differences in the proteome of *mig-17(ola226)* mutants as compared to wild type animals (Table S3). Consistent with the importance of MIG-17 in the remodeling of the basement membrane (Kim and Nishiwaki, 2015), we observed significant differences in the protein levels of basement membrane components for the *mig-17(ola226)* mutants as compared to wild type. Importantly, a number of basement membrane components displayed increased protein levels in the *mig-17(ola226)* mutants, including EMB-9/Collagen IV α1 chain, LET-2/Collagen IV α2 chain, OST-1/Sparc, UNC-52/Perlecan, NID-1/nidogen, EPI-1/laminin-α, LAM-1/laminin-β, and LAM-2/laminin-γ, (Figure 8A and Table S3). Together, our proteomic analyses reveal potential targets of MIG-17, and extends our understanding of the role for this ADAMTS metalloprotease in regulating proteins in the basement membrane.

**Figure 8.**
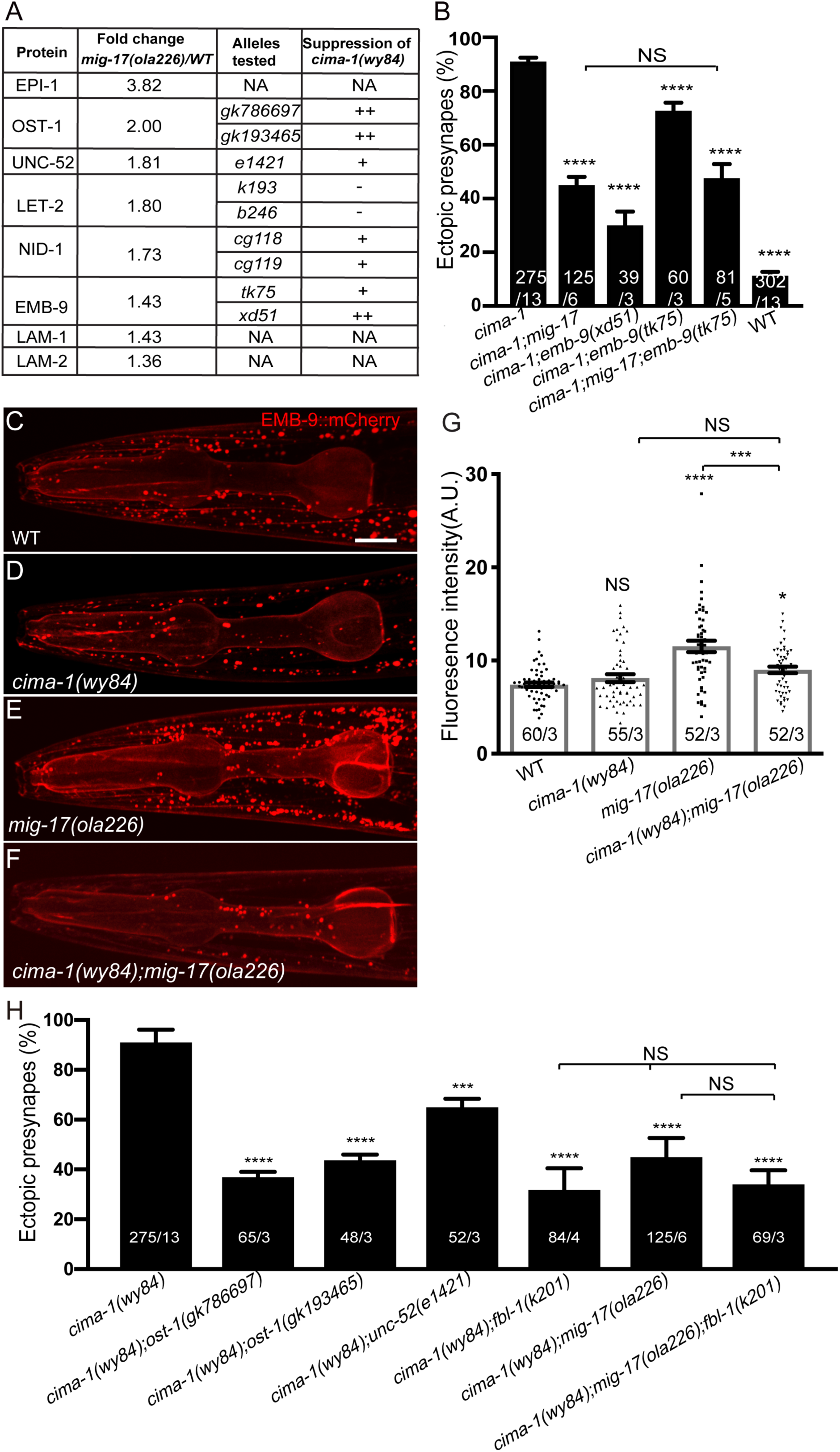
MIG-17 suppresses the synaptic allometry defect in *cima-1(wy84)* mutants through modulation of basement membrane proteins. **(A)** List of basement membrane components upregulated in the mass spectrometry analyses (see also Table S3), and alleles tested with *cima-1* for their capacity to suppress the synaptic allometry phenotypes in adult worms. **(B)** Quantification of the percentage of animals with ectopic synapses in the Zone 1 region of AIY in the indicated the genotypes. **(C-F)** Confocal micrographs of EMB-9::mCherry which allowed examination of EMB-9 protein levels in the adult head of wild type animals (C), *cima-1(wy84)* mutants (D), *mig-17(ola226)* mutants (E), and *cima-1(wy84);mig-17(ola226)* double mutants (F). For a developmental characterization of the expression of EMB-9 see Figure S8. **(G)** Quantification of EMB-9::mCherry fluorescence intensity in the indicated genotypes. **(H)** Quantification of the percentage animals with ectopic synapses in the Zone 1 region of AIY in the indicated genotypes. In all graphs, the total number of animals (N) and the number of times scored (n) are indicated in each bar for each genotype as N/n. Statistical analyses are based on two-tailed student’s t-test. Error bars represent SEM, NS: not significant, *p<0.05, ***p<0.001, ****p< 0.0001 as compared to *cima-1(wy84)* mutants (in B and H) and wild type (in G), unless brackets are used between two compared genotypes.

To examine how MIG-17 regulates basement membrane proteins to modulate synaptic allometry, we visualized AIY synaptic phenotypes in mutants with basement membrane proteins defects. Informed by our proteomic data and previously reported *mig-17* interactors (Kim and Nishiwaki, 2015), we focused our genetic studies on EMB-9/Collagen IV α1, OST-1/Sparc, UNC-52/Perlecan and FBL-1/Fibulin.

EMB-9 is a conserved type IV collagen α1, and a core component of the basement membrane (Guo et al., 1991;Sibley et al., 1993;Graham et al., 1997). In neuromuscular junctions, the basement membrane is directly in contact with the neuron-muscle synapses, and recent studies identified roles for EMB-9 in post-embryonic neuromuscular junction morphology (Kurshan et al., 2014;Qin et al., 2014). However, unlike neuromuscular junctions, the AIY synapses are not directly in contact with the basement membrane (BM) (White et al., 1986). The lack of direct contact between the AIY synapses and the BM is similar to most neuron-neuron synapses in the *C. elegans* brain (Kramer, 2005;Hall, 2008), and neuron-neuron synapses in the central nervous system (CNS) of *Drosophila* and vertebrates (Stork et al., 2008;Krishnaswamy et al., 2019).

To examine if EMB-9 affects synapses in AIY, we visualized AIY synapses in *emb-9* mutant alleles. EMB-9 null alleles are embryonic lethal (Guo et al., 1991;Gupta et al., 1997), so we used neomorphic missense alleles which are predicted to result in overabundant or disorganized collagen (Kubota et al., 2012;Kurshan et al., 2014;Qin et al., 2014;Gotenstein et al., 2018). We observed that *emb-9(xd51)* mutants, which display defects in neuromuscular junction synapses (Qin et al., 2014), do not display detectable defects in AIY synapse distribution (Figure S7). However, we did observe that *emb-9(xd51)* and *emb-9(tk75)* alleles significantly suppress the ectopic synapses in *cima-1(wy84)* mutant animals (Figure 8B). These genetic findings are consistent with the proteomic analyses results for EMB-9, and suggest a genetic interaction between the basement membrane and the CIMA-1 and MIG-17 pathway in regulating synaptic allometry. Our findings also indicate that the genetic requirement of basement membrane proteins for synaptic allometry in the AIY interneurons is distinct from the genetic requirement in the maintenance of NMJ morphology.

To further test the relationship between these genes in synaptic allometry, we generated *cima-1(wy84);mig-17(ola226);emb-9(tk75)* triple mutants. We observed that *emb-9(tk75);cima-1(wy84);mig-17(ola226)* did not display enhanced synaptic defects as compared to *cima-1(wy84);mig-17(ola226)* double mutants (Figure 8B). Our genetic and proteomic findings support a model whereby MIG-17 regulates basement membrane proteins, such as EMB-9, to modulate synaptic allometry. The absence of MIG-17, or the presence of neomorphic alleles of EMB-9, result in overabundant or disorganized EMB-9/Collagen IV that suppress CIMA-1, presumably by modulating the material properties of the basement membrane during growth.

### MIG-17 regulates EMB-9/Collagen IV α1 during post-embryonic growth

To better understand the relationship between MIG-17 and EMB-9 in regulating synaptic allometry, we examined EMB-9 protein levels by using an EMB-9::mCherry translational reporter *in vivo* (Ihara et al., 2011). We observed that like MIG-17, EMB-9 protein levels are regulated during development (Figure S8). However, unlike MIG-17, whose protein levels decrease as animals reach the adult stage, EMB-9 protein levels increase as animals progress through the larval stages, achieving maximal expression in the adult stage (Figure S8). Additionally, we observed that in *mig-17(ola226)* mutant animals, EMB-9::mCherry levels were upregulated as compared to wild type (Figures 8C, 8E, 8G). These data are consistent with our genetic and proteomic results, and support the hypothesis that in *mig-17(ola226)* mutants, *cima-1(wy84)* synaptic allometry defects are suppressed due to an upregulation of type IV Collagen protein EMB-9. Moreover while *cima-1* does not affect EMB-9::mCherry levels on its own, we observed that *cima-1* suppresses the effect of *mig-17(ola226)* on EMB-9::mCherry protein levels (Figures 8D, 8F-8G). Our findings support a model in which MIG-17 and CIMA-1 act antagonistically to each other to modulate synaptic allometry in part by regulating basement membrane protein EMB-9 during post-embryonic growth.

Next, we examined if molecules known to modulate the levels or conformation of EMB-9 would similarly affect synaptic allometry. To test this, we imaged the AIY synapses in alleles of *ost-1/*/Sparc, *unc-52*/Perlecan and *fbl-1*/Fibulin, all involved in regulating the trafficking or function of EMB-9 (Kubota et al., 2012;Qin et al., 2014;Morrissey et al., 2016). Loss-of function alleles *ost-1(gk786697, gk193465)*, and *unc-52(gk3)* did not affect the synaptic phenotypes in AIY neurons (Figure S7). However, and consistent with our model, all the alleles predicted to result in overabundant or disorganized EMB-9/Collagen IV significantly suppress the ectopic synapses observed in *cima-1(wy84)* mutants (Figure 8H). Moreover, the gain-of function *fbl-1(k201)* allele (Kubota et al., 2004), which is predicted to result in overabundant EMB-9 (Kubota et al., 2012), similarly suppresses the ectopic synapses observed in *cima-1(wy84)* mutants (Figure 8H). These genetic findings are consistent with our model that basement membrane proteins modulate *cima-1* regulated synaptic allometry.

The *fbl-1(k201)* allele has been previously reported to also modulate the basement membrane and suppress *mig-17* phenotypes in gonad development (Kubota et al., 2004). Therefore, we tested the interaction between *mig-17(ola226)* and *fbl-1(k201)* in the *cima-1(wy84)* mutant background. We found that the AIY synaptic phenotype in *cima-1(wy84)*;*mig-17(ola226)*;*fbl-1(k201)* triple mutants is similar to that observed in *cima-1(wy84);mig-17(ola226)* or *cima-1(wy84)*;*fbl-1(k201)*, indicating that both *fbl-1(k201)* and *mig-17(ola226)* act antagonically to *cima-1* and in the same pathway in the regulation of synaptic allometry (Figure 8H). Our data are consistent with our model that genetic perturbations which result in overabundant or disorganized EMB-9 suppress *cima-1* allometry defects. We note however that our genetic findings are distinct from those reported for gonad development, and suggest that gonad development and synaptic allometry might have different genetic requirements regarding these alleles and their effects on the basement membrane. Together, our findings support the model that MIG-17 regulates synaptic allometry by modulating basement membrane proteins like EMB-9.

### The VCSC glia bridge epidermal-derived growth signals with the muscle-secreted basement membrane to maintain synaptic allometry

Our genetic, proteomic and cell biological findings strongly indicate that *in vivo*, non-neuronal tissues, including epidermal cells, muscle-derived basement membrane and glia, convey growth information to the nervous system to regulate synaptic allometry. To understand how this cross-tissue communication affects glia to regulate synaptic allometry, we examined electron micrographs that show the relationship between the synapses in AIY interneurons, the VCSC glia, the epidermal cells, the basement membrane and the muscles.

The AIY Zone 2 synaptic region lies in the ventral base of the nerve ring bundle and is in direct contact with the basal side of the VCSC glia (White et al., 1986;Altun, 2019). No basement membrane is present between the VCSC glia and the nerve ring neurons (Figures 9A-9B). On their apical side, the VCSC glia contact two distinct non-neuronal tissues: epidermal cells and muscle-derived basement membrane. VCSC are observed to directly contact epidermal cells, a cell-cell adhesion relationship which we had previously shown is regulated by epidermally-expressed CIMA-1 and the ecto-domain of EGL-15/FGF Receptor (Shao et al., 2013). No basement membrane is present between the VCSC glia and the epidermal cells. However, at regions where the glia are apposed to muscle cells, we observed VCSC glia decorated with basement membrane on the apical side that faces the pseudocoelom, a cavity that contains internal fluids. VCSC glia therefore have three surface regions: direct contact, through the basal side, with neurons, direct contact, through the apical side, with the epidermal cells, and contact with muscle-derived basement membrane (Figure 9A’).

**Figure 9.**
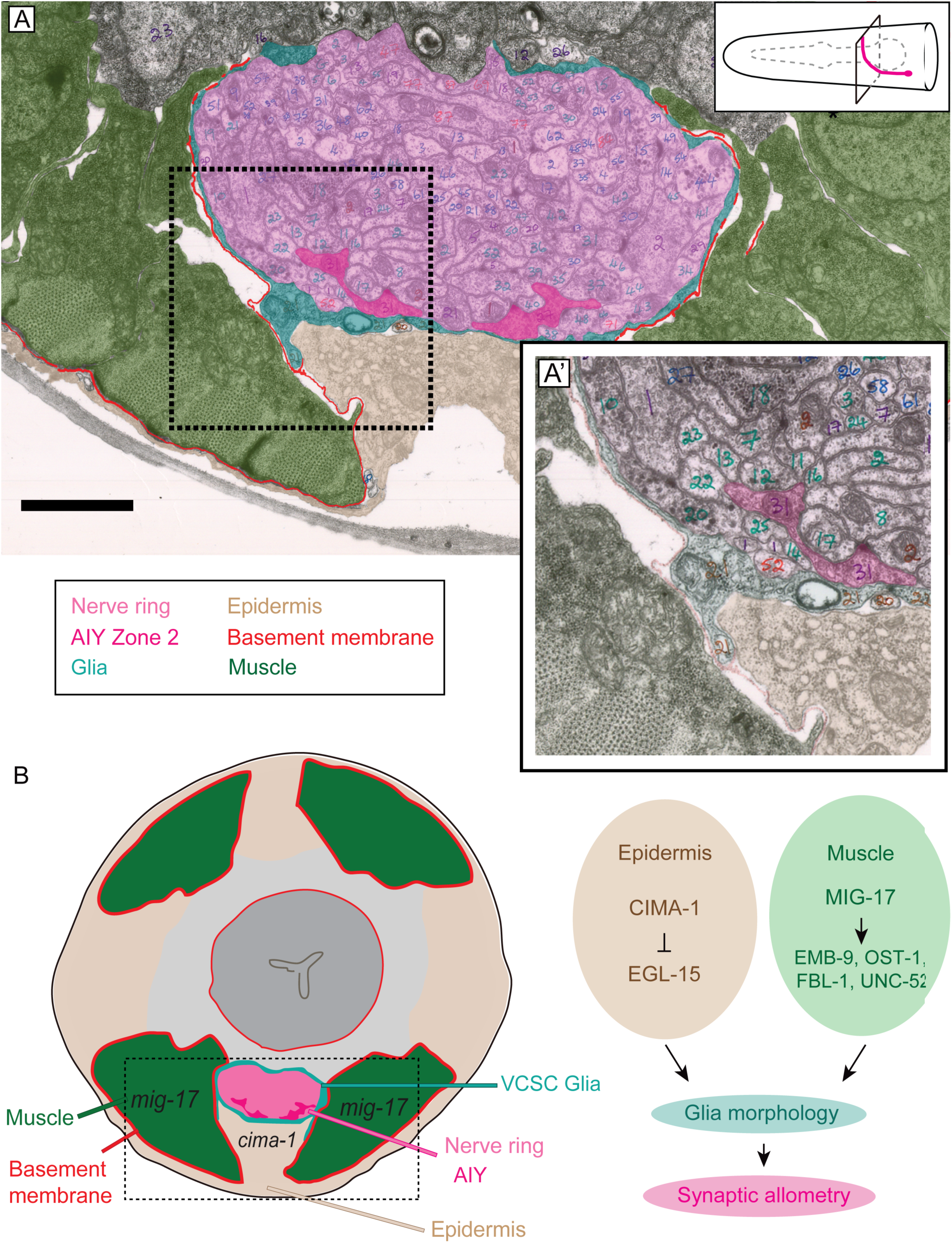
Glia maintain synaptic allometry by bridging epidermal-derived growth signals with the muscle-secreted basement membrane. **(A-A’)** Segmented electron micrograph from a wild type animal (JSH236 from (White et al., 1986). The EM corresponds to the Zone 2 region of AIY, with marked muscles (pseudo-colored green), basement membrane (BM, pseudo-colored red), VCSC glia (pseudo-colored teal), epidermal syncytium (pseudo-colored beige) and the ventral bundle of the nerve ring (pseudo-colored pink, including AIY Zone 2 pseudo-colored dark pink). In (A’), the yellow-boxed region is enlarged and the pseudo-coloring opacity is decreased as to show that the basement membrane, specifically observed between muscle and VCSC glia, but not between glia and epidermal cells or between glia and neurons. **(C)** A cartoon diagram depicting the cross section of *C. elegans* nerve ring as shown in A (modified from WormAtlas.org). As illustrated in the cartoon and the EM image, body wall muscle (green), the nerve ring (pink) and glia (teal) are proximal to the epidermal cells (beige). The nerve ring bundle is surrounded by VCSC glia, which contact it directly at the glia basal side. At the glia apical side, glia interact with muscle-derived basement membrane (red) and epidermal cells (beige). To the right of the schematic, a molecular and cellular model of our *in vivo* data demonstrating the role of non-neuronal cell in glia position and morphology to regulate synaptic allometry during growth.

Given the anatomical relationship between these tissues, we hypothesized that if our model were correct, release of muscle-derived *mig-17* into the pseudocoelom would be sufficient to rescue the AIY synaptic phenotype in *mig-17* mutants. We decided to ectopically express MIG-17 from the epidermal cell or VCSC glia and examine *mig-17* rescue. Both epidermal cells and glia in this region face the pseudocoelom cavity proximal to the basement membrane. Consistent with our model, we observed that expressing *mig-17* in the epidermal cells (via the *dpy-7* promoter) or in VCSC glia (via the *hlh-17* promoter) rescues the *mig-17* suppression in *cima-1(wy84);mig-17(ola226)* mutants (FigureS9). However, expressing *mig-17* from the neurons that face the basal side of the VCSC glia, and do not contact the pseudocoelom or the basement membrane, did not rescue these defects (Figure 5E and S9). Our data collectively indicate that *mig-17* is secreted into the pseudocoelom by body wall muscles to modulate the basement membrane. The basement membrane decorates the basal side of the glia, and with the epidermal cells, jointly modulate glia morphology and position to regulate synaptic allometry during growth (Model in Figure 9B).

## DISCUSSION

Glia regulate synaptic positions during growth. In *C. elegans*, synaptic positions of the interneuron AIY are established early in embryogenesis and maintained during growth to preserve circuit integrity (Shao et al., 2013). Our studies have determined that glia play critical roles, both during embryonic development and during post-embryonic growth, in maintaining synaptic positions. During embryonic development, VCSC glia secrete a chemotrophic factor (Netrin) to coordinate synaptic specificity between AIY and its post-synaptic partners at a glia-specified coordinate (Colon-Ramos et al., 2007). During post-embryonic growth, the same VCSC glia are required for maintaining synaptic positions, but through distinct, Netrin-independent signaling pathways (Shao et al., 2013). Our studies reveal two important features regarding maintenance of synaptic allometry. First, while dependent on the same guidepost cells, synaptic positions are regulated through distinct molecular pathways during embryonic development and post-embryonic allometric growth. Second, our findings underscore the *in vivo* importance of non-neuronal cells, in particular glia, in determining synaptic position. In support of this, we observe, through genetic and *in vivo* cell biological studies, that maintenance of synaptic allometry depends on the relative position of the glia end-feet with respect to the AIY neurite. Specifically, altering glia positions and AIY positions only result in synaptic phenotypes if their region of contact (normally in Zone 2) is altered. Our findings are consistent with vertebrate and invertebrate studies supporting essential roles for glia in regulating synaptic assembly and function *in vivo* (Allen and Eroglu, 2017;Van Horn and Ruthazer, 2019), and extend these findings to highlight a role for glia in maintaining synaptic positions during post-embryonic allometric growth.

Glia morphology and positions are actively maintained during growth. Growth in *C. elegans* is coordinated by epidermal cells and body wall muscles (Chisholm and Hardin, 2005). Epidermal cells express genes that regulate molting, body morphogenesis and animal size (Chisholm and Hsiao, 2012;Chisholm and Xu, 2012). Body wall muscle contractions regulate elongation during embryogenesis, and influence epidermal cytoskeletal remodeling via tension-sensing mechanisms (Williams and Waterston, 1994;Chisholm and Hsiao, 2012;Chisholm and Xu, 2012). While we do not yet understand how growth is sensed in organisms, our findings uncover a cooperative signaling pathway that emerge from these two growth-regulating cell types to position glia during allometry. Our genetic studies indicate that muscle-derived MIG-17 is epistatic to epidermally-derived CIMA-1 and EGL-15, indicating a multi-tissue, non-neuronal pathway that converges in transducing growth information to position glia and regulate synaptic allometry. In this way our findings uncover a non-cell autonomous, two component system that cooperates to transduce growth information to the nervous system through glia.

ADAMTS protease MIG-17 regulates the basement membrane during post-embryonic growth to modulate synaptic allometry. In *Drosophila*, homologous ADAMTS Stl and AdamT-A proteins are required for the development of the peripheral nervous system and the maintenance of the central nervous system architecture (Lhamo and Ismat, 2015;Skeath et al., 2017). In humans, lesions in ADAMTS genes result in biomedically important defects, including short stature and neuronal developmental disorders, among other problems (Miguel et al., 2005;Howell et al., 2012;Cheng et al., 2018). In all organisms, ADAMTS metalloproteases function in the degradation and remodeling of the extracellular matrix (Krishnaswamy et al., 2019). The remodeling of the extracellular matrix in *C. elegans* is important during gonad organogenesis and during pharynx growth, and is mediated in part by the MIG-17 metalloprotease (Nishiwaki et al., 2000;Kubota et al., 2004;Kubota et al., 2008;Kim and Nishiwaki, 2015;Shibata et al., 2016). Most of our understanding of extracellular matrix structures, such as the basement membrane, are derived from studies examining its assembly during embryogenesis. Less is known about how the basement membrane changes during post-embryonic growth (Jayadev and Sherwood, 2017). Our proteomic, genetic and cell biological findings strongly suggest that the basement membrane is a dynamic structure that remodels in part through the activity of MIG-17 metalloproteases. Disruption of basement membrane components hypothesized to affect the biophysical properties of the extracellular matrix result in defective glia positions during growth, and affect synaptic allometry. Together our findings indicate an important role for the regulated remodeling of the basement membrane in orchestrating intercellular interaction when animals expand their volume during growth.

A muscle-epidermal-glia signaling axis, mediated through MIG-17 dependent regulation of the extracellular matrix, is necessary for modulating glia morphology and synaptic positions. Basement membrane proteins have been previously shown to regulate neuromuscular junction synapses (Ackley et al., 2003;Patton, 2003;Kurshan et al., 2014;Qin et al., 2014;Rogers and Nishimune, 2017). Neuromuscular junctions are in direct contact with the basement membrane, while the neurons examined in this study, which are in the nerve ring, are not in direct contact with the basement membrane (White et al., 1986). We find that for nerve ring synapses, MIG-17 regulation of the basement membrane and synaptic allometry is mediated by glia. VCSC glia ensheath the nerve ring to form a physical barrier between the neuropil and adjacent tissues, including the pseudocoelom, the basement membrane and the epidermal cells (Shaham, 2015). At the basal side VCSC glia contact neurons in the nerve ring, while at the apical side VCSC glia are either decorated by basement membrane or in direct contact with epidermal cells. The relationship of the VCSC glia, basement membrane and epidermal cells reflect the genetic relationship uncovered in our forward genetic screens, with epidermal CIMA-1 and EGL-15/FGFR modulating glia morphology through epidermal-glia adhesion, and muscle-derived MIG-17 modulating glia morphology through the extracellular matrix.

The muscle-epidermal-glia signaling axis we uncover here is reminiscent of the neurovascular unit of the blood brain barrier of Drosophila and vertebrates. In the vertebrate neurovascular unit, muscle-related pericyte cells interact with vascular endothelial cells and astrocytes through the basement membrane (Xu et al., 2019). Pericytes, endothelial cells and basement membrane are not in direct contact with neurons. Instead, astrocytes mediate signaling between these non-neuronal cells and neurons, including the coupling of developmental programs that coordinate vasculature development and neurodevelopment (Tam and Watts, 2010), and the coupling of functional programs that coordinate neuronal activity with blood flow (Allan, 2006;Koehler et al., 2009). We note that the extracellular matrix of the blood brain barrier is molecularly similar to that of the basement membrane of *C. elegans,* and includes molecules tested in this study such as laminin, collagen IV and fibulin (Thomsen et al., 2017). While the role of these components in synaptic allometry has not been examined in vertebrates, it is intriguing to speculate that the functional neurovascular unit might help transduce information from the vasculature to mediate synaptic positions during allometric growth. We also hypothesize that analogous structures to the neurovascular unit might represent conserved signaling axis that couple glia-mediated communication between non-neuronal cells and neurons in metazoans.

## ACKNOWLEDGEMENTS

We thank Z.F. Altun and D.H. Hall from WormAtlas for help with schematic figures. We also thank Shiqing Cai, Yidong Shen, Kiyoji Nishiwaki, Mei Ding (Chinese Academy of Sciences), David Sherwood (Duke University) and the Caenorhabditis Genetic Center (funded by NIH (P40 OD010440) for providing strains and plasmids. We thank members in Shao lab and the Colón-Ramos lab for insightful discussions on the work and advice on the project. We thank Mi Zhou for providing technical support on image acquisition. We thank the Research Center for Minority Institutions program, the Marine Biological Laboratories (MBL), and the Instituto de Neurobiología de la Universidad de Puerto Rico for providing meeting and brainstorming platforms. Research in the Z.S. lab was supported by the National Natural Science Foundation of China (31471026, 31872762), Shanghai Municipal Science and Technology Major Project (No. 2018SHZDZX01) and ZJLab. Research in the DAC-R lab was supported by NIH R01NS076558, DP1NS111778 and by an HHMI Scholar Award.

## AUTHORS’ CONTRIBUTION

TJ, KW, JH, MW, XZ, ZS and DAC-R designed experiments. TJ, KW, LF, JH, MW, HD, JS, MW and ZS performed experiments. TJ, KW, LF, JH, MW, XZ, LAM, ZS and DAC-R analyzed and interpreted the data. TJ, ZS and DAC-R wrote the paper.

---

**Figure S1 Model of CIMA-1 site of action**

**(A, C)** Cartoon diagrams of the head of the *C. elegans* of wild type (A) and *cima-1* mutant animals (C). In both cartoons, the epidermal syncytium is represented in beige, ventral cephalic sheath cell (VCSC) glia in red and the AIY interneuron in green. Blue dashed lines indicated sites of epidermal-glia contact. The cartoons are graphical abstracts of the findings of (Shao et al., 2013). In wild type animals, CIMA-1 acts in epidermal cells to suppresses the epidermally-derived FGF Receptor/EGL-15, which in turn maintains VCSC glia morphology, probably by mediating adhesion between the epidermal cell and glia. In *cima-1* loss-of-function mutants (C), EGL-15 protein levels are upregulated, and this promotes VCSC glia endfeet extension, allowing ectopic contact with the AIY Zone 1 region and promoting formation and stability of ectopic synapses in AIY Zone 1. **(B, D)** Confocal micrographs of the VCSC glia (red) and the AIY interneurons (green) with bright field in adult wild type (B) and *cima-1(wy84)* mutant animals (D) for the region in the dashed box (A and C). Brackets indicate AIY Zone 1 region; asterisks indicate Zone 2 region. Scale bar, 10μm.

**Figure S2. Synaptic allometry is affected by the size of the animal**

**(A-C)** Images of wild type (A), *cima-1(wy84);dpy-4(e1166)* (B) and *cima-1(wy84);lon-3 (e2175)* mutant animals. The dumpy or long mutants are ∼25% shorter or longer than wild type animals, respectively. **(D-H’)** Confocal micrographs of the AIY presynaptic sites (visualized with GFP::RAB-3, green) in adult wild type (D, D’), *cima-1(wy84);dpy-4(e1166)* (E, E’), *cima-1(wy84);lon-3(e2175)* (F, F’), *cima-1(wy84)* (G,G’) and *cima-1(wy84);ola226* (H, H’) with bright field (D, E, F, G, H) or without bright field (D’, E’, F’, G’, H’). Brackets indicate the AIY Zone 1 region; asterisks indicate the Zone 2 region. Scale bar in A is 200μm and applies to B and C; scale bar in D is 10μm and applies to all fluorescent micrographs. **(I)** Quantification of the total animal length of wild type, *cima-1(wy84)*, *ola226* and *cima-1(wy84);ola226* mutants. In the graph, the total number of animals (N) and the number of times scored are indicated in each bar for each genotype as N/n. Error bars represent SEM. Statistical analyses are based on two-tailed student’s t-test, NS: not significant or p>0.05.

**Figure S3. *ola226* affects AIY neurite and cell body position**

**(A)** A cartoon diagram of AIY (red) in the *C. elegans* head. The orange arrow indicates the cell body, bracket indicates the Zone 1 region, asterisk indicates Zone 2 region, and vertical dashed line indicates center of pharynx bulb here and in micrographs B-C. Lengths scored for cell body position (graph in D) and Zone 1 length (graph in E) are shown. **(B-C)** Confocal micrographs of AIY (red) with bright field microscopy for wild type (B) and *ola226* mutants (C). In (B), a co-marker used in the study is also visible proximal to AIY. Importantly for the study, while AIYs have the same shape in the inspected genotypes, note the differences in the AIY neurite and cell body positions between these two genotypes as compared to the pharynx bulb (visible with bright field). **(D-E)** Quantification of the position of AIY (length between the tip of pharynx and the AIY cell body, as indicated in the schematic in (A)) (D), and quantification of the length of Zone 1 (E) in wild type and *ola226* mutant animals. In the graph, the total number of animals (N) and the number of times scored (n) are indicated in each bar for each genotype as N/n. Error bars represent SEM. Statistical analyses are based on two-tailed student’s t-test, **** p<0.0001.

**Figure S4. Synaptic phenotypes in *mig-17* alleles**

**(A-C)** Confocal micrographs of the AIY synaptic vesicle marker GFP::RAB-3 (green) in the wild type (A), *mig-17(ola226)* mutant (B) and *mig-17(k113)* mutant (C). The scale bar in A is 10μm and applies to all panels. Brackets mark the Zone 1 region; asterisks indicate the Zone 2 region. **(D)** Quantification of the percentage of adult animals with ectopic synapses in the AIY Zone 1 region. In the graph, the total number of animals (N) and the number of times scored (n) are indicated in each bar for each genotype as N/n. Statistical analyses are based on two-tailed student’s t-test, Error bars represent SEM, NS: not significant or p>0.05.

**Figure S5. *mig-17(ola226)* and *cima-1(wy84)* phenotypes in pharyngeal length**

**(A)** A cartoon diagram of the pharynx in the *C. elegans* head. The pharynx is outlined with a dashed grey line, and the length quantified in (B) is shown. **(B)** Quantification of pharyngeal length of wild type, *cima-1(wy84)*, *mig-17(ola226)* and *cima-1(wy84);mig-17(ola226)* double mutants. The data indicate that both *cima-1(wy84)* and *mig-17(ola226)* significantly increase the pharyngeal length, consistent with (Shibata et al., 2016), and that the double mutant enhances the single mutant effects. The data indicate that unlike the AIY presynaptic phenotype, *mig-17(ola226)* and *cima-1(wy84)* cooperate, rather than antagonize, each other in regulating pharyngeal length. In the graph, the total number of animals (N) and the number of times scored (n) are indicated in each bar for each genotype as N/n. Statistical analyses are based on two-tailed student’s t-test. Error bars represent SEM, *p<0.05, ****p<0.0001 as compared to wild type or between indicated genotypes.

**Figure S6 CRISPR strategies to generate endogenous MIG-17::mNeonGreen and the *mig-17(shc8)* allele**

**(A)** Schematic of the strategy used to fuse to the C-terminus of the endogenous genomic *mig-17* locus with mNeonGreen (Dickinson et al., 2013). Briefly and as indicated, two sgRNAs (blue) were used to insert mNeonGreen::3×Flag into the MIG-17 C-terminus via CRISPR. The *mig-17* exons are indicated in the schematic by black boxes; the green and purple boxes represent mNeonGreen and 3xflag tags. Two synonymous mutations at the PAM sites were made on the repair templates (orange) to avoid cutting by Cas9. **(B)** Schematic of the strategy used to change E303 at MIG-17 and generate the *mig-17(shc8)* allele. The location for the two sgRNAs used (black bolded and blue) and the *shc8* (E303A) repair template are indicated. The repair template comprised of 1.2 kb upstream and 1.2 kb downstream genomic sequence flanking the enzymatic active site E303. The Glutamic acid (GAA) at the 303 was changed into Alanine (GCA) (red). Eight synonymous mutations were made to prevent Cas9 from cutting the repair template (orange). sgRNA design is based on sgRNA design tool (http://crispr.mit.edu).

**Figure S7 Phenotypes of basement membrane genes in AIY synapses**

Quantification of the percentage of adult animals with ectopic synapses in the AIY Zone 1 region in *ost-1(gk786697, gk193465)*, *emb-9(xd51)*, *fbl-1(k201)*, or *unc-52(gk3)* mutants. In the graph, the total number of animals (N) and the number of times scored (n) are indicated in each bar for each genotype as N/n. Statistical analyses are based on two-tailed student’s t-test, Error bars represent SEM, NS: not significant or p>0.05.

**Figure S8. EMB-9 is developmentally regulated**

**(A)** Cartoon diagram of the head of *C. elegans*, similar to Figure 1A. The dotted box indicates the region imaged in the subsequent micrographs. **(B-F)** Confocal micrographs of animals expressing EMB-9::mCherry (Ihara et al., 2011), imaged at larva stage 1 (L1 in (B)), larva stage 2 (L2 in (C)), larva stage 3 (L3 in (D)), larva stage 4 (L4 in (E)) and 1 day-old adults (F). For all images, scale bars are 10μm. **(G-H)** Quantification of EMB-9::mCherry intensity at different developmental stages (G) and the p-value for paired comparison based on two-tailed student’s t-test (H). In the graph, the total number of animals (N) and the number of times scored (n) are indicated in each bar for each genotype as N/n. Statistical analyses are based on two-tailed student’s t-test. Error bars represent SEM.

**Figure S9 Ectopic expression of *mig-17* in epidermal cells or VCSC glia rescues the *mig-17* suppression in *mig-17(ola226);cima-1(wy84)* mutants**

Quantification of the percentage of adult animals with ectopic synapses in the AIY Zone 1 region in the indicated genotypes. The *cima-1(wy84);mig-17(ola226)* no-transgene control, and the body wall muscle rescue control, are the same as in Figure 5E (and were scored at the same time as the experimentals shown here). In the graph, the total number of animals (N) and the number of times scored (n1) are indicated in each bar for each genotype, as are, for the transgenic lines created, the number of transgenic lines (n2) examined (all using the convention N/n1/n2). Statistical analyses are based on two-tailed student’s t-test. Error bars represent SEM, ****p< 0.0001 as compared to the no-transgene control.

**Supplemental Table S1. Strains used in this study**

Strains and the corresponding genotypes are listed in the table S1.

**Supplemental Table S2. Constructs used in this study**

The name of constructs, the primers and the vectors for building the constructs are listed in the table S2. Detailed cloning information is available upon request.

**Supplemental Table S3. Protein levels altered in *mig-17(ola226)* as detected by LS/MS proteomic analyses.**

Proteins upregulated (>1.2 fold) or downregulated (<0.8 fold) are listed in the spreadsheet. Note that *mig-17(ola226)* was isolated from forward genetic screen, which would introduce other background mutations. Further analyses are required to confirm if the protein level changes are due specifically to the *mig-17(ola226)* mutation.

