## Supplementary figures and images for "ADAMTS-family protease MIG-17 regulates synaptic allometry by modifying the extracellular matrix and modulating glia morphology during growth"

### Supplemental Figure 1

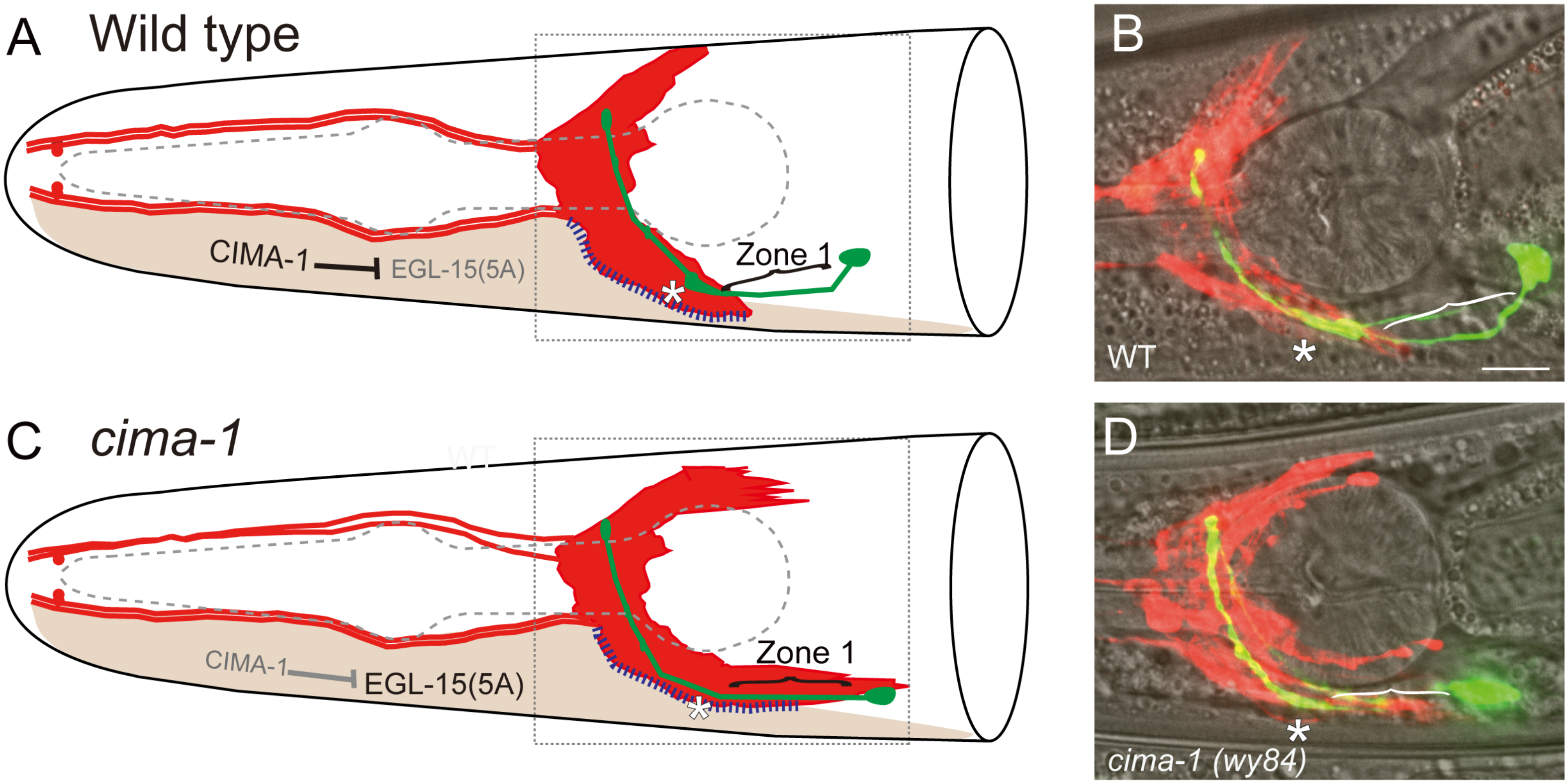

### Supplemental Figure 2

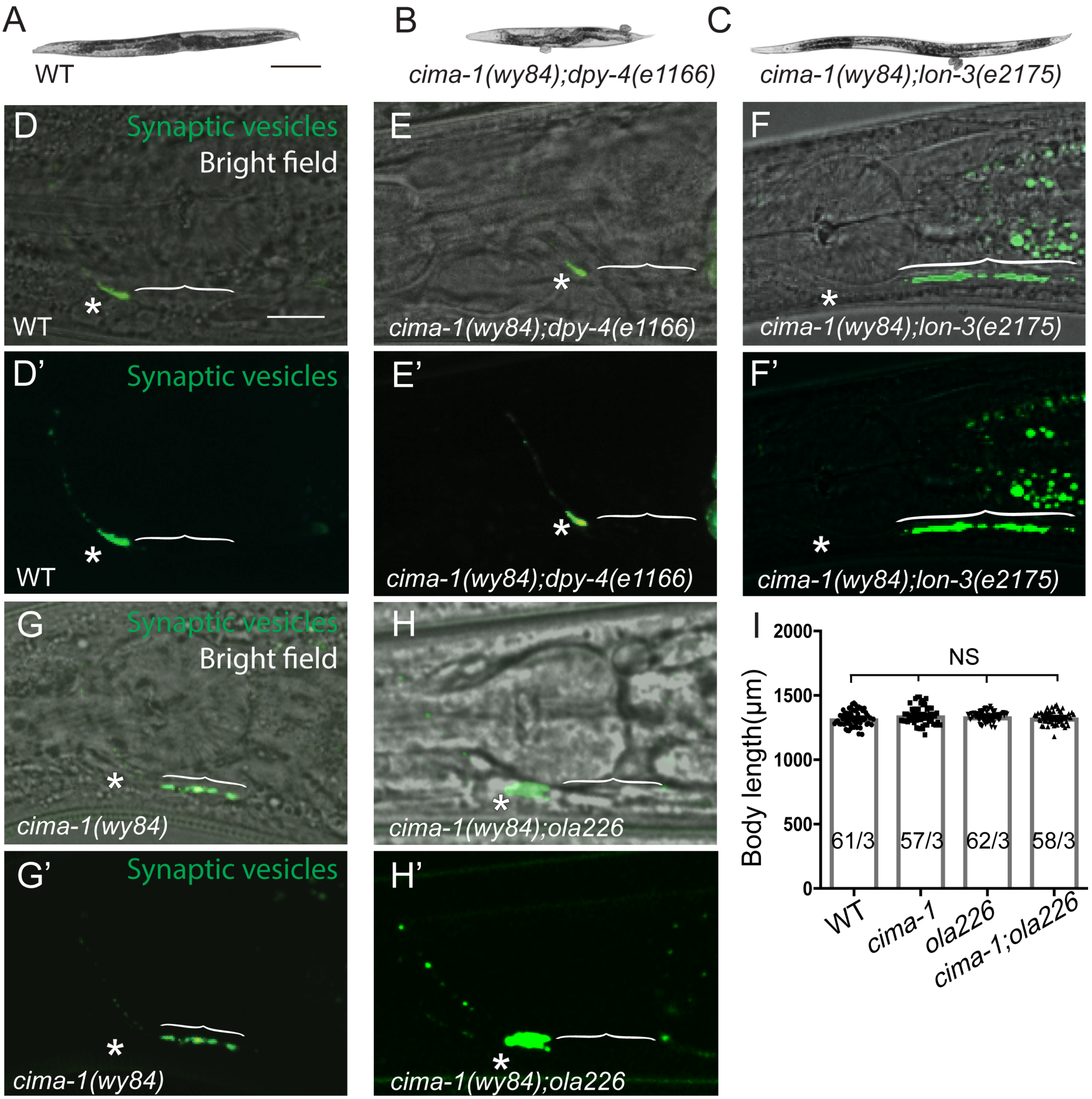

### Supplemental Figure 3

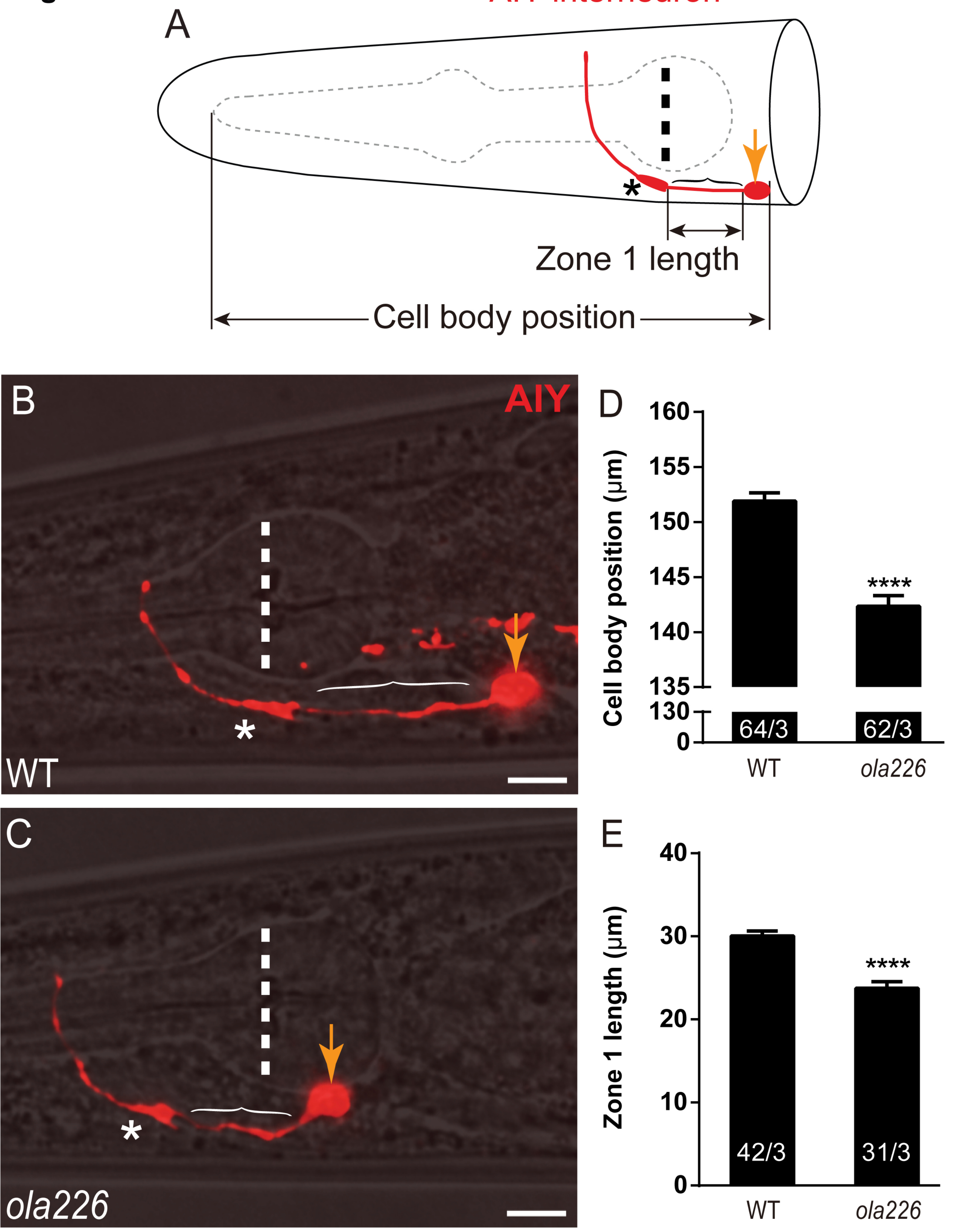

### Supplemental Figure 4

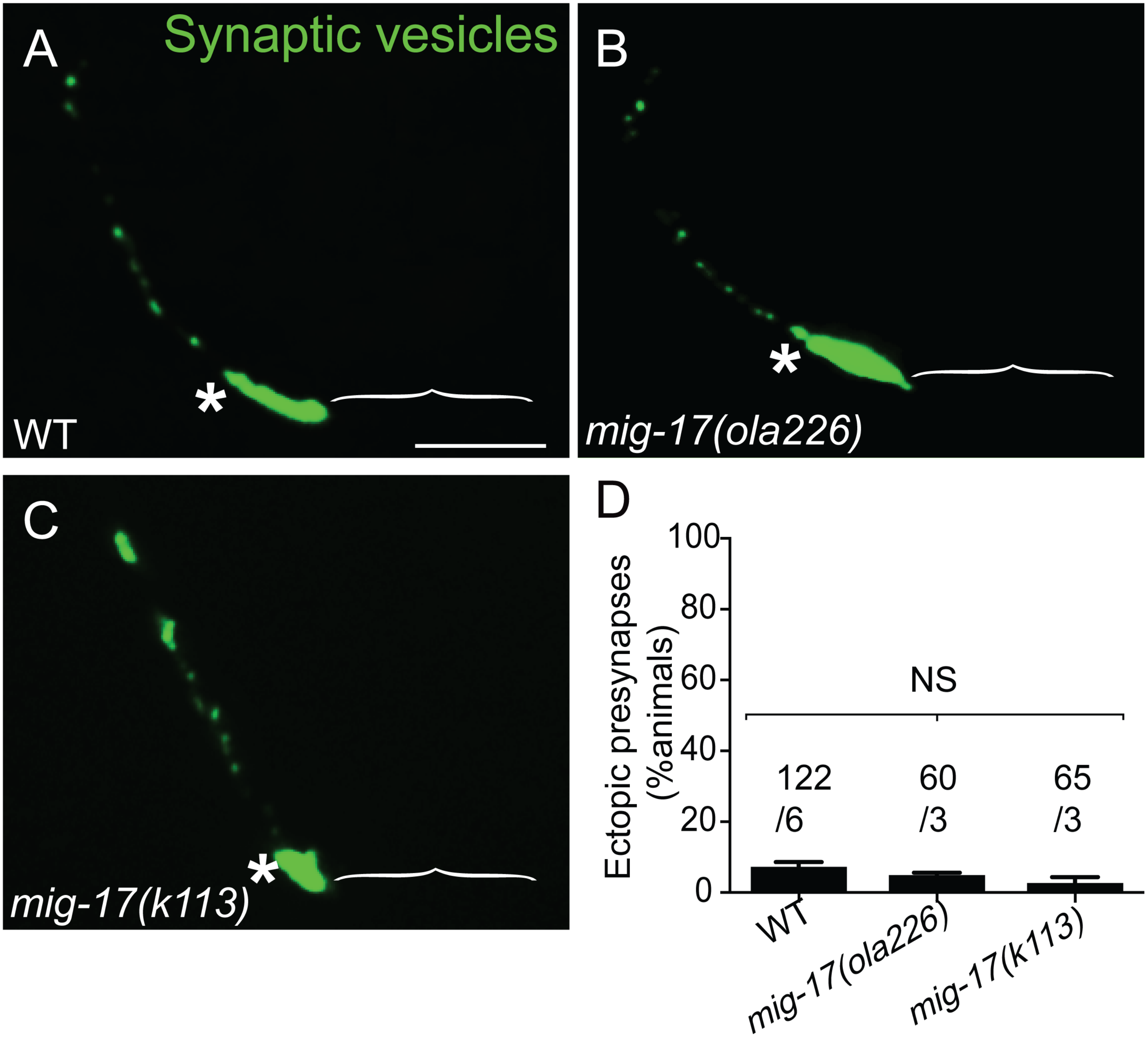

### Supplemental Figure 5

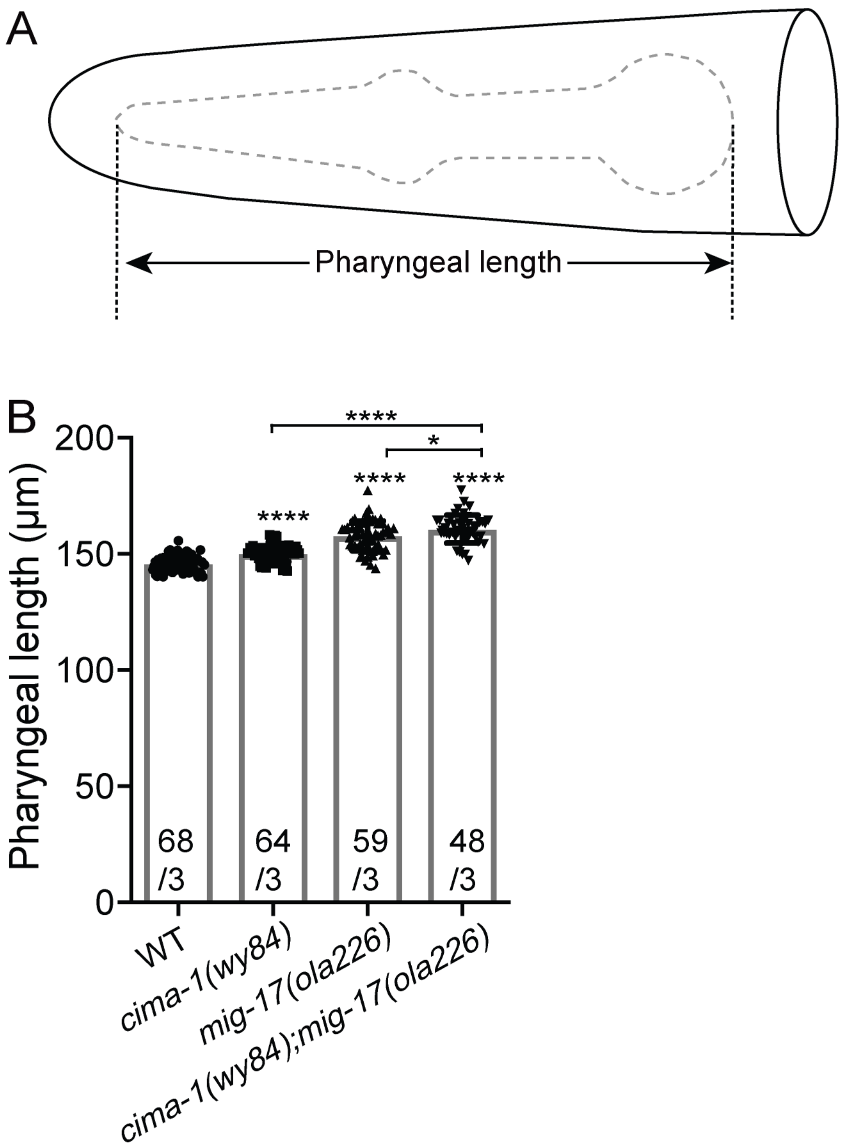

### Supplemental Figure 6

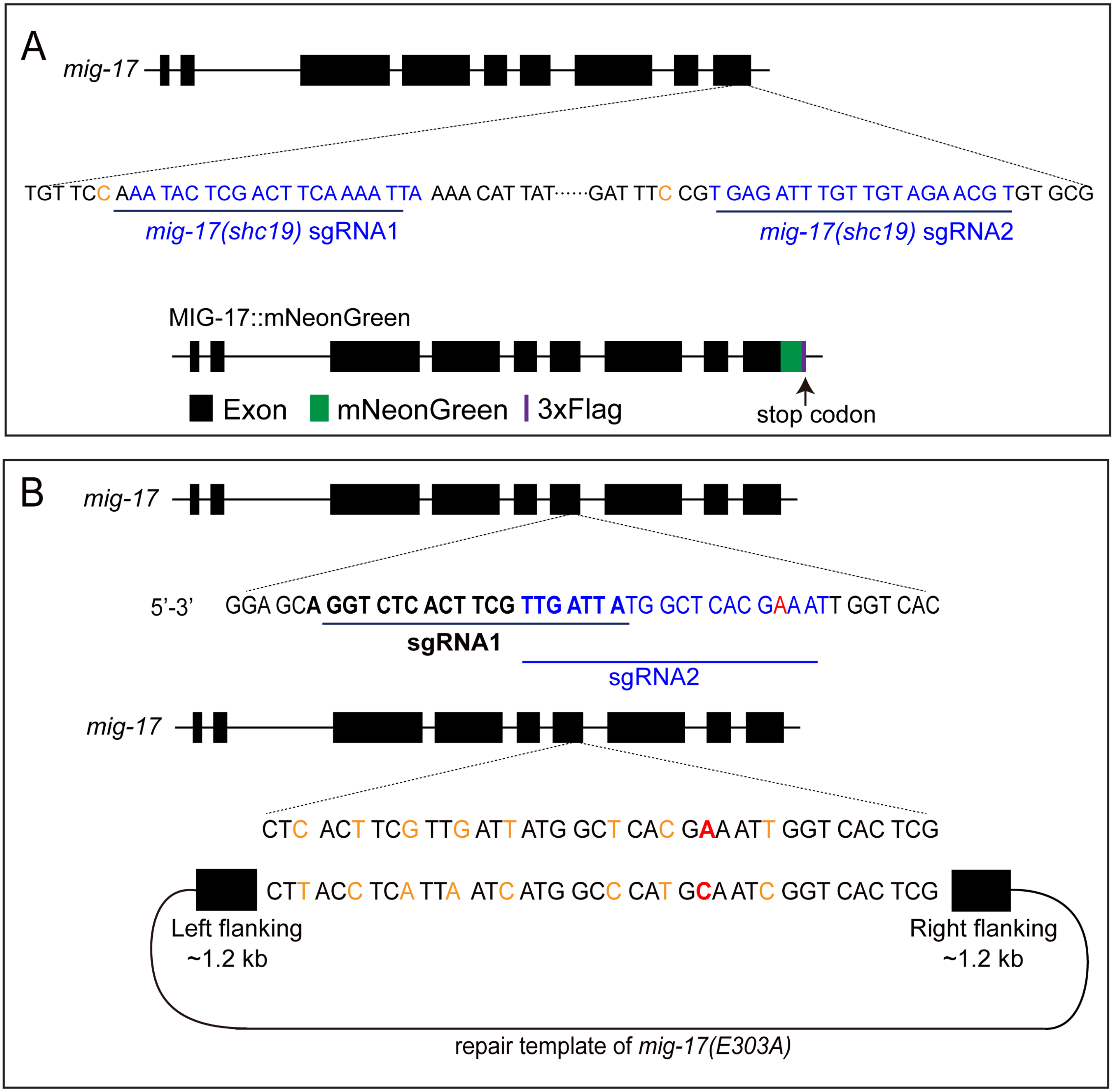

### Supplemental Figure 7

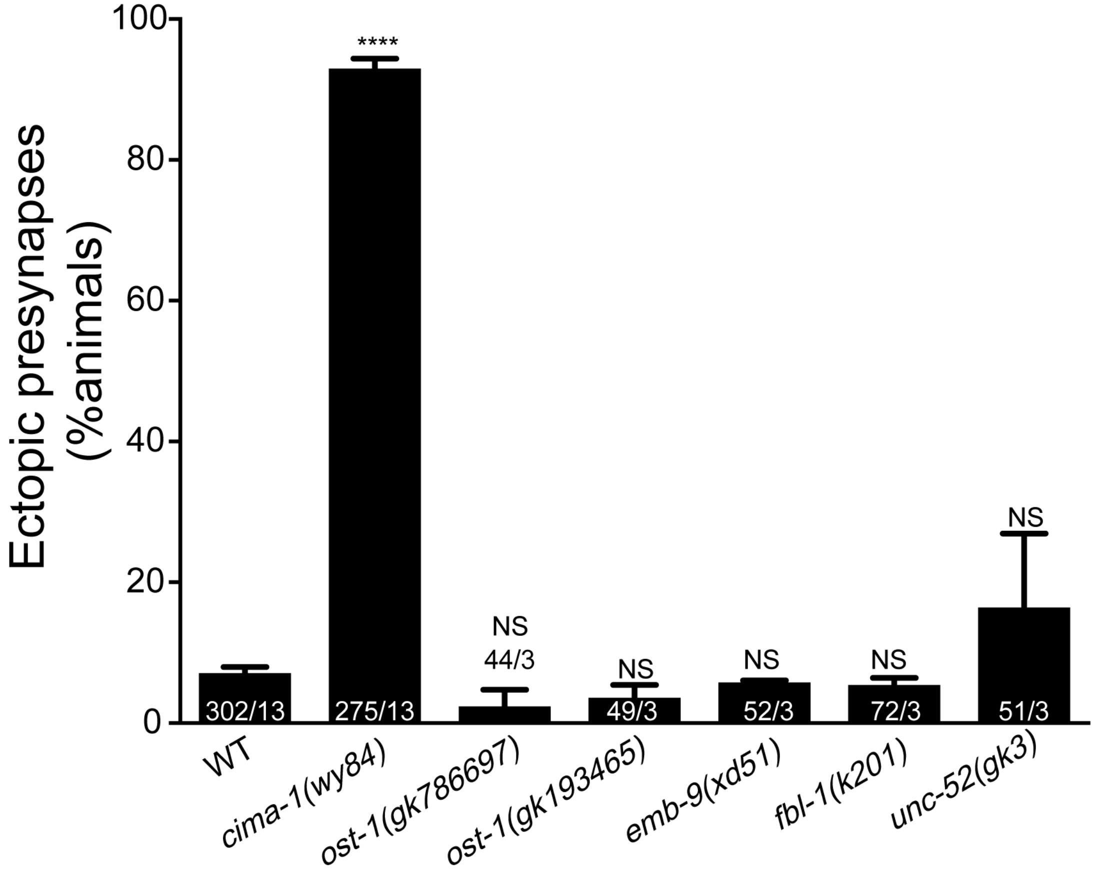

### Supplemental Figure 8

#### Figure S8

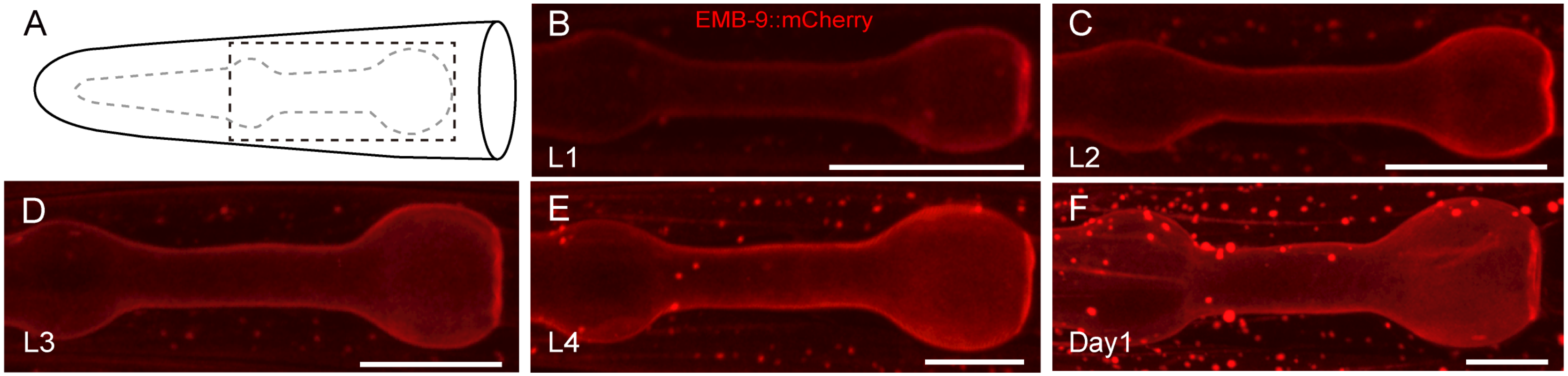


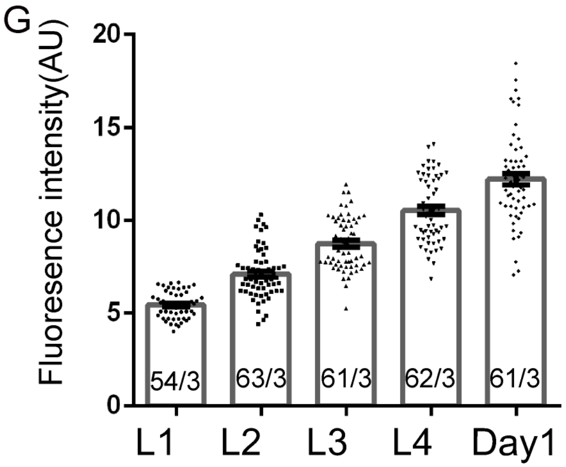


G'

P value for each set of comparison

| | L1 | L2 | L3 | L4 |
|---|---|---|---|---|
| L2 | <0.0001 | | | |
| L3 | <0.0001 | <0.0001 | | |
| L4 | <0.0001 | <0.0001 | <0.0001 | |
| Day1 | <0.0001 | <0.0001 | <0.0001 | <0.0001 |

### Supplemental Figure 9

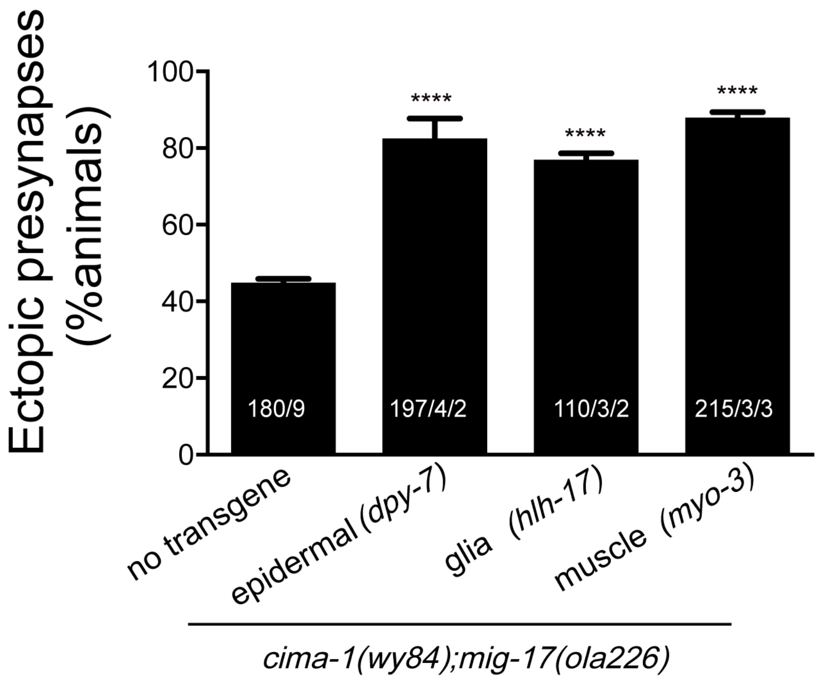
