## Supplemental Table 1 for "ADAMTS-family protease MIG-17 regulates synaptic allometry by modifying the extracellular matrix and modulating glia morphology during growth"

| Table S1. Strains used in this study | |
|---|---|
| **Strain name** | **Genotype** |
| N2 | Wild type |
| FDU2215 | *shcEx1126[Pttx-3::syd-1::GFP;Pttx-3::rab-3::mCherry;Punc-122::RFP]* |
| FDU2216 | *cima-1(wy84)* IV*;shcEx1127[Pttx-3::syd-1::GFP;Pttx-3::rab-3::mCherry;Punc-122::RFP]* |
| FDU2217 | *cima-1(wy84)* IV*;mig-17(ols226)* V*;shcEx1128[Pttx-3::syd-1::GFP;Pttx-3::rab-3::mCherry;Punc-122::RFP]* |
| TV392 | *wyIs45 [Pttx-3::GFP::rab-3; Punc-122::RFP]* X |
| FDU937 | *cima-1(wy84)* IV; *wyIs45 [Pttx-3::GFP::rab-3; Punc-122::RFP]* X |
| FDU2095 | *cima-1(wy84)* IV; *mig-17(ola226)* V; *wyIs45 [Pttx-3::GFP::rab-3; Punc-122::RFP]* X |
| FDU2333 | *cima-1(wy84)* IV; *mig-17(ola226)* V; *wyIs45 [Pttx-3::GFP::rab-3; Punc-122::RFP]* X; *shcEx1146 [Pmig-17::mig-17 genomics; Phlh-* |
| FDU2334 | *cima-1(wy84)* IV; *mig-17(ola226)* V; *wyIs45 [Pttx-3::GFP::rab-3; Punc-122::RFP]* X; *shcEx1147 [Pmig-17::mig-17 genomics; Phlh-* |
| FDU882 | *cima-1(wy84)* IV; *mig-17(k113)* V; *wyIs45 [Pttx-3::GFP::rab-3; Punc-122::RFP]* X |
| FDU2219 | *shcEx1129[Pmig-17::mig-17::SL2::GFP; Pdpy-4::mCherry]* |
| FDU2220 | *shcEx1130[Pmig-17::mig-17::SL2::GFP; Pmyo-3::mCherry]* |
| FDU2318 | *shcEx1131[Pmig-17::mig-17::SL2::GFP; Phlh-17::mCherry]* |
| FDU2462 | *shcEx1410[Pmig-17::mig-17::SL2::GFP; Prab-3::mCherry]* |
| FDU1056 | *mig-17(shc19)V* |
| FDU1778 | *mig-17(shc19)* V; *shcEx845[Phlh-17::mCherry]* |
| FDU2332 | *mig-17(shc19)* V; *shcEx1145[Pdpy-4::mCherry]* |
| FDU1896 | *mig-17(shc19)* V; *shcEx1402[ Pmyo-3::mCherry]* |
| FDU1891 | *mig-17(shc19);shcEx1403[Prab-3::mCherry]* |
| FDU2598 | *mig-17(shc19)V; qyIs46[unc119;emb-9::mCherry]X* |
| FDU882 | *cima-1(wy84)* IV; *mig-17(k113)* V; *wyIs45 [Pttx-3::GFP::rab-3; Punc-122::RFP]* X |
| FDU2214 | *cima-1(wy84)* IV; *mig-17(shc8)* V; *wyIs45 [Pttx-3::GFP::rab-3; Punc-122::RFP]* X |
| FDU2315 | *cima-1(wy84)* IV; *mig-17(k174)* V; *wyIs45 [Pttx-3::GFP::rab-3; Punc-122::RFP]* X |
| FDU2596 | *cima-1(wy84) IV; mig-17(ola226) V; wyIs45 [Pttx-3::GFP::rab-3; Punc-122::RFP] X; shcEx1414 [Pmig-17::mig-17(E303A); Phlh-17::mCherry]* |
| FDU2597 | *cima-1(wy84) IV; mig-17(ola226) V; wyIs45 [Pttx-3::GFP::rab-3; Punc-122::RFP] X; shcEx 1415[Pmig-17::mig-17(E303A); Phlh-17::mCherry]* |
| FDU2320 | *cima-1(wy84)* IV; *mig-17(ola226)* V; *wyIs45 [Pttx-3::GFP::rab-3; Punc-122::RFP]* X; *shcEx1133 [Pmyo-3::mig-17; Phlh-17::mCherry]* |
| FDU2321 | *cima-1(wy84)* IV; *mig-17(ola226)* V; *wyIs45 [Pttx-3::GFP::rab-3; Punc-122::RFP]* X; *shcEx1134 [Pmyo-3::mig-17; Phlh-17::mCherry]* |
| FDU2322 | *cima-1(wy84)* IV; *mig-17(ola226)* V; *wyIs45 [Pttx-3::GFP::rab-3; Punc-122::RFP]* X; *shcEx1135 [Pmyo-3::mig-17; Phlh-17::mCherry]* |
| FDU2323 | *cima-1(wy84)* IV; *mig-17(ola226)* V; *wyIs45 [Pttx-3::GFP::rab-3; Punc-122::RFP]* X; *shcEx1136 [Punc-14::mig-17; Phlh-17::mCherry]* |
| FDU2324 | *cima-1(wy84)* IV; *mig-17(ola226)* V; *wyIs45 [Pttx-3::GFP::rab-3; Punc-122::RFP]* X; *shcEx1137 [Punc-14::mig-17; Phlh-17::mCherry]* |
| FDU2326 | *cima-1(wy84)* IV; *mig-17(ola226)* V; *wyIs45 [Pttx-3::GFP::rab-3; Punc-122::RFP]* X; *shcEx1139 [Phlh-17::mig-17; Phlh-17::mCherry]* |
| FDU2327 | *cima-1(wy84)* IV; *mig-17(ola226)* V; *wyIs45 [Pttx-3::GFP::rab-3; Punc-122::RFP]* X; *shcEx1140 [Phlh-17::mig-17; Phlh-17::mCherry]* |
| FDU2329 | *cima-1(wy84)* IV; *mig-17(ola226)* V; *wyIs45 [Pttx-3::GFP::rab-3; Punc-122::RFP]* X; *shcEx1142 [Pdpy-7::mig-17; Phlh-17::mCherry]* |

| | |
|---|---|
| FDU2330 | *cima-1(wy84)* IV*; mig-17(ola226)* V*; wyIs45 [Pttx-3::GFP::rab-3; Punc-122::RFP]* X*; shcEx1143 [Pdpy-7::mig-17; Phlh-17::mCherry]* |
| DCR1645 | *cima-1(wy84)* IV*; egl-15(n484)* X*; wyIs45 [Pttx-3::GFP::rab-3; Punc-122::RFP]* X |
| FDU942 | *cima-1(wy84)* IV*; mig-17(ola226)* V*; egl-15(n484)* X*; wyIs45 [Pttx-3::GFP::rab-3; Punc-122::RFP]* X |
| FDU38 | *cima-1(wy84)* IV*;fbl-1(k201)* IV*; wyIs45;[Pttx-3::GFP::rab-3; Punc-122::RFP]* X |
| FDU1518 | *cima-1(wy84)* IV*; emb-9(xd51)* III*; wyIs45 [Pttx-3::GFP::rab-3; Punc-122::RFP]* X |
| FDU2242 | *cima-1(wy84)* IV*; ost-1(gk786697)* IV*; wyIs45 [Pttx-3::GFP::rab-3; Punc-122::RFP]* X |
| FDU2292 | *cima-1(wy84)* IV*; nid-1(cg119)* V*; wyIs45 [Pttx-3::GFP::rab-3; Punc-122::RFP]* X |
| FDU1770 | *cima-1(wy84)* IV*; nid-1(cg118)* V*; wyIs45 [Pttx-3::GFP::rab-3; Punc-122::RFP]* X |
| FDU2156 | *cima-1(wy84)* IV*; cle-1(cg120)* I*; wyIs45 [Pttx-3::GFP::rab-3; Punc-122::RFP]* X |
| DCR1475 | *cima-1(wy84)* IV*; unc-52(e1421)* II*; wyIs45[Pttx-3::GFP::rab-3; Punc-122::RFP]* X |
| FDU192 | *cima-1(wy84)* IV*; emb-9(tk75)* III*; wyIs45 [Pttx-3::GFP::rab-3; Punc-122::RFP]* X |
| FDU1139 | *cima-1(wy84)* IV*;gon-1(q518)* IV*; wyIs45;[Pttx-3::GFP::rab-3; Punc-122::RFP]* X |
| FDU1471 | *cima-1(wy84)* IV*; let-2(b246)* X*; wyIs45 [Pttx-3::GFP::rab-3; Punc-122::RFP]* X |
| FDU45 | *cima-1(wy84)* IV*; mig-17(ola226)* V*; fbl-1(k201)* IV*; wyIs45 [Pttx-3::GFP::rab-3; Punc-122::RFP]* X |
| FDU914 | *cima-1(wy84)* IV*; mig-17(ola226)* V*; emb-9(tk75)* III*; wyIs45 [Pttx-3::GFP::rab-3; Punc-122::RFP]* X |
| FDU172 | *cima-1(wy84)* IV*; mig-17(ola226)* V*; let-2(k193)* X*; wyIs45 [Pttx-3::GFP::rab-3; Punc-122::RFP]* X |
| FDU2583 | *cima-1(wy84)* IV*; mig-17(ola226)* V*; let-2(b246)* X*; wyIs45 [Pttx-3::GFP::rab-3; Punc-122::RFP]* X |
| NK364 | *unc-119(ed4)* III*;qyIs46[unc119;emb-9::mCherry]* X |
| FDU2593 | *cima-1(wy84)* IV*; qyIs46[unc119;emb-9::mCherry]* X |
| FDU2594 | *mig-17(ola226)* V*;qyIs46[unc119;emb-9::mCherry]* X |
| FDU2595 | *cima-1(wy84)* IV*; mig-17(ola226)* V*; qyIs46[unc119;emb-9::mCherry]* X |
| FDU1140 | *shcEx776 [Phlh-17::mCherry; Pttx-3::GFP::rab-3]* |
| FDU1784 | *cima-1(wy84)* IV*; shcEx777 [Phlh-17::mCherry; Pttx-3::GFP::rab-3]* |
| FDU1785 | *mig-17(ola226)* V*; shcEx778 [Phlh-17::mCherry; Pttx-3::GFP::rab-3]* |
| FDU1143 | *cima-1(wy84)* IV*; mig-17(ola226)* V*; shcEx780 [Phlh-17::mCherry; Pttx-3::GFP::rab-3]* |
| FDU1144 | *cima-1(wy84)* IV*; let-2(k193)* X*; shcEx781 [Phlh-17::mCherry; Pttx-3::GFP::rab-3]* |
| FDU1025 | *shcEx424 [Pdpy-7::egl-15(5A); Phlh-17::mCherry; Pttx-3::GFP:: rab-3]* |
| FDU1222 | *shcEx536 [Pdpy-7::egl-15(5A); Phlh-17::mCherry; Pttx-3::GFP:: rab-3]* |
| FDU1026 | *mig-17(ola226)* V*; shcEx425 [Pdpy-7::egl-15(5A); Phlh-17::mCherry; Pttx-3::GFP:: rab-3]* |
| FDU1223 | *mig-17(ola226)* V*; shcEx537 [Pdpy-7::egl-15(5A); Phlh-17::mCherry; Pttx-3::GFP:: rab-3]* |
| FDU1224 | *mig-17(ola226)* V*; shcEx538 [Pdpy-7::egl-15(5A); Phlh-17::mCherry; Pttx-3::GFP:: rab-3]* |
| FDU178 | *mig-17(ola226) V; wyIs45 [Pttx-3::GFP::rab-3; Punc-122::RFP] X* |
| FDU883 | *mig-17(k113) V; wyIs45 [Pttx-3::GFP::rab-3; Punc-122::RFP] X* |
| FDU2583 | *wyIs45;shcEx1252[Pmig-17::mig-17 genomics; Phlh-17::mCherry]* |
| FDU2584 | *wyIs45;shcEx1253[Pmig-17::mig-17 genomics; Phlh-17::mCherry]* |

| | |
|---|---|
| FDU2247 | *ost-1(gk193565) IV;wyIs45 [Pttx-3::GFP::rab-3; Punc-122::RFP] X* |
| DCR2656 | *ost-1(gk786697) IV;wyIs45 [Pttx-3::GFP::rab-3; Punc-122::RFP] X* |
| FDU1769 | *nid-1(cg118) V;wyIs45 [Pttx-3::GFP::rab-3; Punc-122::RFP] X* |
| FDU2291 | *nid-1(cg119) V;wyIs45 [Pttx-3::GFP::rab-3; Punc-122::RFP] X* |
| DCR4032 | *fbl-1(k201) IV;wyIs45 [Pttx-3::GFP::rab-3; Punc-122::RFP] X* |
| FDU1470 | *let-2(b246) X;wyIs45 [Pttx-3::GFP::rab-3; Punc-122::RFP] X* |
| FDU2157 | *cle-1(cg120) I;wyIs45 [Pttx-3::GFP::rab-3; Punc-122::RFP] X* |
| FDU1519 | *emb-9(xd51) III; wyIs45 [Pttx-3::GFP::rab-3; Punc-122::RFP] X* |
