## Supplemental Table 2 for "ADAMTS-family protease MIG-17 regulates synaptic allometry by modifying the extracellular matrix and modulating glia morphology during growth"

**Table S2. Construct information**

| Construct name | Primers | Vector | Comments |
|---|---|---|---|
| *Pmig-17::SL2::GFP* | Forward: aaatatacATGAGTAAAggagaagaacttttcactggag<br>Reverse: TTACTCATgtatatttttcctttttcgcacacgttctacaac | pDEST | The Gateway system was described in (Norlia Basherudin et al., 2006). |
| *Pmig-17::mig-17* | | pDEST | SL2 and GFP were removed from *Pmig-17::SL2::GFP* |
| *mig-17(shc19)* Cas9-sgRNA1 | Forward: AATACTCGACTTCAAAATTAgttttagagctagaaatagcaagt<br>Reverse: TAATTTTGAAGTCGAGTATTcaagacatctcgcaatagg | pDD162 | pDD162 was described in (Daniel J Dickinson et al., 2015). This plasmid was constructed using the protocol of Dickinson 2015. |
| *mig-17(shc19)* Cas9-sgRNA2 | Forward: ACGTTCTACAACAAATCTCAgttttagagctagaaatagcaagt<br>Reverse: TGAGATTTGTTGTAGAACGTcaagacatctcgcaatagg | pDD162 | pDD162 was described in (Daniel J Dickinson et al., 2015). This plasmid was constructed using the protocol of Dickinson 2015. |
| *mig-17(shc19)* repair template of CRISPR/Ca | right and left homologous arm:<br>Forward: ttttGGCGCGCCTCTGACAATTCCTCAACGGATGC<br>Reverse: GACGCCTAGGgtaactaggcgggactcaagct<br>mutate NGG of sgRNA1:<br>Forward: AATTCAAAGACATGTTCtAAATACTCGACTTC<br>Reverse: GAACATGTCTTTGAATTAATATCTTTACAATCCATC<br>mutate NGG of sgRNA2:<br>Forward: TCGTGAGATTTGTTGTAGAACGTGTGCGAAAAAAG<br>Reverse: TTCTACAACAAATCTCACGAAAATCTTTCGATTTA<br>insert mNeonGreen:<br>Forward: gacgACCGGTATGGTGTCTAAGGGCGAAGA<br>Reverse: tccccccgggCTTGTACAGCTCGTCCATGC<br>Forward: tccccccgggTGAttggagtaataaacgtttcatatatat<br>Reverse: gacgACCGGTGTATATTTTTCCTTTTTTCGCACACGTTC<br>insert Flag:<br>Forward: cgcggatcctttcagGGAGCCGGATCTG<br>Reverse: tccccccgggTCTCTTGTCATCGTCATCCTTG<br>Forward: tccccccgggTGAttggagtaataaacgtttcatatatat<br>Reverse: cgcggatccCTTGTACAGCTCGTCCATGC<br>verification of knock-in animals:<br>Forward: GCTGGAACTTGGATCGCAAC<br>Reverse: GAAGTGGACCTGTTGAGGTT | pSM | *mig-17(shc19)* contains *mig-17::mNeonGreen::3×flag*. This plasmid was constructed by enzyme digest and T4 ligation. |
| *mig-17(shc8)* Cas9-sgRNA1 | Forward: AGGTCTCACTTCGTTGATTAgttttagagctagaaatagcaagt<br>Reverse: TAATCAACGAAGTGAGACCTcaagacatctcgcaatagg | pDD162 | pDD162 was described in (Daniel J Dickinson et al., 2015). This plasmid was constructed using the protocol of Dickinson 2015. |
| *mig-17(shc8)* Cas9-sgRNA2 | Forward: TTGATTATGGCTCACGAAATgttttagagctagaaatagcaagt<br>Reverse: ATTTCGTGAGCCATAATCAAcaagacatctcgcaatagg | pDD162 | pDD162 was described in (Daniel J Dickinson et al., 2015). This plasmid was constructed using the protocol of Dickinson 2015. |

| | | | |
|---|---|---|---|
| *mig-17(shc8)* repair template of CRISPR/Cas | Forward: ttttGGCGCGCCtttgggctgttaggagtagttc<br>Reverse: gacgACCGGTgttgctggaccacaaggaga<br>Forward: TTAATCATGGCCCATGCAATC GGTCACTCgtgagttaaact<br>Reverse: ATGGGCCATGATTAATGAGGTAAGACCTGCTCCAATGTCCTCTA<br>verifiction of mutant animals:<br>Forward: GACGTGAGAAGCAACAAAGC<br>Reverse: GAAGTGGACCTGTTGAGGTT | pSM | *mig-17(shc8)* contains *mig-17(E303A)* where the protease catalytic site of MIG-17 is altered from E303 to A. This plasmid was constructed by enzyme digest and T4 ligation. |
| *Pmig-17::mig-17(E303A)::GFP* | | pSM | A gift from the lab of Kiyoji Nishiwaki |
| *Phlh-17::mig-17* | Forward: TACCCATCCGCAGACACTG<br>Reverse: AAGCTCCTGAGTCTTGTTCG | pDEST | *Phlh-17* is published in (McMiller and Johnson, 2005). The Gateway system was described in (Norlia Basherudin et al., 2006). |
| *Punc-14::mig-17* | | pDEST | *Punc-14* is published in (Ogura et al., 1997). The Gateway system was described in (Norlia Basherudin et al., 2006). |
| *Ppdy-7::mig-17* | | pDEST | *Pdpy-7* is published in (Bülow et al., 2004). The Gateway system was described in (Norlia Basherudin et al., 2006). |
| *Pmyo-3::mig-17* | | pDEST | *Pmyo-3* was described in (Okkema et al., 1993). The Gateway system was described in (Norlia Basherudin et al., 2006). |
| *dgn-1 RNAi* | Forward: atttgcggccgcAACCCTCAAGTTTACACTGT<br>Reverse: atttgcggccgcAGTGAGAATGACCATTGACC | pPD129.36 | The full length of *dgn-1 cDNA* . |
| *Pdpy-7::egl-15(5A)* | | pDEST | P*dpy-7* is published in (Bülow et al., 2004).The Gateway system was described in (Norlia Basherudin et al., 2006). |
| *Prab-3::mCherry* | | | *Prab-3* was described in (Nonet et al., 1997). From the lab of Daniel A. Colón-Ramos. |
| *Pttx-3g::GFP::rab-3* | | | *Pttx-3g* was described in (Wenick and Hobert, 2004). From the lab of Daniel A. Colón-Ramos. |
| *Phlh-17::mCherry* | | | *Phlh-17* is published in (McMiller and Johnson, 2005). From the lab of Daniel A. Colón-Ramos. |
| *Pdpy-4::mCherry* | | | From the lab of Daniel A. Colón-Ramos. |
