## Supplemental Table 3 for "ADAMTS-family protease MIG-17 regulates synaptic allometry by modifying the extracellular matrix and modulating glia morphology during growth"

**Table S3. The results of MS**

| Accession | Gene names | N2 repeat 1 | N2 repeat 2 | N2 repeat 3 | average | mig-17(ola226) repeat 1 | mig-17(ola226) repeat 2 | mig-17(ola226) repeat 3 | average | mig-17(ola226)/N2 |
|---|---|---|---|---|---|---|---|---|---|---|
| Q20877 | CELE_F56D3.1 F56D3.1 | 0.81 | 0.564 | 0 | 0.458 | 0.19 | 0 | 0 | 0.063333333 | 0.138282387 |
| P30629 | ZK637.2 | 1 | 1 | 0.357 | 0.785666667 | 0 | 0 | 0.495 | 0.165 | 0.210012728 |
| Q20049 | idhg-1 CELE_F35G12.2 F35G12.2 | 0.424 | 1 | 0 | 0.474666667 | 0.412 | 0 | 0 | 0.137333333 | 0.289325843 |
| Q21742 | CELE_R05F9.6 R05F9.6 | 0.327 | 0 | 1 | 0.442333333 | 0.426 | 0 | 0 | 0.142 | 0.321024868 |
| Q9XW20 | CELE_Y18D10A.11 Y18D10A.11 | 0.354 | 0 | 1 | 0.451333333 | 0.435 | 0 | 0 | 0.145 | 0.32127031 |
| Q09482 | C18H9.3/C18H9.2 | 0 | 1 | 0.328 | 0.442666667 | 0 | 0 | 0.444 | 0.148 | 0.334337349 |
| Q2EEM8 | ttr-45 CELE_JC8.14 JC8.14 | 0.272 | 0.466 | 0.563 | 0.433666667 | 0.48 | 0 | 0 | 0.16 | 0.368946964 |
| Q20223 | lbp-1 F40F4.3 | 0 | 0.334 | 0.57 | 0.301333333 | 0 | 0.375 | 0 | 0.125 | 0.414823009 |
| O44144 | perm-4 C44B12.5 CELE_C44B12.5 | 0.34 | 0.499 | 1 | 0.613 | 0.379 | 0.388 | 0 | 0.255666667 | 0.417074497 |
| Q22015 | CELE_R31.2 R31.2 | 0 | 0 | 0.512 | 0.170666667 | 0 | 0 | 0.214 | 0.071333333 | 0.41796875 |
| Q9GUF2 | acp-6 CELE_Y73B6BL.24 Y73B6BL.24 | 0 | 0.337 | 0.501 | 0.279333333 | 0 | 0.384 | 0 | 0.128 | 0.45823389 |
| G5EGA7 | ras-2 Ras2 CELE_F17C8.4 F17C8.4 | 0 | 0.247 | 0 | 0.082333333 | 0 | 0.114 | 0 | 0.038 | 0.461538462 |
| O45006 | CELE_W03D8.9 W03D8.9 | 1 | 0 | 0.183 | 0.394333333 | 0 | 0 | 0.55 | 0.183333333 | 0.464919696 |
| P34654 | ZK632.9 | 0 | 0.513 | 0 | 0.171 | 0 | 0.243 | 0 | 0.081 | 0.473684211 |
| Q9Y0V6 | tin-10 tim-10 Y66D12A.22 | 0 | 0 | 0.675 | 0.225 | 0 | 0 | 0.325 | 0.108333333 | 0.481481481 |
| Q9XWU9 | CELE_Y37D8A.19 Y37D8A.19 | 0.501 | 0 | 0.411 | 0.304 | 0.203 | 0 | 0.237 | 0.146666667 | 0.48245614 |
| O76618 | cars-1 CELE_Y23H5A.7 Y23H5A.7 | 0 | 0 | 0.489 | 0.163 | 0 | 0 | 0.239 | 0.079666667 | 0.488752556 |
| Q9UB28 | let-805 CELE_H19M22.2 H19M22.2 | 0.28 | 0 | 0.59 | 0.29 | 0.452 | 0 | 0 | 0.150666667 | 0.51954023 |
| Q966D6 | mct-2 mct-1 C01B4.9 CELE_C01B4.9 CELE_Y19D10A.12 Y19D10A.12 | 0 | 1 | 0 | 0.333333333 | 0 | 0.532 | 0 | 0.177333333 | 0.532 |
| P91340 | CELE_F55F8.2 F55F8.2 | 0 | 0 | 0.552 | 0.184 | 0 | 0 | 0.295 | 0.098333333 | 0.53442029 |
| Q22782 | rab-6.2 T25G12.4 | 0 | 0.638 | 0 | 0.212666667 | 0 | 0.362 | 0 | 0.120666667 | 0.567398119 |
| P91280 | CELE_F27C1.6 F27C1.6 | 0 | 0 | 0.554 | 0.184666667 | 0 | 0 | 0.315 | 0.105 | 0.568592058 |
| Q18032 | C15H9.9 CELE_C15H9.9 | 0.28 | 1 | 0.305 | 0.528333333 | 0.468 | 0 | 0.438 | 0.302 | 0.571608833 |
| O44954 | sdha-2 C34B2.7 CELE_C34B2.7 | 0.496 | 0 | 0 | 0.165333333 | 0.294 | 0 | 0 | 0.098 | 0.592741935 |
| O44572 | tni-4 W03F8.1 | 0 | 0 | 0.624 | 0.208 | 0 | 0 | 0.376 | 0.125333333 | 0.602564103 |
| Q20277 | fipr-21 CELE_F41E7.5 F41E7.5 | 0.349 | 0.509 | 0.335 | 0.397666667 | 0.37 | 0 | 0.368 | 0.246 | 0.61860855 |
| G5EFB5 | cpn-1 CELE_F43G9.9 F43G9.9 | 0.502 | 0.459 | 0.311 | 0.424 | 0.253 | 0.151 | 0.399 | 0.267666667 | 0.631289308 |
| Q17435 | pfd-4 tag-317 B0035.4 | 0.546 | 0.324 | 0 | 0.29 | 0 | 0.555 | 0 | 0.185 | 0.637931034 |
| Q9XWK2 | CELE_Y54E5A.5 Y54E5A.5 | 0.46 | 0.475 | 0.777 | 0.570666667 | 0.368 | 0.345 | 0.397 | 0.37 | 0.648364486 |
| Q21322 | rnp-2 CELE_K08D10.4 K08D10.4 | 0 | 0.301 | 0 | 0.100333333 | 0 | 0.196 | 0 | 0.065333333 | 0.651162791 |
| O45430 | mccc-1 CELE_F32B6.2 F32B6.2 | 0.311 | 0.312 | 0 | 0.207666667 | 0 | 0.416 | 0 | 0.138666667 | 0.667736758 |

| | | | | | | | | | | |
|---|---|---|---|---|---|---|---|---|---|---|
| O44145 | perm-2 C44B12.1 CELE_C44B12.1 | 0.281 | 0.157 | 0.653 | 0.363666667 | 0.337 | 0.414 | 0 | 0.250333333 | 0.688359303 |
| Q9GZH3 | T22D1.3 | 0.146 | 1 | 0.413 | 0.519666667 | 0.584 | 0 | 0.518 | 0.367333333 | 0.706863374 |
| Q9XWT3 | ule-5 CELE_Y62H9A.6 Y62H9A.6 | 0.36 | 0.553 | 0 | 0.304333333 | 0.393 | 0.255 | 0 | 0.216 | 0.709748083 |
| Q94148 | rab-10 T23H2.5 | 0 | 0.581 | 0 | 0.193666667 | 0 | 0.419 | 0 | 0.139666667 | 0.721170396 |
| O17891 | CELE_F55B11.2 F55B11.2 | 0 | 0 | 0.453 | 0.151 | 0 | 0 | 0.327 | 0.109 | 0.721854305 |
| Q23028 | CELE_R09F10.5 R09F10.5 | 0.596 | 0.455 | 0.456 | 0.502333333 | 0.404 | 0.334 | 0.369 | 0.369 | 0.734571997 |
| Q9XXK7 | tram-1 C24F3.1 CELE_C24F3.1 | 0 | 0.447 | 0 | 0.149 | 0 | 0.338 | 0 | 0.112666667 | 0.756152125 |
| P41847 | T20B12.7 | 0.526 | 0.437 | 0 | 0.321 | 0.351 | 0.387 | 0 | 0.246 | 0.76635514 |
| P34640 | ZK512.2 | 0 | 0.566 | 0 | 0.188666667 | 0 | 0.434 | 0 | 0.144666667 | 0.766784452 |
| G5EET8 | pud-1.2 pud-1.1 CELE_F15E11.13 CELE_Y19D10B.7 F15E11.13 Y19D10B.7 | 0.36 | 0.349 | 0.367 | 0.358666667 | 0.272 | 0.293 | 0.264 | 0.276333333 | 0.770446097 |
| Q9BKU8 | moag-4 CELE_Y37E3.4 Y37E3.4 | 0 | 0 | 0.434 | 0.144666667 | 0 | 0 | 0.337 | 0.112333333 | 0.776497696 |
| Q94053 | msp-78 T13F2.11 | 0.551 | 0.567 | 0.312 | 0.476666667 | 0.397 | 0.361 | 0.361 | 0.373 | 0.782517483 |
| Q21774 | adsl-1 R06C7.5 | 0.411 | 0 | 0 | 0.137 | 0.323 | 0 | 0 | 0.107666667 | 0.785888078 |
| Q9TYV5 | nol-1 CELE_W07E6.1 W07E6.1 | 0 | 0.478 | 0.45 | 0.309333333 | 0 | 0.73 | 0 | 0.243333333 | 0.786637931 |
| Q9N4I3 | CELE_Y71F9AL.9 Y71F9AL.9 | 0.414 | 0.378 | 0.316 | 0.369333333 | 0.44 | 0.498 | 0.392 | 0.443333333 | 1.200361011 |
| Q9NAR3 | col-124 C24F3.6 CELE_C24F3.6 | 0.309 | 0.341 | 0.348 | 0.332666667 | 0.256 | 0.488 | 0.455 | 0.399666667 | 1.201402806 |
| Q9N4N4 | swsn-6 CELE_ZK616.4 ZK616.4 | 0.362 | 0 | 0 | 0.120666667 | 0.435 | 0 | 0 | 0.145 | 1.201657459 |
| Q9TZ33 | ucr-2.3 CELE_T24C4.1 T24C4.1 | 0 | 0.323 | 0.381 | 0.234666667 | 0 | 0.431 | 0.415 | 0.282 | 1.201704545 |
| P19625 | mlc-1 C36E6.3 | 0 | 0 | 0.454 | 0.151333333 | 0 | 0 | 0.548 | 0.182666667 | 1.207048458 |
| Q9XVS1 | tag-72 C25A1.3 | 0 | 0 | 0.453 | 0.151 | 0 | 0 | 0.547 | 0.182333333 | 1.207505519 |
| Q9XXE2 | CELE_Y44A6D.2 Y44A6D.2 | 0 | 0.366 | 0 | 0.122 | 0 | 0.443 | 0 | 0.147666667 | 1.210382514 |
| P34685 | cap-1 D2024.6 | 0.294 | 0 | 0.365 | 0.219666667 | 0.452 | 0 | 0.346 | 0.266 | 1.210925645 |
| O62053 | C08F11.11 | 0.334 | 0.34 | 0.323 | 0.332333333 | 0.417 | 0.394 | 0.398 | 0.403 | 1.212637914 |
| H2KZV8 | mlp-1 CELE_T04C9.4 T04C9.4 | 0.349 | 0.344 | 0.328 | 0.340333333 | 0.439 | 0.39 | 0.411 | 0.413333333 | 1.214495593 |
| Q20588 | F49C12.11 | 0 | 0.292 | 0 | 0.097333333 | 0 | 0.355 | 0 | 0.118333333 | 1.215753425 |
| O01805 | acbp-1 C44E4.6 | 0.321 | 0.389 | 0.344 | 0.351333333 | 0.44 | 0.416 | 0.426 | 0.427333333 | 1.216318786 |
| Q27481 | uba-1 C47E12.5 CELE_C47E12.5 | 0.353 | 0.325 | 0.385 | 0.354333333 | 0.404 | 0.469 | 0.42 | 0.431 | 1.216368768 |
| P91303 | vha-10 F46F11.5 | 0.336 | 0.35 | 0.336 | 0.340666667 | 0.418 | 0.411 | 0.415 | 0.414666667 | 1.217221135 |
| Q93568 | CELE_F25H2.4 F25H2.4 | 0 | 0 | 0.451 | 0.150333333 | 0 | 0 | 0.549 | 0.183 | 1.2172949 |
| Q9N2W7 | Y94H6A.8 | 0 | 0 | 0.338 | 0.112666667 | 0 | 0 | 0.412 | 0.137333333 | 1.218934911 |
| Q95PZ1 | CELE_Y67H2A.5 Y67H2A.5 | 0.305 | 0.39 | 0.384 | 0.359666667 | 0.461 | 0.426 | 0.43 | 0.439 | 1.220574606 |
| Q21962 | CELE_R12C12.1 R12C12.1 | 0.382 | 0.258 | 0 | 0.213333333 | 0.392 | 0.39 | 0 | 0.260666667 | 1.221875 |
| Q9NLD1 | hrp-2 CELE_F58D5.1 F58D5.1 | 0.326 | 0.32 | 0.381 | 0.342333333 | 0.429 | 0.433 | 0.396 | 0.419333333 | 1.224926972 |

| | | | | | | | | | | |
|---|---|---|---|---|---|---|---|---|---|---|
| Q95017 | ubc-9 F29B9.6 | 0 | 0.319 | 0 | 0.106333333 | 0 | 0.391 | 0 | 0.130333333 | 1.225705329 |
| Q86NE1 | asp-2 CELE_T18H9.2 T18H9.2 | 0.322 | 0.355 | 0.348 | 0.341666667 | 0.423 | 0.418 | 0.418 | 0.419666667 | 1.228292683 |
| P34339 | egl-45 eif-3.A C27D11.1 | 0.281 | 0.317 | 0.399 | 0.332333333 | 0.401 | 0.411 | 0.413 | 0.408333333 | 1.228686058 |
| Q95YB2 | acdh-9 CELE_F28A10.6 F28A10.6 | 0 | 0.469 | 0.29 | 0.253 | 0 | 0.474 | 0.459 | 0.311 | 1.229249012 |
| Q94010 | CELE_T08G11.1 T08G11.1 | 0.448 | 0.528 | 0.369 | 0.448333333 | 0.552 | 0.472 | 0.631 | 0.551666667 | 1.230483271 |
| O45569 | nep-17 CELE_F54F11.2 F54F11.2 | 0 | 0 | 0.336 | 0.112 | 0 | 0 | 0.414 | 0.138 | 1.232142857 |
| Q17698 | C06A8.3 CELE_C06A8.3 | 0.312 | 0.318 | 0.316 | 0.315333333 | 0.391 | 0.384 | 0.391 | 0.388666667 | 1.23255814 |
| G5ECW7 | dpt-1 CELE_F02E9.9 F02E9.9 | 0 | 0 | 0.406 | 0.135333333 | 0 | 0 | 0.501 | 0.167 | 1.233990148 |
| P18835 | col-19 ZK1193.1 | 0.293 | 0 | 0 | 0.097666667 | 0.362 | 0 | 0 | 0.120666667 | 1.235494881 |
| Q19766 | tomm-20 F23H12.2 | 0.303 | 0.287 | 0.343 | 0.311 | 0.381 | 0.427 | 0.346 | 0.384666667 | 1.236870311 |
| O17328 | mct-6 C10E2.6 CELE_C10E2.6 | 0.368 | 0.365 | 0 | 0.244333333 | 0.505 | 0.403 | 0 | 0.302666667 | 1.238744884 |
| P55956 | asp-3 H22K11.1 | 0.319 | 0.347 | 0.342 | 0.336 | 0.419 | 0.417 | 0.415 | 0.417 | 1.241071429 |
| P91027 | chdp-1 C10G11.7 CELE_C10G11.7 | 0.302 | 0.419 | 0.313 | 0.344666667 | 0.46 | 0.327 | 0.497 | 0.428 | 1.241779497 |
| Q09936 | C53C9.2 | 0.322 | 0.318 | 0.326 | 0.322 | 0.41 | 0.384 | 0.408 | 0.400666667 | 1.244306418 |
| Q22352 | CELE_T08H10.1 T08H10.1 | 0.328 | 0.359 | 0.328 | 0.338333333 | 0.457 | 0.368 | 0.438 | 0.421 | 1.244334975 |
| P31161 | sod-2 sdm-1 F10D11.1 | 0.322 | 0.308 | 0.45 | 0.36 | 0.438 | 0.456 | 0.45 | 0.448 | 1.244444444 |
| Q9N408 | ddp-1 tim-8 Y39A3CR.4 | 0.432 | 0.265 | 0.436 | 0.377666667 | 0.568 | 0.43 | 0.412 | 0.47 | 1.244483672 |
| O02115 | pcn-1 W03D2.4 | 0.367 | 0.318 | 0.242 | 0.309 | 0.335 | 0.404 | 0.416 | 0.385 | 1.245954693 |
| G3MU83 | math-33 CELE_H19N07.2 H19N07.2 | 0.318 | 0.318 | 0.365 | 0.333666667 | 0.431 | 0.419 | 0.401 | 0.417 | 1.24975025 |
| P55955 | ttr-16 Y5F2A.1 | 0.538 | 0.275 | 0.3 | 0.371 | 0.517 | 0.444 | 0.43 | 0.463666667 | 1.249775382 |
| Q9U296 | men-1 CELE_Y48B6A.12 Y48B6A.12 | 0 | 0 | 0.375 | 0.125 | 0 | 0 | 0.469 | 0.156333333 | 1.250666667 |
| Q20774 | dnj-13 CELE_F54D5.8 F54D5.8 | 0 | 0 | 0.342 | 0.114 | 0 | 0 | 0.428 | 0.142666667 | 1.251461988 |
| P91306 | cey-2 CELE_F46F11.2 F46F11.2 | 0.329 | 0.347 | 0.322 | 0.332666667 | 0.411 | 0.434 | 0.405 | 0.416666667 | 1.25250501 |
| Q19826 | rpb-8 F26F4.11 | 0.35 | 0 | 0 | 0.116666667 | 0.439 | 0 | 0 | 0.146333333 | 1.254285714 |
| Q9XXS2 | set-22 CELE_Y32F6A.1 Y32F6A.1 | 0 | 0 | 0.35 | 0.116666667 | 0 | 0 | 0.439 | 0.146333333 | 1.254285714 |
| O02323 | lbp-7 T22G5.2 | 0.319 | 0 | 0.312 | 0.210333333 | 0.432 | 0 | 0.36 | 0.264 | 1.255150555 |
| Q17820 | aos-1 C08B6.9 | 0 | 0.329 | 0 | 0.109666667 | 0 | 0.413 | 0 | 0.137666667 | 1.255319149 |
| O44955 | C34B2.8 CELE_C34B2.8 | 0 | 0.321 | 0 | 0.107 | 0 | 0.403 | 0 | 0.134333333 | 1.255451713 |
| Q21217 | gta-1 K04D7.3 | 0.337 | 0.311 | 0.33 | 0.326 | 0.407 | 0.403 | 0.42 | 0.41 | 1.257668712 |
| O17607 | ruvb-1 C27H6.2 | 0.301 | 0.358 | 0.337 | 0.332 | 0.434 | 0.396 | 0.423 | 0.417666667 | 1.258032129 |
| Q27535 | ZC434.8 | 0.316 | 0.341 | 0.352 | 0.336333333 | 0.403 | 0.442 | 0.425 | 0.423333333 | 1.258671952 |
| Q19842 | pcca-1 F27D9.5 | 0.332 | 0.419 | 0.284 | 0.345 | 0.474 | 0.36 | 0.469 | 0.434333333 | 1.258937198 |
| Q20107 | fkb-1 CELE_F36H1.1 F36H1.1 | 0.351 | 0 | 0 | 0.117 | 0.442 | 0 | 0 | 0.147333333 | 1.259259259 |
| Q20310 | CELE_F42A10.5 F42A10.5 | 0.386 | 0 | 0.288 | 0.224666667 | 0.423 | 0 | 0.426 | 0.283 | 1.259643917 |

| | | | | | | | | | | |
|---|---|---|---|---|---|---|---|---|---|---|
| Q22235 | enpl-1 T05E11.3 | 0.313 | 0.364 | 0.339 | 0.338666667 | 0.413 | 0.416 | 0.453 | 0.427333333 | 1.261811024 |
| P34388 | mrps-9 F09G8.3 | 0 | 0.038 | 0 | 0.012666667 | 0 | 0.048 | 0 | 0.016 | 1.263157895 |
| Q93714 | idha-1 F43G9.1 | 0.332 | 0.336 | 0.35 | 0.339333333 | 0.436 | 0.424 | 0.426 | 0.428666667 | 1.263261297 |
| O62388 | CELE_W01D2.1 W01D2.1 | 0.341 | 0.315 | 0.325 | 0.327 | 0.41 | 0.422 | 0.409 | 0.413666667 | 1.265035678 |
| Q2XN18 | spp-14 CELE_K09F5.3 K09F5.3 | 0.345 | 0.309 | 0.306 | 0.32 | 0.352 | 0.426 | 0.437 | 0.405 | 1.265625 |
| Q21763 | CELE_R05H5.3 R05H5.3 | 0.35 | 0.284 | 0.378 | 0.337333333 | 0.439 | 0.446 | 0.399 | 0.428 | 1.268774704 |
| Q21000 | myo-5 CELE_F58G4.1 F58G4.1 | 0.345 | 0.298 | 0.513 | 0.385333333 | 0.535 | 0.47 | 0.462 | 0.489 | 1.269031142 |
| H2FLH2 | unc-22 CELE_ZK617.1 ZK617.1 | 0.253 | 0.369 | 0.396 | 0.339333333 | 0.434 | 0.413 | 0.445 | 0.430666667 | 1.269155206 |
| Q9XU97 | CELE_F44E5.1 F44E5.1 | 0.367 | 0.322 | 0.284 | 0.324333333 | 0.356 | 0.427 | 0.453 | 0.412 | 1.270298047 |
| Q18599 | acer-1 C44B7.10 CELE_C44B7.10 | 0.326 | 0.293 | 0.333 | 0.317333333 | 0.408 | 0.423 | 0.38 | 0.403666667 | 1.272058824 |
| Q22037 | hrp-1 rbp-1 F42A6.7 | 0.336 | 0.352 | 0.363 | 0.350333333 | 0.429 | 0.489 | 0.42 | 0.446 | 1.273073264 |
| O45924 | Y39E4A.3 | 0.32 | 0.328 | 0.318 | 0.322 | 0.428 | 0.428 | 0.374 | 0.41 | 1.273291925 |
| Q22781 | acdh-7 CELE_T25G12.5 T25G12.5 | 0.362 | 0.362 | 0.289 | 0.337666667 | 0.41 | 0.433 | 0.447 | 0.43 | 1.273445212 |
| Q23445 | ZK180.4 | 0.328 | 0 | 0 | 0.109333333 | 0.418 | 0 | 0 | 0.139333333 | 1.274390244 |
| Q19264 | F09E5.3 | 0.333 | 0 | 0.338 | 0.223666667 | 0.44 | 0 | 0.416 | 0.285333333 | 1.275707899 |
| Q09359 | ZK1307.1 | 0.348 | 0.337 | 0.322 | 0.335666667 | 0.425 | 0.424 | 0.436 | 0.428333333 | 1.276067527 |
| Q9XXU9 | vha-11 Y38F2AL.3 | 0.385 | 0.342 | 0.323 | 0.35 | 0.455 | 0.43 | 0.455 | 0.446666667 | 1.276190476 |
| P34462 | vha-14 F55H2.2 | 0.311 | 0.468 | 0.3 | 0.359666667 | 0.42 | 0.532 | 0.428 | 0.46 | 1.278962002 |
| P52009 | cyn-1 cyp-1 Y49A3A.5 | 0.376 | 0.296 | 0.303 | 0.325 | 0.422 | 0.425 | 0.401 | 0.416 | 1.28 |
| G5ED41 | cand-1 Y102A5A.1 | 0 | 0 | 0.3 | 0.1 | 0 | 0 | 0.384 | 0.128 | 1.28 |
| O01602 | nuo-2 CELE_T10E9.7 T10E9.7 | 0 | 0 | 0.324 | 0.108 | 0 | 0 | 0.415 | 0.138333333 | 1.280864198 |
| Q23280 | daf-41 p23 ZC395.10 | 0.281 | 0.51 | 0.276 | 0.355666667 | 0.461 | 0.448 | 0.459 | 0.456 | 1.282099344 |
| Q10453 | his-71 F45E1.6 | 0.279 | 0.518 | 0.29 | 0.362333333 | 0.477 | 0.479 | 0.439 | 0.465 | 1.283348666 |
| O62289 | ttr-51 CELE_JC8.8 JC8.8 | 0.336 | 0.311 | 0.33 | 0.325666667 | 0.408 | 0.425 | 0.421 | 0.418 | 1.283520983 |
| P53596 | sucl-1 C05G5.4 | 0.358 | 0.322 | 0.342 | 0.340666667 | 0.407 | 0.473 | 0.433 | 0.437666667 | 1.284735812 |
| Q09508 | sdha-1 C03G5.1/D2021.3 | 0.35 | 0.411 | 0.338 | 0.366333333 | 0.517 | 0.436 | 0.459 | 0.470666667 | 1.284804368 |
| Q20239 | aagr-3 CELE_F40F9.6 F40F9.6 | 0 | 0.437 | 0 | 0.145666667 | 0 | 0.563 | 0 | 0.187666667 | 1.288329519 |
| Q93353 | idhb-1 C37E2.1 | 0.348 | 0.362 | 0.307 | 0.339 | 0.43 | 0.472 | 0.409 | 0.437 | 1.289085546 |
| Q20922 | col-159 CELE_F57B1.3 F57B1.3 | 0 | 0.293 | 0 | 0.097666667 | 0 | 0.378 | 0 | 0.126 | 1.290102389 |
| P90901 | ifa-1 F38B2.1 | 0.346 | 0.356 | 0.33 | 0.344 | 0.521 | 0.363 | 0.449 | 0.444333333 | 1.291666667 |
| Q9U2S6 | cyn-13 CELE_Y116A8C.34 Y116A8C.34 | 0.333 | 0.376 | 0.411 | 0.373333333 | 0.416 | 0.442 | 0.589 | 0.482333333 | 1.291964286 |
| Q23098 | nuo-6 CELE_W01A8.4 W01A8.4 | 0.264 | 0.268 | 0.481 | 0.337666667 | 0.47 | 0.431 | 0.408 | 0.436333333 | 1.292201382 |
| Q03565 | baf-1 B0464.7 | 0.319 | 0.337 | 0.318 | 0.324666667 | 0.441 | 0.419 | 0.399 | 0.419666667 | 1.292607803 |
| O61793 | CELE_R12E2.13 R12E2.13 | 0.394 | 0 | 0.29 | 0.228 | 0.391 | 0 | 0.494 | 0.295 | 1.293859649 |

| | | | | | | | | | | |
|---|---|---|---|---|---|---|---|---|---|---|
| O01615 | T19H12.2 | 0.492 | 0.263 | 0.334 | 0.363 | 0.298 | 0.447 | 0.666 | 0.470333333 | 1.295684114 |
| O01532 | asp-5 CELE_F21F8.3 F21F8.3 | 0.305 | 0.336 | 0.333 | 0.324666667 | 0.416 | 0.442 | 0.404 | 0.420666667 | 1.295687885 |
| Q09EE7 | nsf-1 CELE_H15N14.2 H15N14.2 | 0.34 | 0.33 | 0.355 | 0.341666667 | 0.444 | 0.461 | 0.425 | 0.443333333 | 1.297560976 |
| Q27245 | lap-2 W07G4.4 | 0.342 | 0.31 | 0.34 | 0.330666667 | 0.437 | 0.438 | 0.413 | 0.429333333 | 1.298387097 |
| Q9TZL8 | pfk-1 Y71H10A.1 | 0 | 0.364 | 0.335 | 0.233 | 0 | 0.439 | 0.469 | 0.302666667 | 1.298998569 |
| H2L2E8 | tba-1 CELE_F26E4.8 F26E4.8 | 0.311 | 0 | 0 | 0.103666667 | 0.404 | 0 | 0 | 0.134666667 | 1.29903537 |
| Q9TZS5 | cct-7 CELE_T10B5.5 T10B5.5 | 0.615 | 0.274 | 0.261 | 0.383333333 | 0.57 | 0.471 | 0.457 | 0.499333333 | 1.302608696 |
| Q19832 | msp-3 F26G1.7 | 0.346 | 0 | 0.344 | 0.23 | 0.441 | 0 | 0.458 | 0.299666667 | 1.302898551 |
| Q9N384 | lec-6 CELE_Y55B1AR.1 Y55B1AR.1 | 0.32 | 0.376 | 0.363 | 0.353 | 0.464 | 0.458 | 0.458 | 0.46 | 1.303116147 |
| O17641 | col-178 C34F6.2 CELE_C34F6.2 | 0 | 0.291 | 0.295 | 0.195333333 | 0 | 0.452 | 0.312 | 0.254666667 | 1.303754266 |
| G4SI07 | gei-15 CELE_M03A8.4 M03A8.4 | 0 | 0.384 | 0 | 0.128 | 0 | 0.501 | 0 | 0.167 | 1.3046875 |
| Q20964 | lys-4 CELE_F58B3.1 F58B3.1 | 0.308 | 0.332 | 0.292 | 0.310666667 | 0.417 | 0.385 | 0.414 | 0.405333333 | 1.30472103 |
| Q3LFN1 | lbp-9 CELE_Y40B10A.1 Y40B10A.1 | 0.309 | 0.317 | 0.314 | 0.313333333 | 0.405 | 0.397 | 0.427 | 0.409666667 | 1.307446809 |
| Q21230 | K04G2.1 | 0.364 | 0.316 | 0.295 | 0.325 | 0.407 | 0.433 | 0.435 | 0.425 | 1.307692308 |
| P30642 | eif-3.D R08D7.3 | 0.298 | 0.392 | 0.286 | 0.325333333 | 0.43 | 0.424 | 0.424 | 0.426 | 1.30942623 |
| O44727 | cpn-4 CELE_F49D11.8 F49D11.8 | 0.33 | 0.353 | 0.32 | 0.334333333 | 0.428 | 0.453 | 0.436 | 0.439 | 1.313060818 |
| Q17512 | B0491.5 CELE_B0491.5 | 0.402 | 0.286 | 0.487 | 0.391666667 | 0.508 | 0.466 | 0.569 | 0.514333333 | 1.313191489 |
| Q95XR0 | CELE_Y39G10AR.8 Y39G10AR.8 | 0.374 | 0.279 | 0.351 | 0.334666667 | 0.485 | 0.452 | 0.383 | 0.44 | 1.314741036 |
| O44512 | isp-1 CELE_F42G8.12 F42G8.12 | 0.321 | 0.334 | 0.342 | 0.332333333 | 0.431 | 0.428 | 0.452 | 0.437 | 1.314944835 |
| Q9U2Q8 | fkb-2 CELE_Y18D10A.19 Y18D10A.19 | 0.325 | 0.317 | 0.315 | 0.319 | 0.42 | 0.409 | 0.431 | 0.42 | 1.31661442 |
| P34697 | sod-1 C15F1.7 | 0.329 | 0.315 | 0.323 | 0.322333333 | 0.433 | 0.425 | 0.416 | 0.424666667 | 1.317476732 |
| G5EDD4 | tba-4 CELE_F44F4.11 F44F4.11 | 0.329 | 0.323 | 0.282 | 0.311333333 | 0.414 | 0.419 | 0.399 | 0.410666667 | 1.319057816 |
| P19974 | cyc-2.1 E04A4.7 | 0.33 | 0.339 | 0.339 | 0.336 | 0.446 | 0.442 | 0.442 | 0.443333333 | 1.319444444 |
| Q9BL34 | CELE_Y71H2AM.5 Y71H2AM.5 | 0.315 | 0.305 | 0.33 | 0.316666667 | 0.42 | 0.418 | 0.416 | 0.418 | 1.32 |
| Q18231 | rps-30 C26F1.4 CELE_C26F1.4 | 0.326 | 0.319 | 0.319 | 0.321333333 | 0.388 | 0.394 | 0.491 | 0.424333333 | 1.320539419 |
| G5EBH7 | calu-1 CELE_M03F4.7 M03F4.7 | 0.365 | 0.32 | 0.378 | 0.354333333 | 0.432 | 0.53 | 0.443 | 0.468333333 | 1.32173095 |
| Q21930 | rpl-28 R11D1.8 | 0.334 | 0.323 | 0.327 | 0.328 | 0.434 | 0.432 | 0.435 | 0.433666667 | 1.322154472 |
| Q20363 | sip-1 F43D9.4 | 0.319 | 0.343 | 0.321 | 0.327666667 | 0.432 | 0.44 | 0.428 | 0.433333333 | 1.322482197 |
| Q95XT5 | trap-1 CELE_Y71F9AM.6 Y71F9AM.6 | 0.319 | 0.326 | 0.319 | 0.321333333 | 0.423 | 0.434 | 0.419 | 0.425333333 | 1.323651452 |
| Q9XWI6 | eif-3.B Y54E2A.11 | 0.315 | 0.341 | 0.319 | 0.325 | 0.422 | 0.41 | 0.459 | 0.430333333 | 1.324102564 |
| Q18211 | ran-3 C26D10.1 | 0.413 | 0.325 | 0.277 | 0.338333333 | 0.459 | 0.407 | 0.478 | 0.448 | 1.324137931 |
| P90983 | rps-29 B0412.4 CELE_B0412.4 | 0.342 | 0.274 | 0.268 | 0.294666667 | 0.386 | 0.392 | 0.394 | 0.390666667 | 1.325791855 |

| | | | | | | | | | | |
|---|---|---|---|---|---|---|---|---|---|---|
| Q10020 | T28D9.1 | 0.313 | 0.361 | 0.312 | 0.328666667 | 0.39 | 0.461 | 0.457 | 0.436 | 1.326572008 |
| Q9U329 | CELE_W09C5.8 W09C5.8 | 0.326 | 0.327 | 0.339 | 0.330666667 | 0.452 | 0.443 | 0.423 | 0.439333333 | 1.328629032 |
| P34346 | let-754 C29E4.8 | 0.316 | 0.319 | 0.33 | 0.321666667 | 0.439 | 0.421 | 0.424 | 0.428 | 1.330569948 |
| P52011 | cyn-3 cyp-3 Y75B12B.5 | 0.304 | 0.324 | 0.323 | 0.317 | 0.423 | 0.426 | 0.417 | 0.422 | 1.331230284 |
| P02567 | myo-1 R06C7.10 | 0.328 | 0.331 | 0.34 | 0.333 | 0.452 | 0.435 | 0.443 | 0.443333333 | 1.331331331 |
| Q9U1Z4 | CELE_Y60A3A.9 Y60A3A.9 | 0.411 | 0 | 0.234 | 0.215 | 0.383 | 0 | 0.476 | 0.286333333 | 1.331782946 |
| O45011 | CELE_W10C8.5 W10C8.5 | 0.313 | 0.33 | 0.311 | 0.318 | 0.405 | 0.418 | 0.448 | 0.423666667 | 1.332285115 |
| Q09517 | let-767 dhs-10 C56G2.6 | 0.361 | 0.321 | 0.308 | 0.33 | 0.441 | 0.449 | 0.431 | 0.440333333 | 1.334343434 |
| Q9N4J2 | vit-3 F59D8.1 | 0.32 | 0.284 | 0.304 | 0.302666667 | 0.404 | 0.414 | 0.394 | 0.404 | 1.334801762 |
| Q9GYI1 | CELE_F29B9.11 F29B9.11 | 0.521 | 0.257 | 0.273 | 0.350333333 | 0.469 | 0.486 | 0.449 | 0.468 | 1.335870599 |
| Q21531 | ifg-1 CELE_M110.4 M110.4 | 0.366 | 0.344 | 0.306 | 0.338666667 | 0.43 | 0.514 | 0.416 | 0.453333333 | 1.338582677 |
| Q9GZH4 | ribo-1 T22D1.4 | 0.362 | 0.306 | 0.3 | 0.322666667 | 0.426 | 0.424 | 0.447 | 0.432333333 | 1.339876033 |
| O01816 | cytb-5.2 CELE_W02D3.1 W02D3.1 | 0.304 | 0.283 | 0.306 | 0.297666667 | 0.355 | 0.423 | 0.419 | 0.399 | 1.340425532 |
| P24894 | ctc-2 coII cox-2 MTCE.31 | 0.287 | 0.267 | 0.338 | 0.297333333 | 0.396 | 0.412 | 0.388 | 0.398666667 | 1.340807175 |
| Q18090 | tomm-40 C18E9.6 | 0.299 | 0.312 | 0.34 | 0.317 | 0.439 | 0.413 | 0.426 | 0.426 | 1.34384858 |
| Q2V4S2 | gdi-1 CELE_Y57G11C.10 Y57G11C.10 | 0.321 | 0.438 | 0.318 | 0.359 | 0.463 | 0.529 | 0.458 | 0.483333333 | 1.346332405 |
| Q86NH9 | ttr-41 CELE_F10G7.11 F10G7.11 | 0.317 | 0.306 | 0.288 | 0.303666667 | 0.406 | 0.399 | 0.424 | 0.409666667 | 1.349066959 |
| O61907 | lin-40 CELE_T27C4.4 T27C4.4 | 0.322 | 0 | 0 | 0.107333333 | 0.435 | 0 | 0 | 0.145 | 1.350931677 |
| Q9U307 | gln-3 CELE_Y105C5B.28 Y105C5B.28 | 0.366 | 0.303 | 0.311 | 0.326666667 | 0.447 | 0.434 | 0.443 | 0.441333333 | 1.351020408 |
| O02642 | sucl-2 CELE_F23H11.3 F23H11.3 | 0.33 | 0.306 | 0.296 | 0.310666667 | 0.404 | 0.435 | 0.421 | 0.42 | 1.35193133 |
| P54889 | alh-13 T22H6.2 | 0 | 0 | 0.322 | 0.107333333 | 0 | 0 | 0.436 | 0.145333333 | 1.354037267 |
| Q09450 | C05C10.3 | 0.29 | 0.31 | 0.332 | 0.310666667 | 0.419 | 0.446 | 0.397 | 0.420666667 | 1.354077253 |
| P54811 | cdc-48.1 C06A1.1 | 0 | 0.322 | 0.301 | 0.207666667 | 0 | 0.42 | 0.424 | 0.281333333 | 1.354735152 |
| Q23237 | pbs-3 Y38A8.2 | 0.343 | 0.359 | 0.341 | 0.347666667 | 0.417 | 0.522 | 0.474 | 0.471 | 1.354745925 |
| Q19286 | ifb-2 F10C1.7 | 0.32 | 0.314 | 0.332 | 0.322 | 0.436 | 0.439 | 0.434 | 0.436333333 | 1.355072464 |
| P50093 | phb-2 T24H7.1 | 0.295 | 0.299 | 0.321 | 0.305 | 0.424 | 0.424 | 0.392 | 0.413333333 | 1.355191257 |
| Q21770 | wago-1 R06C7.1 | 0.309 | 0 | 0 | 0.103 | 0.419 | 0 | 0 | 0.139666667 | 1.355987055 |
| Q20655 | ftt-2 F52D10.3 | 0.317 | 0.326 | 0.306 | 0.316333333 | 0.436 | 0.435 | 0.416 | 0.429 | 1.356164384 |
| O61199 | ogdh-1 T22B11.5 | 0.322 | 0.309 | 0.329 | 0.32 | 0.429 | 0.443 | 0.431 | 0.434333333 | 1.357291667 |
| Q20135 | col-81 CELE_F38A3.1 F38A3.1 | 0 | 0.229 | 0.235 | 0.154666667 | 0 | 0.22 | 0.41 | 0.21 | 1.357758621 |
| P48053 | C05D11.1 | 0 | 0.324 | 0 | 0.108 | 0 | 0.44 | 0 | 0.146666667 | 1.358024691 |
| Q95008 | pas-5 F25H2.9 | 0.337 | 0.288 | 0.316 | 0.313666667 | 0.421 | 0.426 | 0.431 | 0.426 | 1.358129649 |
| Q965Q1 | CELE_Y22D7AL.10 Y22D7AL.10 | 0.322 | 0.304 | 0.326 | 0.317333333 | 0.424 | 0.431 | 0.438 | 0.431 | 1.358193277 |

| | | | | | | | | | | |
|---|---|---|---|---|---|---|---|---|---|---|
| Q02335 | ZK370.8 | 0.335 | 0 | 0 | 0.111666667 | 0.455 | 0 | 0 | 0.151666667 | 1.358208955 |
| Q18823 | lam-2 C54D1.5 | 0.331 | 0 | 0.385 | 0.238666667 | 0.482 | 0 | 0.491 | 0.324333333 | 1.358938547 |
| O44549 | acdh-3 CELE_K06A5.6 K06A5.6 | 0.329 | 0.305 | 0.293 | 0.309 | 0.391 | 0.418 | 0.452 | 0.420333333 | 1.36030205 |
| O44400 | F37C4.5 | 0.334 | 0.309 | 0.319 | 0.320666667 | 0.413 | 0.443 | 0.453 | 0.436333333 | 1.360706861 |
| O18239 | sec-61 CELE_Y57G11C.15 Y57G11C.15 | 0.304 | 0.342 | 0.32 | 0.322 | 0.382 | 0.491 | 0.442 | 0.438333333 | 1.361283644 |
| G5EDP2 | daf-22 Y57A10C.6 | 0.301 | 0.465 | 0.288 | 0.351333333 | 0.444 | 0.497 | 0.495 | 0.478666667 | 1.362428843 |
| P12845 | myo-2 T18D3.4 | 0.322 | 0.318 | 0.322 | 0.320666667 | 0.435 | 0.439 | 0.439 | 0.437666667 | 1.364864865 |
| O45713 | CELE_R09B3.3 R09B3.3 | 0.321 | 0.294 | 0.319 | 0.311333333 | 0.505 | 0.394 | 0.376 | 0.425 | 1.36509636 |
| Q19328 | tsn-1 CELE_F10G7.2 F10G7.2 | 0.299 | 0.327 | 0.331 | 0.319 | 0.446 | 0.436 | 0.426 | 0.436 | 1.36677116 |
| O45734 | cpl-1 CELE_T03E6.7 T03E6.7 | 0.318 | 0.318 | 0.327 | 0.321 | 0.441 | 0.414 | 0.462 | 0.439 | 1.367601246 |
| Q23158 | atn-1 CELE_W04D2.1 W04D2.1 | 0.32 | 0.318 | 0.283 | 0.307 | 0.394 | 0.401 | 0.465 | 0.42 | 1.368078176 |
| Q17832 | vgln-1 C08H9.2 CELE_C08H9.2 | 0.301 | 0.322 | 0.332 | 0.318333333 | 0.442 | 0.439 | 0.426 | 0.435666667 | 1.368586387 |
| Q9TYW1 | vha-19 CELE_Y55H10A.1 Y55H10A.1 | 0.312 | 0.322 | 0.302 | 0.312 | 0.444 | 0.403 | 0.434 | 0.427 | 1.368589744 |
| Q95XS2 | CELE_Y38F2AR.9 Y38F2AR.9 | 0.427 | 0 | 0.417 | 0.281333333 | 0.573 | 0 | 0.583 | 0.385333333 | 1.369668246 |
| Q94230 | plp-1 CELE_F45E4.2 F45E4.2 | 0.305 | 0.332 | 0.312 | 0.316333333 | 0.435 | 0.428 | 0.439 | 0.434 | 1.371970495 |
| G5EDD1 | ucr-2.1 CELE_VW06B3R.1 VW06B3R.1 | 0.3 | 0.384 | 0.296 | 0.326666667 | 0.444 | 0.456 | 0.445 | 0.448333333 | 1.37244898 |
| P91427 | pgk-1 T03F1.3 | 0.306 | 0.316 | 0.311 | 0.311 | 0.434 | 0.407 | 0.44 | 0.427 | 1.372990354 |
| G5EGK8 | let-92 F38H4.9 | 0 | 0 | 0.297 | 0.099 | 0 | 0 | 0.408 | 0.136 | 1.373737374 |
| Q21351 | gtbp-1 CELE_K08F4.2 K08F4.2 | 0.309 | 0.301 | 0.296 | 0.302 | 0.42 | 0.402 | 0.423 | 0.415 | 1.374172185 |
| Q19202 | apy-1 F08C6.6 | 0 | 0 | 0.312 | 0.104 | 0 | 0 | 0.429 | 0.143 | 1.375 |
| O17921 | tbb-1 CELE_K01G5.7 K01G5.7 | 0.329 | 0.316 | 0.301 | 0.315333333 | 0.419 | 0.436 | 0.446 | 0.433666667 | 1.375264271 |
| O44738 | CELE_F57B10.5 F57B10.5 | 0 | 0.271 | 0.408 | 0.226333333 | 0 | 0.342 | 0.592 | 0.311333333 | 1.375552283 |
| Q20779 | tag-174 F54D8.2 | 0.295 | 0.318 | 0.32 | 0.311 | 0.452 | 0.419 | 0.415 | 0.428666667 | 1.378349411 |
| Q17574 | tag-165 C01G6.6 | 0.315 | 0.309 | 0.321 | 0.315 | 0.443 | 0.435 | 0.425 | 0.434333333 | 1.378835979 |
| G5ECA7 | CELE_T02D1.8 T02D1.8 | 0 | 0 | 0.321 | 0.107 | 0 | 0 | 0.443 | 0.147666667 | 1.380062305 |
| Q20970 | mrs-1 F58B3.5 | 0.301 | 0.298 | 0.337 | 0.312 | 0.441 | 0.429 | 0.422 | 0.430666667 | 1.38034188 |
| Q19126 | asb-2 CELE_F02E8.1 F02E8.1 | 0.313 | 0.304 | 0.323 | 0.313333333 | 0.426 | 0.435 | 0.437 | 0.432666667 | 1.380851064 |
| B7WNA0 | pyk-1 CELE_F25H5.3 F25H5.3 | 0.346 | 0.33 | 0.308 | 0.328 | 0.431 | 0.462 | 0.466 | 0.453 | 1.381097561 |
| Q20684 | lec-2 CELE_F52H3.7 F52H3.7 | 0.321 | 0.313 | 0.316 | 0.316666667 | 0.442 | 0.459 | 0.413 | 0.438 | 1.383157895 |
| Q23655 | nlt-1 ZK892.2 | 0.301 | 0 | 0.284 | 0.195 | 0.36 | 0 | 0.45 | 0.27 | 1.384615385 |
| P55954 | cco-2 Y37D8A.14 | 0.336 | 0.316 | 0.316 | 0.322666667 | 0.451 | 0.456 | 0.434 | 0.447 | 1.385330579 |
| Q9TYK1 | Y66H1A.4 | 0.309 | 0.344 | 0.338 | 0.330333333 | 0.458 | 0.507 | 0.409 | 0.458 | 1.386478305 |

| | | | | | | | | | | |
|---|---|---|---|---|---|---|---|---|---|---|
| O62102 | pbs-2 C47B2.4 CELE_C47B2.4 | 0.357 | 0.418 | 0 | 0.258333333 | 0.47 | 0.605 | 0 | 0.358333333 | 1.387096774 |
| G5EGK1 | tln-1 CELE_Y71G12B.11 Y71G12B.11 | 0.284 | 0 | 0 | 0.094666667 | 0.394 | 0 | 0 | 0.131333333 | 1.387323944 |
| Q9UAQ6 | rab-1 C39F7.4 CELE_C39F7.4 | 0.326 | 0.303 | 0.299 | 0.309333333 | 0.407 | 0.434 | 0.447 | 0.429333333 | 1.387931034 |
| Q9BKU4 | phb-1 Y37E3.9 | 0.325 | 0.288 | 0.301 | 0.304666667 | 0.421 | 0.427 | 0.421 | 0.423 | 1.388402626 |
| Q9U2D6 | cnp-2 Y46G5A.10 | 0 | 0 | 0.299 | 0.099666667 | 0 | 0 | 0.416 | 0.138666667 | 1.391304348 |
| Q22494 | vha-15 T14F9.1 | 0.32 | 0.342 | 0.301 | 0.321 | 0.455 | 0.447 | 0.438 | 0.446666667 | 1.391484943 |
| Q9NEN6 | rps-6 Y71A12B.1 | 0.311 | 0.319 | 0.335 | 0.321666667 | 0.456 | 0.445 | 0.442 | 0.447666667 | 1.391709845 |
| Q95Q60 | fkb-5 C50F2.6 CELE_C50F2.6 | 0.471 | 0.246 | 0.356 | 0.357666667 | 0.458 | 0.494 | 0.542 | 0.498 | 1.392357875 |
| Q86S57 | alh-4 CELE_T05H4.13 T05H4.13 | 0.292 | 0 | 0.323 | 0.205 | 0.467 | 0 | 0.39 | 0.285666667 | 1.393495935 |
| Q9XX57 | dct-16 CELE_Y38H6C.1 Y38H6C.1 | 0.309 | 0.311 | 0.31 | 0.31 | 0.43 | 0.428 | 0.438 | 0.432 | 1.393548387 |
| Q8WQA8 | rps-20 CELE_Y105E8A.16 Y105E8A.16 | 0.31 | 0.314 | 0.316 | 0.313333333 | 0.438 | 0.435 | 0.437 | 0.436666667 | 1.393617021 |
| O16249 | CELE_F44E7.4 F44E7.4 | 0.318 | 0.309 | 0.354 | 0.327 | 0.446 | 0.449 | 0.473 | 0.456 | 1.394495413 |
| Q18212 | hel-1 C26D10.2 | 0.292 | 0.316 | 0.35 | 0.319333333 | 0.462 | 0.445 | 0.429 | 0.445333333 | 1.394572025 |
| Q94246 | gfi-1 CELE_F57F4.3 F57F4.3 | 0.3 | 0.297 | 0.329 | 0.308666667 | 0.437 | 0.431 | 0.424 | 0.430666667 | 1.39524838 |
| G5EBJ7 | fbp-1 fbp CELE_K07A3.1 K07A3.1 | 0.342 | 0.303 | 0.296 | 0.313666667 | 0.423 | 0.453 | 0.438 | 0.438 | 1.396386823 |
| Q9N3B0 | Y54G2A.23 | 0.287 | 0.318 | 0.311 | 0.305333333 | 0.401 | 0.432 | 0.447 | 0.426666667 | 1.397379913 |
| Q20676 | pccb-1 CELE_F52E4.1 F52E4.1 | 0.326 | 0.317 | 0.306 | 0.316333333 | 0.446 | 0.437 | 0.444 | 0.442333333 | 1.398314015 |
| Q9XTI0 | B0250.5 | 0.292 | 0.294 | 0.297 | 0.294333333 | 0.409 | 0.405 | 0.421 | 0.411666667 | 1.398640997 |
| Q95X44 | vha-8 C17H12.14 CELE_C17H12.14 | 0.31 | 0.315 | 0.32 | 0.315 | 0.437 | 0.441 | 0.444 | 0.440666667 | 1.398941799 |
| Q18577 | C42D4.1 CELE_C42D4.1 | 0.331 | 0 | 0.308 | 0.213 | 0.433 | 0 | 0.461 | 0.298 | 1.399061033 |
| O76367 | CELE_F29C4.2 F29C4.2 | 0.285 | 0.256 | 0 | 0.180333333 | 0.38 | 0.377 | 0 | 0.252333333 | 1.399260628 |
| G5EE96 | slc-25a10 slc-25A10 CELE_K11G12.5 K11G12.5 | 0 | 0 | 0.318 | 0.106 | 0 | 0 | 0.445 | 0.148333333 | 1.399371069 |
| Q4W5P0 | sqd-1 CELE_Y73B6BL.6 Y73B6BL.6 | 0.333 | 0.313 | 0.332 | 0.326 | 0.418 | 0.535 | 0.416 | 0.456333333 | 1.399795501 |
| Q21746 | sgt-1 CELE_R05F9.10 R05F9.10 | 0.317 | 0.313 | 0.324 | 0.318 | 0.431 | 0.455 | 0.452 | 0.446 | 1.402515723 |
| Q22633 | hpd-1 T21C12.2 | 0.296 | 0.365 | 0.307 | 0.322666667 | 0.47 | 0.431 | 0.457 | 0.452666667 | 1.402892562 |
| Q9N5U1 | CELE_T22F3.3 T22F3.3 | 0.305 | 0.328 | 0.302 | 0.311666667 | 0.443 | 0.446 | 0.423 | 0.437333333 | 1.403208556 |
| Q8MNV8 | C14B9.10 CELE_C14B9.10 | 0 | 0.273 | 0.322 | 0.198333333 | 0 | 0.436 | 0.399 | 0.278333333 | 1.403361345 |
| Q7Z1Q3 | alh-12 CELE_Y69F12A.2 Y69F12A.2 | 0.239 | 0.321 | 0 | 0.186666667 | 0.369 | 0.417 | 0 | 0.262 | 1.403571429 |
| H2KZF5 | pmt-1 CELE_ZK622.3 ZK622.3 | 0.331 | 0.346 | 0.301 | 0.326 | 0.444 | 0.463 | 0.466 | 0.457666667 | 1.403885481 |
| Q10129 | mrps-16 F56D1.3 | 0 | 0.297 | 0 | 0.099 | 0 | 0.417 | 0 | 0.139 | 1.404040404 |
| Q95XI9 | CELE_Y69A2AR.18 Y69A2AR.18 | 0.315 | 0.312 | 0.306 | 0.311 | 0.428 | 0.439 | 0.443 | 0.436666667 | 1.404072883 |
| O01802 | rpl-7 F53G12.10 | 0.306 | 0.315 | 0.335 | 0.318666667 | 0.453 | 0.446 | 0.444 | 0.447666667 | 1.404811715 |

| | | | | | | | | | | |
|---|---|---|---|---|---|---|---|---|---|---|
| Q20224 | lbp-2 F40F4.2 | 0.322 | 0.362 | 0.279 | 0.321 | 0.431 | 0.453 | 0.469 | 0.451 | 1.404984424 |
| B3WFT8 | C14F11.4 CELE_C14F11.4 | 0 | 0 | 0.301 | 0.100333333 | 0 | 0 | 0.423 | 0.141 | 1.405315615 |
| P11141 | hsp-6 hsp70f C37H5.8 | 0.302 | 0.317 | 0.3 | 0.306333333 | 0.433 | 0.435 | 0.424 | 0.430666667 | 1.405875952 |
| O62213 | cey-1 CELE_F33A8.3 F33A8.3 | 0.325 | 0.327 | 0.326 | 0.326 | 0.452 | 0.458 | 0.467 | 0.459 | 1.40797546 |
| Q9XTT9 | rpt-6 CELE_Y49E10.1 Y49E10.1 | 0.415 | 0.348 | 0.298 | 0.353666667 | 0.487 | 0.485 | 0.522 | 0.498 | 1.408105561 |
| G5EEK8 | sca-1 mca-4 CELE_K11D9.2 K11D9.2 | 0.306 | 0.314 | 0.31 | 0.31 | 0.439 | 0.44 | 0.432 | 0.437 | 1.409677419 |
| Q8MXS8 | CELE_Y47G6A.22 Y47G6A.22 | 0 | 0.316 | 0.329 | 0.215 | 0 | 0.47 | 0.44 | 0.303333333 | 1.410852713 |
| O18650 | rps-19 T05F1.3 | 0.295 | 0.333 | 0.309 | 0.312333333 | 0.428 | 0.459 | 0.435 | 0.440666667 | 1.410885806 |
| Q9XXK1 | H28O16.1 | 0.302 | 0.318 | 0.312 | 0.310666667 | 0.439 | 0.431 | 0.445 | 0.438333333 | 1.410944206 |
| P34328 | hsp-12.2 hsp12-2 C14B9.1 | 0.328 | 0.285 | 0.314 | 0.309 | 0.445 | 0.43 | 0.433 | 0.436 | 1.411003236 |
| G5EBP5 | CELE_ZC247.1 ZC247.1 | 0.298 | 0.308 | 0.315 | 0.307 | 0.439 | 0.43 | 0.431 | 0.433333333 | 1.411509229 |
| O01692 | rps-17 T08B2.10 | 0.308 | 0.318 | 0.31 | 0.312 | 0.438 | 0.439 | 0.446 | 0.441 | 1.413461538 |
| Q7K707 | gpi-1 CELE_Y87G2A.8 Y87G2A.8 | 0.294 | 0.419 | 0.342 | 0.351666667 | 0.524 | 0.51 | 0.458 | 0.497333333 | 1.414218009 |
| P54412 | eef-1G F17C11.9 | 0.324 | 0.32 | 0.311 | 0.318333333 | 0.454 | 0.452 | 0.445 | 0.450333333 | 1.414659686 |
| G5EFG4 | abcf-2 CELE_T27E9.7 T27E9.7 | 0.304 | 0.319 | 0.324 | 0.315666667 | 0.423 | 0.475 | 0.442 | 0.446666667 | 1.41499472 |
| P90762 | rmd-2 C27H6.4 CELE_C27H6.4 | 0.31 | 0.313 | 0.295 | 0.306 | 0.427 | 0.433 | 0.439 | 0.433 | 1.41503268 |
| P10299 | gst-1 R107.7 | 0.328 | 0.311 | 0.371 | 0.336666667 | 0.483 | 0.511 | 0.438 | 0.477333333 | 1.417821782 |
| H2KZ19 | ampd-1 C34F11.3 CELE_C34F11.3 | 0.322 | 0 | 0 | 0.107333333 | 0.457 | 0 | 0 | 0.152333333 | 1.419254658 |
| P90735 | eat-6 B0365.3 CELE_B0365.3 | 0.305 | 0.321 | 0.304 | 0.31 | 0.439 | 0.442 | 0.439 | 0.44 | 1.419354839 |
| O16517 | atp-4 CELE_T05H4.12 T05H4.12 | 0.298 | 0.3 | 0.308 | 0.302 | 0.427 | 0.429 | 0.43 | 0.428666667 | 1.419426049 |
| Q17763 | atp-5 C06H2.1 CELE_C06H2.1 | 0.31 | 0.3 | 0.299 | 0.303 | 0.424 | 0.44 | 0.427 | 0.430333333 | 1.420242024 |
| Q10576 | dpy-18 phy-1 Y47D3B.10 | 0.414 | 0.277 | 0.292 | 0.327666667 | 0.384 | 0.528 | 0.485 | 0.465666667 | 1.421159715 |
| Q09975 | lys-8 C17G10.5 CELE_C17G10.5 | 0.299 | 0 | 0 | 0.099666667 | 0.425 | 0 | 0 | 0.141666667 | 1.421404682 |
| P09588 | his-3 T10C6.12; his-7 F45F2.4; his-12 ZK131.6; his-16 ZK131.10; his-19 K06C4.11; his-21 K06C4.3; his-30 F35H10.1; his-33 F17E9.13; his-43 F08G2.2; his-47 B0035.7; his-51 F07B7.10; his-53 F07B7.3; his-57 F54E12.5; his-61 F55G1.10; his-65 H02I12.7; his-68 T23D8.6 | 0.331 | 0.322 | 0.353 | 0.335333333 | 0.558 | 0.438 | 0.434 | 0.476666667 | 1.421471173 |
| P28548 | kin-10 kin-5 T01G9.6 | 0.327 | 0 | 0 | 0.109 | 0.465 | 0 | 0 | 0.155 | 1.422018349 |
| Q18678 | srs-2 C47E12.1 | 0.3 | 0.298 | 0.317 | 0.305 | 0.435 | 0.416 | 0.451 | 0.434 | 1.42295082 |
| Q9NES7 | CELE_Y39B6A.1 Y39B6A.1 | 0.235 | 0.274 | 0.304 | 0.271 | 0.359 | 0.406 | 0.392 | 0.385666667 | 1.423124231 |
| Q21481 | dhs-28 CELE_M03A8.1 M03A8.1 | 0.317 | 0.3 | 0.314 | 0.310333333 | 0.442 | 0.446 | 0.437 | 0.441666667 | 1.423200859 |

| | | | | | | | | | | |
|---|---|---|---|---|---|---|---|---|---|---|
| O45552 | acaa-2 CELE_F53A2.7 F53A2.7 | 0.323 | 0.29 | 0.315 | 0.309333333 | 0.439 | 0.449 | 0.433 | 0.440333333 | 1.423491379 |
| P53014 | mlc-3 F09F7.2 | 0.314 | 0.308 | 0.31 | 0.310666667 | 0.439 | 0.444 | 0.444 | 0.442333333 | 1.423819742 |
| P50880 | rpl-3 F13B10.2 | 0.308 | 0.301 | 0.316 | 0.308333333 | 0.448 | 0.439 | 0.432 | 0.439666667 | 1.425945946 |
| P34466 | clu-1 F55H2.6 | 0.481 | 0.3 | 0.265 | 0.348666667 | 0.519 | 0.467 | 0.506 | 0.497333333 | 1.426386233 |
| Q20585 | rpn-7 F49C12.8 | 0.342 | 0.277 | 0 | 0.206333333 | 0.409 | 0.474 | 0 | 0.294333333 | 1.426494346 |
| O16305 | cmd-1 T21H3.3 | 0.296 | 0.326 | 0.311 | 0.311 | 0.436 | 0.456 | 0.439 | 0.443666667 | 1.426580922 |
| P34460 | eef-1B.1 F54H12.6 | 0.342 | 0.332 | 0.266 | 0.313333333 | 0.431 | 0.44 | 0.47 | 0.447 | 1.426595745 |
| P49196 | rps-12 F54E7.2 | 0.284 | 0.314 | 0.3 | 0.299333333 | 0.432 | 0.422 | 0.428 | 0.427333333 | 1.427616927 |
| G5EDV3 | cey-4 CELE_Y39A1C.3 Y39A1C.3 | 0.305 | 0.3 | 0.331 | 0.312 | 0.458 | 0.438 | 0.441 | 0.445666667 | 1.428418803 |
| P06125 | vit-5 C04F6.1 | 0.3 | 0.292 | 0.275 | 0.289 | 0.411 | 0.412 | 0.417 | 0.413333333 | 1.430219146 |
| Q21525 | CELE_M05D6.6 M05D6.6 | 0.303 | 0 | 0.299 | 0.200666667 | 0.451 | 0 | 0.41 | 0.287 | 1.430232558 |
| Q23451 | rad-23 CELE_ZK20.3 ZK20.3 | 0 | 0.33 | 0 | 0.11 | 0 | 0.472 | 0 | 0.157333333 | 1.43030303 |
| Q09533 | rpl-10 F10B5.1 | 0.305 | 0.33 | 0.313 | 0.316 | 0.453 | 0.455 | 0.448 | 0.452 | 1.430379747 |
| Q21408 | spc-1 CELE_K10B3.10 K10B3.10 | 0.361 | 0.274 | 0.324 | 0.319666667 | 0.475 | 0.44 | 0.457 | 0.457333333 | 1.430656934 |
| P27639 | inf-1 F57B9.6 | 0.34 | 0.286 | 0.302 | 0.309333333 | 0.429 | 0.452 | 0.447 | 0.442666667 | 1.431034483 |
| Q21443 | lmn-1 lam-1 DY3.2 | 0.32 | 0.333 | 0.326 | 0.326333333 | 0.47 | 0.457 | 0.474 | 0.467 | 1.431052094 |
| Q9BL15 | CELE_Y48G8AL.5 Y48G8AL.5 | 0.37 | 0.254 | 0 | 0.208 | 0.414 | 0.479 | 0 | 0.297666667 | 1.431089744 |
| Q21750 | aagr-2 CELE_R05F9.12 R05F9.12 | 0.283 | 0 | 0 | 0.094333333 | 0.405 | 0 | 0 | 0.135 | 1.431095406 |
| P50432 | mel-32 gly-1 C05D11.11 | 0.307 | 0.304 | 0.321 | 0.310666667 | 0.435 | 0.452 | 0.448 | 0.445 | 1.432403433 |
| P17139 | emb-9 clb-2 K04H4.1 | 0.308 | 0.278 | 0.316 | 0.300666667 | 0.436 | 0.435 | 0.422 | 0.431 | 1.433481153 |
| Q22641 | lpd-9 CELE_T21C9.5 T21C9.5 | 0.258 | 0.353 | 0 | 0.203666667 | 0.479 | 0.397 | 0 | 0.292 | 1.433715221 |
| Q03577 | drs-1 B0464.1 | 0.318 | 0.279 | 0.326 | 0.307666667 | 0.442 | 0.459 | 0.423 | 0.441333333 | 1.434452871 |
| Q22716 | rpl-32 CELE_T24B8.1 T24B8.1 | 0.304 | 0.306 | 0.305 | 0.305 | 0.444 | 0.431 | 0.438 | 0.437666667 | 1.434972678 |
| Q9XWP7 | eif-3.j eif-3.J CELE_Y40B1B.5 Y40B1B.5 | 0.332 | 0.253 | 0.35 | 0.311666667 | 0.473 | 0.423 | 0.446 | 0.447333333 | 1.435294118 |
| G5ECM6 | tufm-1 CELE_Y71H2AM.23 Y71H2AM.23 | 0.319 | 0.317 | 0.306 | 0.314 | 0.467 | 0.43 | 0.456 | 0.451 | 1.436305732 |
| A7LPG5 | cisd-1 CELE_W02B12.15 W02B12.15 | 0.275 | 0.337 | 0.291 | 0.301 | 0.44 | 0.425 | 0.432 | 0.432333333 | 1.436323367 |
| Q19706 | eif-3.G F22B5.2 | 0.364 | 0 | 0.527 | 0.297 | 0.388 | 0.599 | 0.293 | 0.426666667 | 1.436588103 |
| Q19749 | dlat-1 F23B12.5 | 0.291 | 0.306 | 0.31 | 0.302333333 | 0.421 | 0.432 | 0.45 | 0.434333333 | 1.43660419 |
| O45815 | act-5 CELE_T25C8.2 T25C8.2 | 0.304 | 0.3 | 0.319 | 0.307666667 | 0.44 | 0.45 | 0.437 | 0.442333333 | 1.437703142 |
| Q9U2H9 | eef-1B.2 Y41E3.10 | 0.312 | 0.307 | 0.312 | 0.310333333 | 0.432 | 0.445 | 0.462 | 0.446333333 | 1.438238453 |
| Q95YF3 | cgh-1 C07H6.5 | 0.3 | 0.293 | 0.316 | 0.303 | 0.422 | 0.436 | 0.45 | 0.436 | 1.438943894 |
| Q09622 | puf-12 ZK945.3 | 0 | 0.41 | 0 | 0.136666667 | 0 | 0.59 | 0 | 0.196666667 | 1.43902439 |
| O01829 | ssp-19 C55C2.2 | 0.32 | 0.304 | 0.23 | 0.284666667 | 0.369 | 0.438 | 0.422 | 0.409666667 | 1.43911007 |

| | | | | | | | | | | |
|---|---|---|---|---|---|---|---|---|---|---|
| Q9NA78 | CELE_Y57A10A.23 Y57A10A.23 | 0.349 | 0.317 | 0.299 | 0.321666667 | 0.472 | 0.485 | 0.433 | 0.463333333 | 1.440414508 |
| G5EEH6 | ivd-1 C02B10.1 CELE_C02B10.1 | 0.313 | 0.263 | 0.289 | 0.288333333 | 0.421 | 0.398 | 0.427 | 0.415333333 | 1.440462428 |
| P53013 | eft-3 F31E3.5; eft-4 R03G5.1 | 0.305 | 0.308 | 0.304 | 0.305666667 | 0.442 | 0.441 | 0.438 | 0.440333333 | 1.440567067 |
| Q9N4J8 | cct-3 F54A3.3 | 0.315 | 0.312 | 0.312 | 0.313 | 0.464 | 0.444 | 0.445 | 0.451 | 1.440894569 |
| O61820 | eif-3.E B0511.10 | 0 | 0.336 | 0.319 | 0.218333333 | 0 | 0.463 | 0.481 | 0.314666667 | 1.441221374 |
| Q9U348 | col-93 CELE_W05B2.5 W05B2.5 | 0.304 | 0.311 | 0.338 | 0.317666667 | 0.459 | 0.452 | 0.463 | 0.458 | 1.441762854 |
| Q93379 | gsr-1 C46F11.2 | 0.362 | 0.294 | 0.29 | 0.315333333 | 0.485 | 0.45 | 0.429 | 0.454666667 | 1.441860465 |
| Q21832 | rnp-4 RBM8 Y14 CELE_R07E5.14 R07E5.14 | 0 | 0.267 | 0 | 0.089 | 0 | 0.385 | 0 | 0.128333333 | 1.441947566 |
| Q22288 | ttr-15 T07C4.5 | 0.301 | 0.284 | 0.311 | 0.298666667 | 0.44 | 0.422 | 0.43 | 0.430666667 | 1.441964286 |
| Q27527 | enol-1 T21B10.2 | 0.305 | 0.312 | 0.305 | 0.307333333 | 0.445 | 0.447 | 0.44 | 0.444 | 1.444685466 |
| O45060 | C35B1.5 CELE_C35B1.5 | 0.288 | 0.333 | 0.366 | 0.329 | 0.469 | 0.497 | 0.46 | 0.475333333 | 1.444782168 |
| P36573 | lec-1 W09H1.6 | 0.305 | 0.303 | 0.292 | 0.3 | 0.434 | 0.432 | 0.435 | 0.433666667 | 1.445555556 |
| O44898 | CELE_ZK484.5 ZK484.5 | 0.296 | 0.282 | 0.29 | 0.289333333 | 0.416 | 0.414 | 0.425 | 0.418333333 | 1.445852535 |
| O61848 | CELE_K03E5.2 K03E5.2 | 0 | 0.259 | 0.303 | 0.187333333 | 0 | 0.402 | 0.411 | 0.271 | 1.446619217 |
| Q17361 | usp-14 tgt-1 C13B4.2 | 0 | 0.36 | 0.264 | 0.208 | 0 | 0.428 | 0.475 | 0.301 | 1.447115385 |
| P48150 | rps-14 F37C12.9 | 0.299 | 0.325 | 0.317 | 0.313666667 | 0.458 | 0.455 | 0.449 | 0.454 | 1.447396387 |
| Q27888 | ldh-1 F13D12.2 | 0.272 | 0.329 | 0.317 | 0.306 | 0.412 | 0.372 | 0.545 | 0.443 | 1.447712418 |
| Q23621 | gdh-1 CELE_ZK829.4 ZK829.4 | 0.303 | 0.289 | 0.301 | 0.297666667 | 0.429 | 0.437 | 0.427 | 0.431 | 1.447928331 |
| H9G2T4 | idh-1 CELE_F59B8.2 F59B8.2 | 0.313 | 0.312 | 0.307 | 0.310666667 | 0.457 | 0.453 | 0.441 | 0.450333333 | 1.449570815 |
| Q10454 | F46H5.3 | 0.315 | 0.286 | 0.311 | 0.304 | 0.442 | 0.44 | 0.442 | 0.441333333 | 1.451754386 |
| Q9N5B3 | CELE_W08E12.7 W08E12.7 | 0.3 | 0.297 | 0.305 | 0.300666667 | 0.44 | 0.436 | 0.435 | 0.437 | 1.453436807 |
| P34519 | K11H3.3 | 0.28 | 0.302 | 0.347 | 0.309666667 | 0.472 | 0.439 | 0.44 | 0.450333333 | 1.454251884 |
| Q7JNG1 | atp-3 CELE_F27C1.7 F27C1.7 | 0.299 | 0.286 | 0.31 | 0.298333333 | 0.441 | 0.433 | 0.428 | 0.434 | 1.454748603 |
| P34455 | aco-2 F54H12.1 | 0.303 | 0.29 | 0.309 | 0.300666667 | 0.438 | 0.432 | 0.443 | 0.437666667 | 1.455654102 |
| P49180 | rpl-33 F10E7.7 | 0.316 | 0.325 | 0.302 | 0.314333333 | 0.455 | 0.459 | 0.459 | 0.457666667 | 1.455991516 |
| P98080 | ucr-1 F56D2.1 | 0.296 | 0.3 | 0.309 | 0.301666667 | 0.445 | 0.443 | 0.43 | 0.439333333 | 1.456353591 |
| O61708 | pqn-59 CELE_R119.4 R119.4 | 0.291 | 0.289 | 0.3 | 0.293333333 | 0.397 | 0.434 | 0.451 | 0.427333333 | 1.456818182 |
| Q21824 | prdx-3 R07E5.2 | 0.325 | 0.283 | 0.315 | 0.307666667 | 0.443 | 0.45 | 0.453 | 0.448666667 | 1.458288191 |
| Q9N5U5 | cdc-73 F35F11.1 | 0.303 | 0 | 0 | 0.101 | 0.442 | 0 | 0 | 0.147333333 | 1.458745875 |
| P02566 | unc-54 myo-4 F11C3.3 | 0.306 | 0.304 | 0.305 | 0.305 | 0.443 | 0.449 | 0.443 | 0.445 | 1.459016393 |
| Q18853 | cyc-1 C54G4.8 CELE_C54G4.8 | 0.303 | 0.291 | 0.316 | 0.303333333 | 0.443 | 0.44 | 0.445 | 0.442666667 | 1.459340659 |
| Q9XXR4 | ttr-24 CELE_Y51A2D.9 Y51A2D.9 | 0.241 | 0 | 0.252 | 0.164333333 | 0.387 | 0 | 0.334 | 0.240333333 | 1.462474645 |
| G5EGP4 | vha-6 CELE_VW02B12L.1 VW02B12L.1 | 0 | 0 | 0.406 | 0.135333333 | 0 | 0 | 0.594 | 0.198 | 1.463054187 |

| | | | | | | | | | | |
|---|---|---|---|---|---|---|---|---|---|---|
| Q18885 | icd-1 C56C10.8 | 0.308 | 0.292 | 0.311 | 0.303666667 | 0.438 | 0.455 | 0.44 | 0.444333333 | 1.463227223 |
| O18240 | rps-18 CELE_Y57G11C.16 Y57G11C.16 | 0.313 | 0.295 | 0.309 | 0.305666667 | 0.441 | 0.442 | 0.459 | 0.447333333 | 1.46346783 |
| Q18680 | pyp-1 uba-1 C47E12.4 | 0.289 | 0.31 | 0.285 | 0.294666667 | 0.427 | 0.429 | 0.438 | 0.431333333 | 1.463800905 |
| P37165 | ubl-1 H06I04.4 | 0.308 | 0.313 | 0.301 | 0.307333333 | 0.448 | 0.453 | 0.449 | 0.45 | 1.464208243 |
| Q20772 | F54D5.7 | 0 | 0.285 | 0.288 | 0.191 | 0 | 0.406 | 0.433 | 0.279666667 | 1.464223386 |
| P49181 | rpl-36 F37C12.4 | 0.303 | 0.322 | 0.314 | 0.313 | 0.46 | 0.45 | 0.465 | 0.458333333 | 1.464323749 |
| O17861 | F37H8.5 | 0.277 | 0.281 | 0.296 | 0.284666667 | 0.436 | 0.4 | 0.415 | 0.417 | 1.464871194 |
| P49029 | mag-1 R09B3.5 | 0 | 0.307 | 0.273 | 0.193333333 | 0 | 0.414 | 0.436 | 0.283333333 | 1.465517241 |
| Q9N4M4 | anc-1 ZK973.6 | 0.309 | 0.309 | 0.307 | 0.308333333 | 0.455 | 0.453 | 0.448 | 0.452 | 1.465945946 |
| Q22993 | pmt-2 CELE_F54D11.1 F54D11.1 | 0.295 | 0.321 | 0.309 | 0.308333333 | 0.462 | 0.45 | 0.445 | 0.452333333 | 1.467027027 |
| Q86D21 | tars-1 C47D12.6 CELE_C47D12.6 | 0.314 | 0.294 | 0.282 | 0.296666667 | 0.422 | 0.427 | 0.458 | 0.435666667 | 1.468539326 |
| Q17624 | arrd-25 C04C11.2 CELE_C04C11.2 | 0.344 | 0 | 0.296 | 0.213333333 | 0.484 | 0 | 0.456 | 0.313333333 | 1.46875 |
| P0CG71 | ubq-1 ubia F25B5.4 | 0.286 | 0.296 | 0.284 | 0.288666667 | 0.421 | 0.425 | 0.427 | 0.424333333 | 1.469976905 |
| P91423 | CELE_T03F1.11 T03F1.11 | 1 | 0 | 0.253 | 0.417666667 | 0.688 | 0.692 | 0.462 | 0.614 | 1.470071828 |
| P52013 | cyn-5 cyp-5 F31C3.1 | 0.305 | 0.287 | 0.296 | 0.296 | 0.445 | 0.429 | 0.432 | 0.435333333 | 1.470720721 |
| Q22347 | acdh-10 T08G2.3 | 0.412 | 0.294 | 0.277 | 0.327666667 | 0.56 | 0.444 | 0.442 | 0.482 | 1.471007121 |
| P91134 | adss-1 C37H5.6 | 0.337 | 0.323 | 0.314 | 0.324666667 | 0.452 | 0.548 | 0.433 | 0.477666667 | 1.471252567 |
| Q19420 | ttx-7 F13G3.5 | 0.35 | 0.262 | 0.327 | 0.313 | 0.421 | 0.467 | 0.494 | 0.460666667 | 1.471778488 |
| Q19339 | acs-14 CELE_F11A3.1 F11A3.1 | 0 | 0.279 | 0.31 | 0.196333333 | 0 | 0.418 | 0.449 | 0.289 | 1.471986418 |
| Q20647 | rpl-25.2 F52B5.6 | 0.29 | 0.309 | 0.309 | 0.302666667 | 0.445 | 0.454 | 0.438 | 0.445666667 | 1.47246696 |
| P51404 | rps-13 C16A3.9 | 0.307 | 0.306 | 0.314 | 0.309 | 0.463 | 0.454 | 0.448 | 0.455 | 1.472491909 |
| H2L0I2 | lfi-1 CELE_ZC8.4 ZC8.4 | 0 | 0.328 | 0 | 0.109333333 | 0 | 0.483 | 0 | 0.161 | 1.472560976 |
| Q9BKU5 | CELE_Y37E3.8 Y37E3.8 | 0.304 | 0.296 | 0.298 | 0.299333333 | 0.44 | 0.442 | 0.442 | 0.441333333 | 1.474387528 |
| P91128 | rpl-13 C32E8.2 | 0.317 | 0.294 | 0.308 | 0.306333333 | 0.451 | 0.455 | 0.449 | 0.451666667 | 1.474428727 |
| P52015 | cyn-7 cyp-7 Y75B12B.2 | 0.303 | 0.31 | 0.308 | 0.307 | 0.448 | 0.452 | 0.458 | 0.452666667 | 1.474484256 |
| Q19087 | dnpp-1 F01F1.9 | 0.294 | 0.321 | 0.352 | 0.322333333 | 0.47 | 0.504 | 0.452 | 0.475333333 | 1.474663909 |
| P47207 | cct-2 cctb T21B10.7 | 0.321 | 0.298 | 0.316 | 0.311666667 | 0.452 | 0.475 | 0.452 | 0.459666667 | 1.47486631 |
| Q9TYY0 | CELE_M57.2 M57.2 | 0.309 | 0 | 0 | 0.103 | 0.456 | 0 | 0 | 0.152 | 1.475728155 |
| Q18943 | CELE_D1054.10 D1054.10 | 0.389 | 0.278 | 0.306 | 0.324333333 | 0.489 | 0.513 | 0.434 | 0.478666667 | 1.475847893 |
| Q966C6 | rpl-7A Y24D9A.4 | 0.308 | 0.306 | 0.312 | 0.308666667 | 0.458 | 0.45 | 0.459 | 0.455666667 | 1.476241901 |
| G5EGP8 | cpz-1 CELE_F32B5.8 F32B5.8 | 0.317 | 0.292 | 0.318 | 0.309 | 0.441 | 0.459 | 0.469 | 0.456333333 | 1.476806904 |
| D0IMZ5 | fln-1 CELE_Y66H1B.2 Y66H1B.2 | 0.33 | 0.295 | 0.293 | 0.306 | 0.443 | 0.448 | 0.465 | 0.452 | 1.477124183 |
| G5EGL2 | CELE_T28D6.6 T28D6.6 | 0 | 0 | 0.289 | 0.096333333 | 0 | 0 | 0.427 | 0.142333333 | 1.477508651 |
| O01541 | aars-2 ars-2 F28H1.3 | 0.313 | 0.31 | 0.312 | 0.311666667 | 0.46 | 0.474 | 0.448 | 0.460666667 | 1.478074866 |

| | | | | | | | | | | |
|---|---|---|---|---|---|---|---|---|---|---|
| Q17633 | C04G2.9 CELE_C04G2.9 | 0.291 | 0 | 0.263 | 0.184666667 | 0.404 | 0 | 0.415 | 0.273 | 1.47833935 |
| Q07750 | unc-60 C38C3.5 | 0.269 | 0.339 | 0.337 | 0.315 | 0.479 | 0.449 | 0.47 | 0.466 | 1.479365079 |
| O62431 | ers-1 qrs-5 Y41E3.4 | 0.32 | 0.256 | 0.333 | 0.303 | 0.431 | 0.475 | 0.439 | 0.448333333 | 1.479647965 |
| Q19278 | hach-1 CELE_F09F7.4 F09F7.4 | 0.314 | 0.321 | 0.309 | 0.314666667 | 0.461 | 0.471 | 0.465 | 0.465666667 | 1.479872881 |
| P91871 | fasn-1 CELE_F32H2.5 F32H2.5 | 0.322 | 0.356 | 0.295 | 0.324333333 | 0.482 | 0.47 | 0.488 | 0.48 | 1.47995889 |
| O02108 | cdc-37 W08F4.8 | 0.302 | 0 | 0 | 0.100666667 | 0.447 | 0 | 0 | 0.149 | 1.48013245 |
| P34689 | glh-1 T21G5.3 | 0.31 | 0.319 | 0.304 | 0.311 | 0.445 | 0.464 | 0.473 | 0.460666667 | 1.481243301 |
| P46563 | aldo-2 F01F1.12 | 0.314 | 0.299 | 0.298 | 0.303666667 | 0.445 | 0.453 | 0.452 | 0.45 | 1.481888035 |
| P91253 | gst-7 F11G11.2 | 0.291 | 0.31 | 0.347 | 0.316 | 0.53 | 0.444 | 0.432 | 0.468666667 | 1.483122363 |
| O17406 | attf-2 CELE_F09G2.9 F09G2.9 | 0.301 | 0.343 | 0.279 | 0.307666667 | 0.452 | 0.414 | 0.503 | 0.456333333 | 1.483206934 |
| P61866 | rpl-12 JC8.3 | 0.303 | 0.294 | 0.299 | 0.298666667 | 0.442 | 0.446 | 0.441 | 0.443 | 1.483258929 |
| Q27389 | rpl-16 M01F1.2 | 0.315 | 0.288 | 0.305 | 0.302666667 | 0.444 | 0.452 | 0.451 | 0.449 | 1.483480176 |
| G5EBY3 | nmy-2 CELE_F20G4.3 F20G4.3 | 0 | 0 | 0.254 | 0.084666667 | 0 | 0 | 0.377 | 0.125666667 | 1.484251969 |
| Q9U3Q0 | mrps-22 C14A4.14 CELE_C14A4.14 | 0.309 | 0 | 0 | 0.103 | 0.459 | 0 | 0 | 0.153 | 1.485436893 |
| Q09544 | F58F12.1 | 0.295 | 0.294 | 0.298 | 0.295666667 | 0.442 | 0.437 | 0.439 | 0.439333333 | 1.485907554 |
| P52899 | T05H10.6 | 0.279 | 0.31 | 0.312 | 0.300333333 | 0.447 | 0.437 | 0.455 | 0.446333333 | 1.486126526 |
| G5EGA5 | fat-2 W02A2.1 | 0.263 | 0.297 | 0.312 | 0.290666667 | 0.467 | 0.398 | 0.431 | 0.432 | 1.486238532 |
| Q20938 | rpn-6.1 F57B9.10 | 0.321 | 0.301 | 0.224 | 0.282 | 0.445 | 0.459 | 0.354 | 0.419333333 | 1.486997636 |
| Q9U2M4 | Y38F1A.6 | 0.279 | 0.295 | 0.333 | 0.302333333 | 0.476 | 0.446 | 0.427 | 0.449666667 | 1.487320838 |
| Q22866 | lev-11 tmy-1 Y105E8B.1 | 0.306 | 0.296 | 0.312 | 0.304666667 | 0.454 | 0.454 | 0.452 | 0.453333333 | 1.487964989 |
| P10567 | unc-15 F07A5.7 | 0.29 | 0.294 | 0.302 | 0.295333333 | 0.441 | 0.444 | 0.434 | 0.439666667 | 1.488713318 |
| Q95005 | pas-4 C36B1.4 | 0.297 | 0.296 | 0.296 | 0.296333333 | 0.45 | 0.442 | 0.432 | 0.441333333 | 1.489313836 |
| Q9NAI5 | CELE_Y39G8B.1 Y39G8B.1 | 0.307 | 0.308 | 0.285 | 0.3 | 0.43 | 0.462 | 0.449 | 0.447 | 1.49 |
| Q19626 | vha-12 F20B6.2 | 0.304 | 0.289 | 0.313 | 0.302 | 0.461 | 0.438 | 0.452 | 0.450333333 | 1.491169978 |
| P52275 | tbb-2 C36E8.5 | 0.313 | 0.297 | 0.307 | 0.305666667 | 0.451 | 0.461 | 0.456 | 0.456 | 1.491821156 |
| P34575 | cts-1 T20G5.2 | 0.29 | 0.287 | 0.291 | 0.289333333 | 0.437 | 0.411 | 0.447 | 0.431666667 | 1.491935484 |
| Q19057 | acdh-12 CELE_E04F6.5 E04F6.5 | 0.287 | 0.398 | 0.327 | 0.337333333 | 0.552 | 0.454 | 0.504 | 0.503333333 | 1.492094862 |
| Q11176 | unc-78 C04F6.4 | 0.316 | 0.298 | 0.302 | 0.305333333 | 0.444 | 0.448 | 0.476 | 0.456 | 1.493449782 |
| Q18864 | sft-4 surf-4 C54H2.5 | 0 | 0.401 | 0 | 0.133666667 | 0 | 0.599 | 0 | 0.199666667 | 1.493765586 |
| P20163 | hsp-4 hsp70d F43E2.8 | 0.294 | 0.289 | 0.3 | 0.294333333 | 0.435 | 0.425 | 0.459 | 0.439666667 | 1.493771234 |
| Q09543 | paa-1 F48E8.5 | 0.369 | 0.374 | 0.184 | 0.309 | 0.475 | 0.418 | 0.492 | 0.461666667 | 1.494066882 |
| Q9U302 | pab-1 CELE_Y106G6H.2 Y106G6H.2 | 0.301 | 0.296 | 0.321 | 0.306 | 0.456 | 0.463 | 0.453 | 0.457333333 | 1.494553377 |
| Q17849 | hsp-25 C09B8.6 CELE_C09B8.6 | 0 | 0.392 | 0.305 | 0.232333333 | 0 | 0.608 | 0.434 | 0.347333333 | 1.494978479 |
| O45148 | dlst-1 CELE_W02F12.5 W02F12.5 | 0.316 | 0.301 | 0.307 | 0.308 | 0.439 | 0.499 | 0.444 | 0.460666667 | 1.495670996 |
| O01974 | eif-3.H C41D11.2 | 0.345 | 0.242 | 0.399 | 0.328666667 | 0.416 | 0.543 | 0.516 | 0.491666667 | 1.495943205 |

| | | | | | | | | | |
|---|---|---|---|---|---|---|---|---|---|
| H9G2P9 | cbs-1 CELE_ZC373.1 ZC373.1 | 0 | 0.296 | 0.331 | 0.209 | 0 | 0.463 | 0.475 | 0.312666667 | 1.496012759 |
| Q21732 | CELE_R04F11.2 R04F11.2 | 0.288 | 0.294 | 0.299 | 0.293666667 | 0.443 | 0.443 | 0.432 | 0.439333333 | 1.496027242 |
| O45319 | tin-13 tim-13 DY3.1 | 0.309 | 0.26 | 0.312 | 0.293666667 | 0.419 | 0.461 | 0.438 | 0.439333333 | 1.496027242 |
| Q23500 | aco-1 gei-22 ZK455.1 | 0.309 | 0.303 | 0.307 | 0.306333333 | 0.466 | 0.448 | 0.462 | 0.458666667 | 1.497279652 |
| Q19853 | CELE_F28B4.3 F28B4.3 | 0.329 | 0.31 | 0.307 | 0.315333333 | 0.453 | 0.473 | 0.491 | 0.472333333 | 1.497885835 |
| Q86S26 | pmp-5 CELE_T10H9.5 T10H9.5 | 0.403 | 0.312 | 0.313 | 0.342666667 | 0.597 | 0.467 | 0.476 | 0.513333333 | 1.498054475 |
| P41932 | par-5 ftt-1 M117.2 | 0.291 | 0.311 | 0.309 | 0.303666667 | 0.453 | 0.453 | 0.459 | 0.455 | 1.498353458 |
| Q20626 | CELE_F49E2.5 F49E2.5 | 0.398 | 0.251 | 0.316 | 0.321666667 | 0.602 | 0.401 | 0.444 | 0.482333333 | 1.499481865 |
| Q20053 | asb-1 CELE_F35G12.10 F35G12.10 | 0.266 | 0.319 | 0.384 | 0.323 | 0.514 | 0.416 | 0.523 | 0.484333333 | 1.499484004 |
| Q9BL03 | CELE_Y54F10AM.5 Y54F10AM.5 | 0.296 | 0.275 | 0.261 | 0.277333333 | 0.428 | 0.37 | 0.45 | 0.416 | 1.5 |
| G5EEK9 | vha-5 CELE_F35H10.4 F35H10.4 | 0.324 | 0.289 | 0 | 0.204333333 | 0.446 | 0.474 | 0 | 0.306666667 | 1.500815661 |
| P34334 | rpl-21 C14B9.7 | 0.319 | 0.295 | 0.302 | 0.305333333 | 0.455 | 0.471 | 0.449 | 0.458333333 | 1.501091703 |
| Q20203 | csq-1 CELE_F40E10.3 F40E10.3 | 0.365 | 0.287 | 0.283 | 0.311666667 | 0.443 | 0.456 | 0.505 | 0.468 | 1.501604278 |
| P49049 | imp-2 T05E11.5 | 0.275 | 0.266 | 0.379 | 0.306666667 | 0.497 | 0.465 | 0.42 | 0.460666667 | 1.502173913 |
| O45218 | ads-1 Y50D7A.7 | 0.289 | 0.282 | 0.303 | 0.291333333 | 0.435 | 0.455 | 0.423 | 0.437666667 | 1.50228833 |
| Q9U241 | CELE_Y56A3A.19 Y56A3A.19 | 0.314 | 0.339 | 0.394 | 0.349 | 0.507 | 0.46 | 0.606 | 0.524333333 | 1.502387775 |
| O01812 | lbp-6 W02D3.5 | 0.316 | 0.289 | 0.312 | 0.305666667 | 0.459 | 0.473 | 0.446 | 0.459333333 | 1.502726281 |
| Q19869 | rpl-26 F28C6.7 | 0.307 | 0.3 | 0.31 | 0.305666667 | 0.451 | 0.452 | 0.475 | 0.459333333 | 1.502726281 |
| Q22038 | rho-1 rhoa Y51H4A.3 | 0.297 | 0.297 | 0.305 | 0.299666667 | 0.457 | 0.447 | 0.447 | 0.450333333 | 1.502780868 |
| Q9N5E4 | CELE_T02H6.11 T02H6.11 | 0.361 | 0.298 | 0.279 | 0.312666667 | 0.459 | 0.467 | 0.484 | 0.47 | 1.503198294 |
| P91477 | pbs-4 T20F5.2 | 0.346 | 0 | 0.27 | 0.205333333 | 0.497 | 0 | 0.429 | 0.308666667 | 1.503246753 |
| O17759 | tkt-1 CELE_F01G10.1 F01G10.1 | 0.279 | 0.324 | 0.309 | 0.304 | 0.452 | 0.468 | 0.451 | 0.457 | 1.503289474 |
| Q19007 | nap-1 CELE_D2096.8 D2096.8 | 0.296 | 0.302 | 0.297 | 0.298333333 | 0.453 | 0.447 | 0.446 | 0.448666667 | 1.503910615 |
| P34559 | ech-6 T05G5.6 | 0.287 | 0.292 | 0.3 | 0.293 | 0.439 | 0.447 | 0.436 | 0.440666667 | 1.503981797 |
| G4SPY0 | CELE_Y71G10AL.1 Y71G10AL.1 | 0 | 0.317 | 0 | 0.105666667 | 0 | 0.477 | 0 | 0.159 | 1.504731861 |
| P50140 | hsp-60 hsp60 Y22D7AL.5 | 0.301 | 0.292 | 0.304 | 0.299 | 0.444 | 0.458 | 0.448 | 0.45 | 1.505016722 |
| O01504 | rpa-2 C37A2.7 | 0.307 | 0.283 | 0.303 | 0.297666667 | 0.445 | 0.451 | 0.448 | 0.448 | 1.505039194 |
| Q9TXI4 | CELE_F23C8.5 F23C8.5 | 0.299 | 0.284 | 0.302 | 0.295 | 0.436 | 0.459 | 0.437 | 0.444 | 1.505084746 |
| O01542 | cpn-3 CELE_F28H1.2 F28H1.2 | 0.29 | 0.295 | 0.314 | 0.299666667 | 0.46 | 0.451 | 0.443 | 0.451333333 | 1.506117909 |
| Q9XWS4 | rpl-30 CELE_Y106G6H.3 | 0.317 | 0.314 | 0.303 | 0.311333333 | 0.468 | 0.469 | 0.471 | 0.469333333 | 1.507494647 |
| P54216 | aldo-1 T05D4.1 | 0.312 | 0.279 | 0.299 | 0.296666667 | 0.441 | 0.446 | 0.455 | 0.447333333 | 1.507865169 |
| P52819 | rpl-22 C27A2.2 | 0.301 | 0.294 | 0.308 | 0.301 | 0.45 | 0.455 | 0.457 | 0.454 | 1.508305648 |
| Q22021 | R53.4 | 0.362 | 0.288 | 0.277 | 0.309 | 0.514 | 0.451 | 0.434 | 0.466333333 | 1.509169364 |
| Q22620 | pars-1 CELE_T20H4.3 T20H4.3 | 0.278 | 0.296 | 0.298 | 0.290666667 | 0.465 | 0.436 | 0.415 | 0.438666667 | 1.509174312 |
| P51403 | rps-2 C49H3.11 | 0.306 | 0.307 | 0.294 | 0.302333333 | 0.456 | 0.45 | 0.463 | 0.456333333 | 1.509371555 |

| | | | | | | | | | | |
|---|---|---|---|---|---|---|---|---|---|---|
| O02286 | pck-2 CELE_R11A5.4 R11A5.4 | 0.298 | 0.292 | 0.291 | 0.293666667 | 0.447 | 0.454 | 0.429 | 0.443333333 | 1.509648127 |
| Q21067 | ifc-2 M6.1 | 0.356 | 0.3 | 0.295 | 0.317 | 0.513 | 0.448 | 0.475 | 0.478666667 | 1.509989485 |
| Q93615 | F27D4.1 | 0.288 | 0.285 | 0.301 | 0.291333333 | 0.437 | 0.452 | 0.432 | 0.440333333 | 1.511441648 |
| H2KZG6 | acdh-1 C55B7.4 CELE_C55B7.4 | 0.294 | 0.304 | 0.301 | 0.299666667 | 0.454 | 0.456 | 0.449 | 0.453 | 1.511679644 |
| Q93573 | tct-1 F25H2.11 | 0.302 | 0.297 | 0.289 | 0.296 | 0.45 | 0.449 | 0.444 | 0.447666667 | 1.512387387 |
| Q9U2X0 | prmt-1 CELE_Y113G7B.17 Y113G7B.17 | 0.288 | 0.287 | 0.295 | 0.29 | 0.441 | 0.447 | 0.428 | 0.438666667 | 1.512643678 |
| P18948 | vit-6 K07H8.6 | 0.275 | 0.27 | 0.282 | 0.275666667 | 0.415 | 0.42 | 0.416 | 0.417 | 1.512696493 |
| O01806 | C44E4.4 CELE_C44E4.4 | 0.304 | 0.242 | 0.329 | 0.291666667 | 0.452 | 0.441 | 0.431 | 0.441333333 | 1.513142857 |
| P34382 | far-1 F02A9.2 | 0.294 | 0.281 | 0.298 | 0.291 | 0.437 | 0.439 | 0.445 | 0.440333333 | 1.513172967 |
| Q9N4H7 | copa-1 CELE_Y71F9AL.17 Y71F9AL.17 | 0.302 | 0.307 | 0.271 | 0.293333333 | 0.472 | 0.449 | 0.411 | 0.444 | 1.513636364 |
| Q20507 | acbp-3 F47B10.7 | 0.291 | 0.289 | 0.308 | 0.296 | 0.478 | 0.436 | 0.431 | 0.448333333 | 1.51463964 |
| P29691 | eef-2 F25H5.4 | 0.301 | 0.297 | 0.297 | 0.298333333 | 0.454 | 0.458 | 0.444 | 0.452 | 1.515083799 |
| Q9N3D9 | nduf-5 CELE_Y54E10BL.5 Y54E10BL.5 | 0.412 | 0.227 | 0.244 | 0.294333333 | 0.36 | 0.472 | 0.506 | 0.446 | 1.515288788 |
| Q93576 | ndk-1 CELE_F25H2.5 F25H2.5 | 0.279 | 0.283 | 0.289 | 0.283666667 | 0.439 | 0.42 | 0.431 | 0.43 | 1.51586369 |
| Q21233 | nuo-4 CELE_K04G7.4 K04G7.4 | 0.315 | 0.277 | 0.313 | 0.301666667 | 0.561 | 0.369 | 0.444 | 0.458 | 1.518232044 |
| O17643 | idh-2 C34F6.8 CELE_C34F6.8 | 0 | 0.397 | 0 | 0.132333333 | 0 | 0.603 | 0 | 0.201 | 1.518891688 |
| Q22100 | kat-1 T02G5.8 | 0.307 | 0.264 | 0.369 | 0.313333333 | 0.536 | 0.477 | 0.415 | 0.476 | 1.519148936 |
| G5EC91 | dpy-11 CELE_F46E10.9 F46E10.9 | 0.307 | 0.301 | 0.268 | 0.292 | 0.433 | 0.447 | 0.451 | 0.443666667 | 1.519406393 |
| P53588 | suca-1 F47B10.1 | 0.323 | 0.309 | 0.283 | 0.305 | 0.464 | 0.504 | 0.424 | 0.464 | 1.521311475 |
| O17214 | fum-1 H14A12.2 | 0.296 | 0.32 | 0.281 | 0.299 | 0.458 | 0.438 | 0.469 | 0.455 | 1.52173913 |
| Q9UAV5 | mdh-1 CELE_F46E10.10 F46E10.10 | 0.303 | 0.3 | 0.291 | 0.298 | 0.444 | 0.456 | 0.462 | 0.454 | 1.523489933 |
| Q21193 | pfn-3 K03E6.6 | 0 | 0 | 0.298 | 0.099333333 | 0 | 0 | 0.454 | 0.151333333 | 1.523489933 |
| O17536 | hil-4 C18G1.5 | 0.298 | 0.295 | 0.341 | 0.311333333 | 0.489 | 0.481 | 0.453 | 0.474333333 | 1.523554604 |
| G5ED07 | pdi-3 CELE_H06O01.1 H06O01.1 | 0.296 | 0.304 | 0.298 | 0.299333333 | 0.447 | 0.458 | 0.464 | 0.456333333 | 1.524498886 |
| Q9N3X2 | rps-4 Y43B11AR.4 | 0.276 | 0.284 | 0.309 | 0.289666667 | 0.446 | 0.441 | 0.438 | 0.441666667 | 1.524741082 |
| Q20719 | F53F4.10 | 0.309 | 0.287 | 0.284 | 0.293333333 | 0.421 | 0.485 | 0.437 | 0.447666667 | 1.526136364 |
| O62277 | dct-18 CELE_F58G1.4 F58G1.4 | 0.29 | 0.286 | 0.279 | 0.285 | 0.449 | 0.427 | 0.43 | 0.435333333 | 1.52748538 |
| Q27371 | mup-2 T22E5.5 | 0.286 | 0.281 | 0.35 | 0.305666667 | 0.484 | 0.474 | 0.443 | 0.467 | 1.52780807 |
| O17271 | heh-1 R148.6 | 0.304 | 0.384 | 0.333 | 0.340333333 | 0.42 | 0.639 | 0.501 | 0.52 | 1.52791381 |
| H2KYR1 | vig-1 CELE_F56D12.5 F56D12.5 | 0.306 | 0.281 | 0.314 | 0.300333333 | 0.452 | 0.477 | 0.448 | 0.459 | 1.528301887 |
| P27420 | hsp-3 hsp70c C15H9.6 | 0.316 | 0.294 | 0.292 | 0.300666667 | 0.455 | 0.454 | 0.47 | 0.459666667 | 1.528824834 |
| G1K0V8 | iff-1 CELE_T05G5.10 T05G5.10 | 0.283 | 0.301 | 0 | 0.194666667 | 0.423 | 0.471 | 0 | 0.298 | 1.530821918 |

| | | | | | | | | | | |
|---|---|---|---|---|---|---|---|---|---|---|
| O62146 | F09B12.3 | 0 | 0.302 | 0.252 | 0.184666667 | 0 | 0.5 | 0.349 | 0.283 | 1.532490975 |
| G5EC31 | alh-6 CELE_F56D12.1 F56D12.1 | 0.313 | 0.307 | 0.289 | 0.303 | 0.472 | 0.474 | 0.448 | 0.464666667 | 1.533553355 |
| Q18421 | C34C12.8 | 0.284 | 0.237 | 0.339 | 0.286666667 | 0.468 | 0.46 | 0.391 | 0.439666667 | 1.53372093 |
| Q18688 | daf-21 C47E8.5 | 0.302 | 0.297 | 0.304 | 0.301 | 0.463 | 0.467 | 0.455 | 0.461666667 | 1.533776301 |
| Q21465 | zig-12 CELE_M02D8.1 M02D8.1 | 0 | 0.276 | 0.327 | 0.201 | 0 | 0.464 | 0.461 | 0.308333333 | 1.533996683 |
| O01868 | rpl-24.1 D1007.12 | 0.283 | 0.285 | 0.284 | 0.284 | 0.428 | 0.441 | 0.439 | 0.436 | 1.535211268 |
| H2KYQ5 | gyg-1 F56B6.4 | 0.308 | 0 | 0 | 0.102666667 | 0.473 | 0 | 0 | 0.157666667 | 1.535714286 |
| Q966C7 | tald-1 CELE_Y24D9A.8 Y24D9A.8 | 0.309 | 0.287 | 0.286 | 0.294 | 0.448 | 0.452 | 0.455 | 0.451666667 | 1.536281179 |
| Q9XW92 | vha-13 Y49A3A.2 | 0.299 | 0.286 | 0.288 | 0.291 | 0.449 | 0.447 | 0.446 | 0.447333333 | 1.53722795 |
| Q20780 | alh-1 CELE_F54D8.3 F54D8.3 | 0.285 | 0.306 | 0.294 | 0.295 | 0.455 | 0.454 | 0.452 | 0.453666667 | 1.537853107 |
| Q9GYK2 | C05D9.3 | 0.394 | 0 | 0 | 0.131333333 | 0.606 | 0 | 0 | 0.202 | 1.538071066 |
| H2KZS2 | CELE_T23E7.2 T23E7.2 | 0.322 | 0.284 | 0.258 | 0.288 | 0.425 | 0.465 | 0.44 | 0.443333333 | 1.539351852 |
| Q8MNU8 | C29E4.12 | 0.274 | 0.321 | 0 | 0.198333333 | 0.483 | 0.433 | 0 | 0.305333333 | 1.539495798 |
| O45444 | clec-63 CELE_F35C5.6 F35C5.6 | 0.292 | 0.278 | 0.309 | 0.293 | 0.481 | 0.422 | 0.451 | 0.451333333 | 1.540386803 |
| Q9GRZ9 | tcc-1 CELE_Y59A8A.3 Y59A8A.3 | 0.208 | 0 | 0.26 | 0.156 | 0.384 | 0 | 0.337 | 0.240333333 | 1.540598291 |
| O16462 | grd-5 CELE_F41E6.2 F41E6.2 | 0.283 | 0.306 | 0.271 | 0.286666667 | 0.451 | 0.419 | 0.455 | 0.441666667 | 1.540697674 |
| P27798 | crt-1 Y38A10A.5 | 0.3 | 0.29 | 0.29 | 0.293333333 | 0.439 | 0.454 | 0.463 | 0.452 | 1.540909091 |
| O17732 | pyc-1 D2023.2 | 0.28 | 0.267 | 0.329 | 0.292 | 0.53 | 0.397 | 0.423 | 0.45 | 1.54109589 |
| P91020 | C07D8.6 CELE_C07D8.6 | 0.309 | 0.266 | 0.301 | 0.292 | 0.434 | 0.448 | 0.469 | 0.450333333 | 1.542237443 |
| Q17967 | pdi-1 C14B1.1 | 0.291 | 0.284 | 0.291 | 0.288666667 | 0.446 | 0.446 | 0.445 | 0.445666667 | 1.543879908 |
| Q65ZK0 | rars-1 CELE_F26F4.10 F26F4.10 | 0.271 | 0.305 | 0.336 | 0.304 | 0.525 | 0.456 | 0.428 | 0.469666667 | 1.54495614 |
| Q19289 | ifb-1 F10C1.2 | 0.295 | 0.303 | 0.299 | 0.299 | 0.458 | 0.469 | 0.459 | 0.462 | 1.545150502 |
| P17329 | gpd-2 K10B3.8 | 0.284 | 0.309 | 0.298 | 0.297 | 0.47 | 0.458 | 0.45 | 0.459333333 | 1.54657688 |
| O17626 | C31C9.2 CELE_C31C9.2 | 0.252 | 0.321 | 0.283 | 0.285333333 | 0.446 | 0.44 | 0.438 | 0.441333333 | 1.546728972 |
| Q94269 | CELE_K10C2.1 K10C2.1 | 0.319 | 0.396 | 0.303 | 0.339333333 | 0.461 | 0.604 | 0.51 | 0.525 | 1.547151277 |
| Q86S66 | icd-2 Y65B4BR.5 | 0.302 | 0.286 | 0.307 | 0.298333333 | 0.463 | 0.449 | 0.473 | 0.461666667 | 1.547486034 |
| P52713 | alh-8 F13D12.4 | 0.304 | 0.292 | 0.295 | 0.297 | 0.45 | 0.471 | 0.458 | 0.459666667 | 1.547699214 |
| P27604 | ahcy-1 ahh dpy-14 K02F2.2 | 0.293 | 0.292 | 0.293 | 0.292666667 | 0.456 | 0.455 | 0.448 | 0.453 | 1.547835991 |
| Q23315 | ears-1 CELE_ZC434.5 ZC434.5 | 0.288 | 0.359 | 0.274 | 0.307 | 0.531 | 0.415 | 0.48 | 0.475333333 | 1.548317047 |
| Q95Y90 | rpl-9 R13A5.8 | 0.293 | 0.301 | 0.283 | 0.292333333 | 0.449 | 0.455 | 0.454 | 0.452666667 | 1.548460661 |
| O44989 | col-49 CELE_K09H9.3 K09H9.3 | 0.2 | 0.106 | 0.151 | 0.152333333 | 0.223 | 0.235 | 0.25 | 0.236 | 1.549234136 |
| Q20412 | F44G4.2 | 0 | 0.27 | 0.307 | 0.192333333 | 0 | 0.442 | 0.452 | 0.298 | 1.549393414 |
| O45293 | gly-8 Y66A7A.6 | 0 | 0 | 0.302 | 0.100666667 | 0 | 0 | 0.468 | 0.156 | 1.549668874 |
| G5EDZ9 | cpi-1 CELE_K08B4.6 K08B4.6 | 0.392 | 0 | 0 | 0.130666667 | 0.608 | 0 | 0 | 0.202666667 | 1.551020408 |
| Q18803 | asg-2 C53B7.4 | 0.287 | 0.299 | 0.301 | 0.295666667 | 0.489 | 0.469 | 0.418 | 0.458666667 | 1.551296505 |

| | | | | | | | | | | |
|---|---|---|---|---|---|---|---|---|---|---|
| O17680 | sams-1 C49F5.1 | 0.304 | 0.296 | 0.306 | 0.302 | 0.47 | 0.475 | 0.461 | 0.468666667 | 1.55187638 |
| O45622 | erfa-3 CELE_H19N07.1 H19N07.1 | 0.303 | 0.289 | 0.285 | 0.292333333 | 0.461 | 0.467 | 0.435 | 0.454333333 | 1.554161916 |
| Q18787 | rpt-1 C52E4.4 | 0 | 0.27 | 0.3 | 0.19 | 0 | 0.452 | 0.434 | 0.295333333 | 1.554385965 |
| Q05036 | hsp-110 C30C11.4 | 0.303 | 0.304 | 0.291 | 0.299333333 | 0.455 | 0.472 | 0.469 | 0.465333333 | 1.554565702 |
| Q9NEW6 | rsp-3 Y111B2A.18 | 0.237 | 0.308 | 0.22 | 0.255 | 0.408 | 0.416 | 0.366 | 0.396666667 | 1.555555556 |
| Q17761 | T25B9.9 | 0.325 | 0.265 | 0.298 | 0.296 | 0.452 | 0.478 | 0.453 | 0.461 | 1.557432432 |
| P46561 | atp-2 C34E10.6 | 0.299 | 0.274 | 0.284 | 0.285666667 | 0.444 | 0.452 | 0.439 | 0.445 | 1.557759627 |
| O44156 | pas-6 CD4.6 | 0.283 | 0.286 | 0.29 | 0.286333333 | 0.421 | 0.473 | 0.445 | 0.446333333 | 1.55878929 |
| Q09261 | mrps-31 C32A3.2 | 0.288 | 0 | 0 | 0.096 | 0.449 | 0 | 0 | 0.149666667 | 1.559027778 |
| Q2L6Y6 | hsp-75 CELE_R151.7 R151.7 | 0.288 | 0 | 0 | 0.096 | 0.449 | 0 | 0 | 0.149666667 | 1.559027778 |
| Q21752 | vdac-1 R05G6.7 | 0.296 | 0.297 | 0.29 | 0.294333333 | 0.462 | 0.453 | 0.462 | 0.459 | 1.559456399 |
| G5EEE5 | elo-1 ceelo1 CELE_F56H11.4 F56H11.4 | 0.285 | 0.3 | 0.298 | 0.294333333 | 0.445 | 0.48 | 0.453 | 0.459333333 | 1.560588901 |
| O44480 | rpl-20 E04A4.8 | 0.279 | 0.289 | 0.313 | 0.293666667 | 0.453 | 0.47 | 0.452 | 0.458333333 | 1.560726447 |
| P04255 | his-11 ZK131.5; his-15 ZK131.9; his-29 F35H10.11; his-34 F17E9.9; his-44 F08G2.1 | 0.292 | 0.281 | 0.295 | 0.289333333 | 0.448 | 0.46 | 0.447 | 0.451666667 | 1.561059908 |
| Q21215 | rack-1 K04D7.1 | 0.281 | 0.298 | 0.302 | 0.293666667 | 0.461 | 0.458 | 0.457 | 0.458666667 | 1.561861521 |
| P41938 | B0272.3 | 0.293 | 0.244 | 0.302 | 0.279666667 | 0.414 | 0.413 | 0.484 | 0.437 | 1.562574493 |
| Q20206 | rps-11 CELE_F40F11.1 F40F11.1 | 0.291 | 0.287 | 0.281 | 0.286333333 | 0.456 | 0.444 | 0.443 | 0.447666667 | 1.563445867 |
| Q9GZE9 | ldp-1 CELE_F22F7.1 F22F7.1 | 0.32 | 0.25 | 0.311 | 0.293666667 | 0.412 | 0.556 | 0.412 | 0.46 | 1.566401816 |
| Q95YD5 | vha-16 C30F8.2 CELE_C30F8.2 | 0.272 | 0.26 | 0.333 | 0.288333333 | 0.437 | 0.455 | 0.463 | 0.451666667 | 1.566473988 |
| O16264 | F40A3.3 | 0.319 | 0.296 | 0.306 | 0.307 | 0.441 | 0.567 | 0.435 | 0.481 | 1.566775244 |
| Q9XUS5 | CELE_K08E3.5 K08E3.5 | 0.231 | 0 | 0 | 0.077 | 0.362 | 0 | 0 | 0.120666667 | 1.567099567 |
| Q20228 | rps-9 F40F8.10 | 0.29 | 0.299 | 0.289 | 0.292666667 | 0.467 | 0.451 | 0.458 | 0.458666667 | 1.567198178 |
| Q9N599 | pas-3 Y110A7A.14 | 0.309 | 0.319 | 0.287 | 0.305 | 0.448 | 0.444 | 0.542 | 0.478 | 1.567213115 |
| O76840 | mig-6 ppn-1 C37C3.6 | 0.279 | 0.28 | 0.29 | 0.283 | 0.465 | 0.429 | 0.438 | 0.444 | 1.568904594 |
| O17218 | rps-22 CELE_F53A3.3 F53A3.3 | 0.317 | 0.295 | 0.286 | 0.299333333 | 0.475 | 0.461 | 0.473 | 0.469666667 | 1.569042316 |
| P91374 | rpl-15 K11H12.2 | 0.294 | 0.306 | 0.289 | 0.296333333 | 0.471 | 0.455 | 0.469 | 0.465 | 1.569178853 |
| O44906 | pck-1 CELE_W05G11.6 W05G11.6 | 0.282 | 0.247 | 0.3 | 0.276333333 | 0.397 | 0.447 | 0.457 | 0.433666667 | 1.569360676 |
| O01761 | unc-89 C09D1.1 | 0.296 | 0.28 | 0.288 | 0.288 | 0.438 | 0.464 | 0.454 | 0.452 | 1.569444444 |
| O02158 | CELE_T09B4.8 T09B4.8 | 0 | 0.295 | 0 | 0.098333333 | 0 | 0.463 | 0 | 0.154333333 | 1.569491525 |
| H2KYJ5 | mtch-1 CELE_F43E2.7 F43E2.7 | 0.24 | 0.365 | 0.329 | 0.311333333 | 0.508 | 0.519 | 0.44 | 0.489 | 1.570663812 |
| P52717 | F41C3.5 | 0.29 | 0.286 | 0.279 | 0.285 | 0.46 | 0.444 | 0.439 | 0.447666667 | 1.570760234 |
| P91913 | rla-1 rpa-1 Y37E3.7 | 0.302 | 0.292 | 0.28 | 0.291333333 | 0.457 | 0.451 | 0.465 | 0.457666667 | 1.570938215 |

| | | | | | | | | | | |
|---|---|---|---|---|---|---|---|---|---|---|
| P90868 | pbs-7 CELE_F39H11.5 F39H11.5 | 0.249 | 0 | 0.321 | 0.19 | 0.452 | 0 | 0.444 | 0.298666667 | 1.571929825 |
| O45865 | ant-1.1 CELE_T27E9.1 T27E9.1 | 0.286 | 0.283 | 0.298 | 0.289 | 0.45 | 0.46 | 0.453 | 0.454333333 | 1.572087659 |
| Q95QW0 | eif-3.L C17G10.9 | 0 | 0.311 | 0.311 | 0.207333333 | 0 | 0.521 | 0.457 | 0.326 | 1.572347267 |
| Q9U1X9 | rla-2 CELE_Y62E10A.1 Y62E10A.1 | 0.266 | 0.274 | 0.286 | 0.275333333 | 0.421 | 0.432 | 0.446 | 0.433 | 1.572639225 |
| O16309 | fkb-3 C05C8.3 CELE_C05C8.3 | 0.289 | 0.244 | 0.28 | 0.271 | 0.436 | 0.383 | 0.46 | 0.426333333 | 1.573185732 |
| Q9U1Q4 | vrs-2 Y87G2A.5 | 0.286 | 0.307 | 0.293 | 0.295333333 | 0.455 | 0.474 | 0.466 | 0.465 | 1.574492099 |
| P48156 | rps-8 F42C5.8 | 0.284 | 0.279 | 0.299 | 0.287333333 | 0.445 | 0.461 | 0.452 | 0.452666667 | 1.575406032 |
| Q9N4I4 | rpl-10a rpl-1 Y71F9AL.13 | 0.28 | 0.29 | 0.281 | 0.283666667 | 0.439 | 0.449 | 0.453 | 0.447 | 1.575793184 |
| Q27487 | ctl-2 cat cat-1 Y54G11A.5 | 0.274 | 0.238 | 0.276 | 0.262666667 | 0.444 | 0.384 | 0.414 | 0.414 | 1.576142132 |
| P34496 | mtss-1 PAR2.1 | 0.302 | 0 | 0 | 0.100666667 | 0.476 | 0 | 0 | 0.158666667 | 1.57615894 |
| P46769 | rps-0 B0393.1 | 0.297 | 0.288 | 0.293 | 0.292666667 | 0.462 | 0.461 | 0.461 | 0.461333333 | 1.576309795 |
| O45946 | rpl-18 Y45F10D.12 | 0.293 | 0.275 | 0.289 | 0.285666667 | 0.452 | 0.449 | 0.45 | 0.450333333 | 1.576429405 |
| O17586 | pas-1 C15H11.7 | 0.32 | 0.289 | 0.259 | 0.289333333 | 0.462 | 0.438 | 0.469 | 0.456333333 | 1.57718894 |
| O45499 | rps-26 F39B2.6 | 0.288 | 0.279 | 0.294 | 0.287 | 0.447 | 0.456 | 0.455 | 0.452666667 | 1.577235772 |
| P48162 | rpl-25.1 F55D10.2 | 0.294 | 0.308 | 0.296 | 0.299333333 | 0.475 | 0.466 | 0.476 | 0.472333333 | 1.577951002 |
| P91917 | ola-1 tag-210 W08E3.3 | 0.276 | 0.267 | 0.303 | 0.282 | 0.456 | 0.439 | 0.44 | 0.445 | 1.578014184 |
| O02266 | alh-7 CELE_F45H10.1 F45H10.1 | 0.294 | 0 | 0 | 0.098 | 0.464 | 0 | 0 | 0.154666667 | 1.578231293 |
| Q09581 | lec-3 ZK892.1 | 0 | 0.336 | 0.267 | 0.201 | 0 | 0.467 | 0.485 | 0.317333333 | 1.578772803 |
| P34383 | far-2 F02A9.3 | 0.265 | 0.29 | 0.274 | 0.276333333 | 0.429 | 0.438 | 0.442 | 0.436333333 | 1.579010856 |
| Q27493 | rpoa-2 CELE_F14B4.3 F14B4.3 | 0.297 | 0 | 0 | 0.099 | 0.469 | 0 | 0 | 0.156333333 | 1.579124579 |
| P34500 | ttr-2 K03H1.4 | 0.287 | 0.304 | 0.306 | 0.299 | 0.495 | 0.455 | 0.467 | 0.472333333 | 1.579710145 |
| G5EEA8 | nex-1 NEX-1 CELE_ZC155.1 ZC155.1 | 0.287 | 0.292 | 0.293 | 0.290666667 | 0.452 | 0.473 | 0.453 | 0.459333333 | 1.580275229 |
| O44750 | app-1 W03G9.4 | 0.259 | 0 | 0.32 | 0.193 | 0.469 | 0 | 0.446 | 0.305 | 1.580310881 |
| O02639 | rpl-19 C09D4.5 | 0.317 | 0.278 | 0.288 | 0.294333333 | 0.43 | 0.445 | 0.521 | 0.465333333 | 1.580973952 |
| O17528 | sel-9 W02D7.7 | 0.303 | 0.269 | 0.277 | 0.283 | 0.454 | 0.435 | 0.454 | 0.447666667 | 1.581861013 |
| Q9N5V3 | imb-3 C53D5.6 CELE_C53D5.6 | 0.272 | 0.302 | 0.298 | 0.290666667 | 0.453 | 0.465 | 0.462 | 0.46 | 1.582568807 |
| Q9BKP8 | C17F4.7 CELE_C17F4.7 | 0.283 | 0.311 | 0.269 | 0.287666667 | 0.449 | 0.448 | 0.469 | 0.455333333 | 1.582850521 |
| Q17993 | C14F11.6 CELE_C14F11.6 | 0 | 0.307 | 0 | 0.102333333 | 0 | 0.486 | 0 | 0.162 | 1.583061889 |
| P34662 | rpl-35 ZK652.4 | 0.262 | 0.293 | 0.285 | 0.28 | 0.451 | 0.432 | 0.447 | 0.443333333 | 1.583333333 |
| P41994 | rpia-1 B0280.3 | 0.274 | 0.275 | 0.268 | 0.272333333 | 0.429 | 0.445 | 0.42 | 0.431333333 | 1.583843329 |
| O76449 | CELE_ZK1055.7 ZK1055.7 | 0.323 | 0.285 | 0 | 0.202666667 | 0.417 | 0.546 | 0 | 0.321 | 1.583881579 |
| Q9GYF1 | unc-27 tni-2 ZK721.2 | 0.299 | 0.28 | 0.305 | 0.294666667 | 0.456 | 0.479 | 0.466 | 0.467 | 1.584841629 |
| Q21926 | irs-1 R11A8.6 | 0.294 | 0.299 | 0.301 | 0.298 | 0.466 | 0.481 | 0.47 | 0.472333333 | 1.585011186 |
| P48152 | rps-3 C23G10.3 | 0.288 | 0.288 | 0.29 | 0.288666667 | 0.454 | 0.468 | 0.451 | 0.457666667 | 1.585450346 |

| | | | | | | | | | | |
|---|---|---|---|---|---|---|---|---|---|---|
| Q22370 | ucr-2.2 CELE_T10B10.2 T10B10.2 | 0.263 | 0.291 | 0.276 | 0.276666667 | 0.437 | 0.442 | 0.438 | 0.439 | 1.586746988 |
| Q19246 | dhs-25 CELE_F09E10.3 F09E10.3 | 0.293 | 0.256 | 0.338 | 0.295666667 | 0.447 | 0.504 | 0.457 | 0.469333333 | 1.587373168 |
| G5EEI4 | asp-1 CELE_Y39B6A.20 Y39B6A.20 | 0.244 | 0.241 | 0.236 | 0.240333333 | 0.384 | 0.388 | 0.375 | 0.382333333 | 1.590846047 |
| P34690 | tba-2 C47B2.3 | 0.275 | 0.272 | 0.276 | 0.274333333 | 0.441 | 0.438 | 0.431 | 0.436666667 | 1.591737546 |
| Q23588 | upp-1 ZK783.2 | 0.25 | 0 | 0 | 0.083333333 | 0.398 | 0 | 0 | 0.132666667 | 1.592 |
| Q7Z072 | tnt-2 CELE_F53A9.10 F53A9.10 | 0.287 | 0.288 | 0.286 | 0.287 | 0.457 | 0.455 | 0.459 | 0.457 | 1.592334495 |
| P37806 | unc-87 F08B6.4 | 0.284 | 0.3 | 0.282 | 0.288666667 | 0.476 | 0.444 | 0.459 | 0.459666667 | 1.592378753 |
| Q8WSN3 | spp-17 C54G6.5 CELE_C54G6.5 | 0 | 0 | 0.305 | 0.101666667 | 0 | 0 | 0.486 | 0.162 | 1.593442623 |
| O18000 | pes-9 CELE_R11H6.1 R11H6.1 | 0.282 | 0.278 | 0.301 | 0.287 | 0.473 | 0.452 | 0.447 | 0.457333333 | 1.593495935 |
| O45012 | nol-5 CELE_W01B11.3 W01B11.3 | 0.3 | 0.277 | 0.304 | 0.293666667 | 0.477 | 0.47 | 0.458 | 0.468333333 | 1.594778661 |
| Q93831 | CELE_F59C6.5 F59C6.5 | 0.279 | 0.268 | 0.306 | 0.284333333 | 0.468 | 0.47 | 0.423 | 0.453666667 | 1.595545135 |
| Q93619 | tag-173 CELE_F27D4.5 F27D4.5 | 0.274 | 0.248 | 0.288 | 0.27 | 0.403 | 0.342 | 0.548 | 0.431 | 1.596296296 |
| O17919 | K01G5.5 | 0.269 | 0.296 | 0.302 | 0.289 | 0.455 | 0.466 | 0.464 | 0.461666667 | 1.597462514 |
| Q93934 | CELE_R07H5.8 R07H5.8 | 0.289 | 0.295 | 0.286 | 0.29 | 0.471 | 0.464 | 0.455 | 0.463333333 | 1.597701149 |
| P53589 | sucg-1 C50F7.4 | 0.303 | 0.28 | 0.297 | 0.293333333 | 0.481 | 0.453 | 0.472 | 0.468666667 | 1.597727273 |
| Q9N5S7 | CELE_F49H12.5 F49H12.5 | 0.291 | 0.258 | 0.295 | 0.281333333 | 0.457 | 0.475 | 0.417 | 0.449666667 | 1.598341232 |
| P12844 | myo-3 mhcA K12F2.1 | 0.303 | 0.312 | 0.293 | 0.302666667 | 0.491 | 0.48 | 0.484 | 0.485 | 1.602422907 |
| Q93572 | rpa-0 F25H2.10 | 0.29 | 0.286 | 0.291 | 0.289 | 0.471 | 0.465 | 0.455 | 0.463666667 | 1.60438293 |
| Q17474 | B0334.3 CELE_B0334.3 | 0 | 0.278 | 0.256 | 0.178 | 0 | 0.406 | 0.451 | 0.285666667 | 1.604868914 |
| Q9TXU7 | eif-1.a eif-1.A CELE_H06H21.3 H06H21.3 | 0.297 | 0.293 | 0.269 | 0.286333333 | 0.458 | 0.446 | 0.475 | 0.459666667 | 1.605355064 |
| O44503 | CELE_R02D3.1 R02D3.1 | 0.255 | 0 | 0.27 | 0.175 | 0.384 | 0 | 0.459 | 0.281 | 1.605714286 |
| Q18310 | C29F5.1 CELE_C29F5.1 | 0 | 0.315 | 0.279 | 0.198 | 0 | 0.487 | 0.467 | 0.318 | 1.606060606 |
| Q21735 | ran-4 R05D11.3 | 0.286 | 0 | 0.281 | 0.189 | 0.466 | 0 | 0.445 | 0.303666667 | 1.60670194 |
| Q95Y28 | CELE_Y34B4A.9 Y34B4A.9 | 0 | 0 | 0.28 | 0.093333333 | 0 | 0 | 0.45 | 0.15 | 1.607142857 |
| Q22099 | krs-1 T02G5.9 | 0.286 | 0.275 | 0.272 | 0.277666667 | 0.452 | 0.462 | 0.425 | 0.446333333 | 1.607442977 |
| O01833 | hil-5 B0414.3 | 0.325 | 0.312 | 0.192 | 0.276333333 | 0.412 | 0.42 | 0.501 | 0.444333333 | 1.607961399 |
| Q22562 | CELE_T19B10.2 T19B10.2 | 0.295 | 0.286 | 0.259 | 0.28 | 0.43 | 0.442 | 0.479 | 0.450333333 | 1.608333333 |
| P05690 | vit-2 C42D8.2 | 0.264 | 0.273 | 0.274 | 0.270333333 | 0.431 | 0.439 | 0.435 | 0.435 | 1.609124538 |
| Q21219 | pept-1 cptb opt-2 pep-2 K04E7.2 | 0 | 0.479 | 0 | 0.159666667 | 0 | 0.771 | 0 | 0.257 | 1.60960334 |
| Q9N3H3 | CELE_Y53G8AL.2 Y53G8AL.2 | 0.269 | 0 | 0.294 | 0.187666667 | 0.461 | 0 | 0.446 | 0.302333333 | 1.611012433 |
| O44509 | CELE_F42G8.10 F42G8.10 | 0.265 | 0 | 0 | 0.088333333 | 0.427 | 0 | 0 | 0.142333333 | 1.611320755 |
| Q94261 | cif-1 csn-7 eif-3.M K08F11.3 | 0.278 | 0.259 | 0.347 | 0.294666667 | 0.449 | 0.498 | 0.479 | 0.475333333 | 1.613122172 |
| O45418 | fkb-6 CELE_F31D4.3 F31D4.3 | 0.266 | 0.254 | 0.336 | 0.285333333 | 0.45 | 0.496 | 0.435 | 0.460333333 | 1.613317757 |
| H2L0Q1 | CELE_ZK6.11 ZK6.11 | 0.298 | 0.286 | 0.286 | 0.29 | 0.445 | 0.485 | 0.475 | 0.468333333 | 1.614942529 |

| | | | | | | | | | | |
|---|---|---|---|---|---|---|---|---|---|---|
| O01869 | rps-10 CELE_D1007.6 D1007.6 | 0.299 | 0.288 | 0.286 | 0.291 | 0.483 | 0.471 | 0.456 | 0.47 | 1.615120275 |
| G8JXY2 | mthf-1 C06A8.1 | 0.305 | 0.31 | 0.299 | 0.304666667 | 0.472 | 0.496 | 0.509 | 0.492333333 | 1.615973742 |
| P18947 | vit-4 F59D8.2 | 0.277 | 0.263 | 0.27 | 0.27 | 0.437 | 0.444 | 0.428 | 0.436333333 | 1.616049383 |
| P55326 | F13E6.1 | 0.279 | 0.314 | 0.354 | 0.315666667 | 0.53 | 0.527 | 0.474 | 0.510333333 | 1.616684266 |
| Q17880 | nuo-1 C09H10.3 CELE_C09H10.3 | 0.269 | 0.263 | 0.289 | 0.273666667 | 0.452 | 0.44 | 0.437 | 0.443 | 1.618757613 |
| Q19416 | dylt-1 CELE_F13G3.4 F13G3.4 | 0 | 0 | 0.265 | 0.088333333 | 0 | 0 | 0.429 | 0.143 | 1.618867925 |
| Q18785 | mif-2 C52E4.2 | 0.294 | 0.545 | 0 | 0.279666667 | 0.4 | 0.426 | 0.533 | 0.453 | 1.619785459 |
| Q93168 | C01G10.8 CELE_C01G10.8 | 0.27 | 0.277 | 0.281 | 0.276 | 0.422 | 0.494 | 0.427 | 0.447666667 | 1.621980676 |
| Q09486 | C30G12.2 CELE_C30G12.2 | 0 | 0.279 | 0 | 0.093 | 0 | 0.453 | 0 | 0.151 | 1.623655914 |
| G5EFZ1 | ipgm-1 F57B10.3 | 0.296 | 0.293 | 0.298 | 0.295666667 | 0.457 | 0.467 | 0.517 | 0.480333333 | 1.624577227 |
| Q9XVS4 | dao-5 C25A1.10 CELE_C25A1.10 | 0.381 | 0 | 0 | 0.127 | 0.619 | 0 | 0 | 0.206333333 | 1.624671916 |
| Q20720 | CELE_F53F4.11 F53F4.11 | 0.298 | 0 | 0.267 | 0.188333333 | 0.467 | 0 | 0.451 | 0.306 | 1.624778761 |
| G5EC10 | lec-9 C16H3.2 CELE_C16H3.2 | 0 | 0.284 | 0.28 | 0.188 | 0 | 0.46 | 0.457 | 0.305666667 | 1.625886525 |
| Q9GYT0 | ech-1.2 CELE_T08B2.7 T08B2.7 | 0.297 | 0.293 | 0.26 | 0.283333333 | 0.454 | 0.454 | 0.476 | 0.461333333 | 1.628235294 |
| P09446 | hsp-1 hsp70a F26D10.3 | 0.282 | 0.271 | 0.282 | 0.278333333 | 0.456 | 0.451 | 0.453 | 0.453333333 | 1.628742515 |
| Q18040 | C16A3.10 | 0.286 | 0.283 | 0.285 | 0.284666667 | 0.466 | 0.466 | 0.459 | 0.463666667 | 1.628805621 |
| O17694 | ril-1 C53A5.1 CELE_C53A5.1 | 0 | 0.278 | 0 | 0.092666667 | 0 | 0.453 | 0 | 0.151 | 1.629496403 |
| P47208 | cct-4 K01C8.10 | 0.28 | 0.278 | 0.302 | 0.286666667 | 0.478 | 0.469 | 0.455 | 0.467333333 | 1.630232558 |
| P49596 | ppm-2 T23F11.1 | 0.274 | 0 | 0 | 0.091333333 | 0.447 | 0 | 0 | 0.149 | 1.631386861 |
| O02640 | mdh-2 F20H11.3 | 0.284 | 0.248 | 0.307 | 0.279666667 | 0.457 | 0.462 | 0.45 | 0.456333333 | 1.63170441 |
| Q9N358 | cct-8 Y55F3AR.3 | 0.303 | 0.284 | 0.307 | 0.298 | 0.474 | 0.49 | 0.496 | 0.486666667 | 1.63310962 |
| G5EC23 | hcf-1 C46A5.9 CELE_C46A5.9 | 0.138 | 0 | 0.427 | 0.188333333 | 0.575 | 0 | 0.348 | 0.307666667 | 1.633628319 |
| Q10657 | tpi-1 Y17G7B.7 | 0.286 | 0.29 | 0.285 | 0.287 | 0.482 | 0.464 | 0.461 | 0.469 | 1.634146341 |
| Q9XVE9 | rpl-14 C04F12.4 CELE_C04F12.4 | 0.277 | 0.269 | 0.278 | 0.274666667 | 0.446 | 0.452 | 0.449 | 0.449 | 1.634708738 |
| Q9NF11 | hprt-1 CELE_Y105E8B.5 Y105E8B.5 | 0.311 | 0.3 | 0.32 | 0.310333333 | 0.517 | 0.55 | 0.455 | 0.507333333 | 1.634801289 |
| P49041 | rps-5 T05E11.1 | 0.278 | 0.287 | 0.284 | 0.283 | 0.466 | 0.461 | 0.461 | 0.462666667 | 1.634864547 |
| P54812 | cdc-48.2 C41C4.8 | 0.291 | 0.295 | 0.284 | 0.29 | 0.5 | 0.474 | 0.451 | 0.475 | 1.637931034 |
| O44451 | pdhb-1 C04C3.3 | 0.265 | 0.304 | 0.301 | 0.29 | 0.466 | 0.513 | 0.446 | 0.475 | 1.637931034 |
| Q86FL8 | spp-5 CELE_T08A9.9 T08A9.9 | 0.278 | 0.247 | 0 | 0.175 | 0.428 | 0.432 | 0 | 0.286666667 | 1.638095238 |
| Q19877 | rps-23 F28D1.7 | 0.33 | 0.266 | 0.288 | 0.294666667 | 0.516 | 0.445 | 0.488 | 0.483 | 1.639140271 |
| P41956 | mev-1 cyt-1 T07C4.7 | 0.282 | 0.314 | 0.293 | 0.296333333 | 0.476 | 0.534 | 0.448 | 0.486 | 1.640044994 |
| Q20122 | CELE_F37C12.3 F37C12.3 | 0.264 | 0 | 0 | 0.088 | 0.433 | 0 | 0 | 0.144333333 | 1.640151515 |
| Q09444 | ubh-4 C08B11.7 | 0.308 | 0 | 0.224 | 0.177333333 | 0.459 | 0 | 0.414 | 0.291 | 1.640977444 |
| G5EE04 | hip-1 CELE_T12D8.8 T12D8.8 | 0.284 | 0.338 | 0.293 | 0.305 | 0.487 | 0.515 | 0.5 | 0.500666667 | 1.641530055 |
| G5EDE8 | ile-1 CELE_K07A1.8 K07A1.8 | 0.294 | 0 | 0 | 0.098 | 0.483 | 0 | 0 | 0.161 | 1.642857143 |

| | | | | | | | | | | |
|---|---|---|---|---|---|---|---|---|---|---|
| P47991 | rpl-6 R151.3 | 0.275 | 0.283 | 0.288 | 0.282 | 0.467 | 0.462 | 0.462 | 0.463666667 | 1.644208038 |
| Q21898 | vha-1 R10E11.8 | 0.284 | 0 | 0 | 0.094666667 | 0.467 | 0 | 0 | 0.155666667 | 1.644366197 |
| H2KZL7 | prdx-2 CELE_F09E5.15 F09E5.15 | 0.262 | 0.287 | 0.285 | 0.278 | 0.442 | 0.482 | 0.448 | 0.457333333 | 1.645083933 |
| Q17994 | got-2.2 C14F11.1 CELE_C14F11.1 | 0.284 | 0.288 | 0.294 | 0.288666667 | 0.489 | 0.482 | 0.455 | 0.475333333 | 1.64665127 |
| G5EFI4 | lec-5 CELE_ZK1248.16 ZK1248.16 | 0.272 | 0.254 | 0.309 | 0.278333333 | 0.468 | 0.458 | 0.449 | 0.458333333 | 1.646706587 |
| Q9GRY9 | CELE_Y59A8B.10 Y59A8B.10 | 0 | 0.363 | 0.317 | 0.226666667 | 0 | 0.439 | 0.683 | 0.374 | 1.65 |
| P49405 | rpl-5 F54C9.5 | 0.286 | 0.282 | 0.29 | 0.286 | 0.461 | 0.47 | 0.485 | 0.472 | 1.65034965 |
| Q4TT88 | pam-1 CELE_F49E8.3 F49E8.3 | 0.282 | 0.291 | 0.299 | 0.290666667 | 0.509 | 0.448 | 0.483 | 0.48 | 1.651376147 |
| G4SLH0 | ttn-1 W06H8.8 | 0.279 | 0.299 | 0.26 | 0.279333333 | 0.461 | 0.45 | 0.474 | 0.461666667 | 1.65274463 |
| Q20819 | tsfm-1 F55C5.5 | 0 | 0 | 0.267 | 0.089 | 0 | 0 | 0.442 | 0.147333333 | 1.655430712 |
| P40614 | F01G4.6 | 0.281 | 0.277 | 0.281 | 0.279666667 | 0.465 | 0.469 | 0.456 | 0.463333333 | 1.656734207 |
| Q22799 | dlc-1 T26A5.9 | 0.277 | 0.293 | 0.275 | 0.281666667 | 0.472 | 0.471 | 0.457 | 0.466666667 | 1.656804734 |
| Q95ZS5 | CELE_F56A8.3 F56A8.3 | 0.275 | 0.299 | 0.268 | 0.280666667 | 0.451 | 0.492 | 0.455 | 0.466 | 1.660332542 |
| A7LPE6 | gpdh-2 CELE_K11H3.1 K11H3.1 | 0 | 0.265 | 0 | 0.088333333 | 0 | 0.44 | 0 | 0.146666667 | 1.660377358 |
| Q9U2A8 | rpl-43 rpl-37a Y48B6A.2 | 0.25 | 0.267 | 0.277 | 0.264666667 | 0.444 | 0.441 | 0.434 | 0.439666667 | 1.661209068 |
| Q23312 | rps-7 ZC434.2 | 0.277 | 0.263 | 0.293 | 0.277666667 | 0.46 | 0.463 | 0.461 | 0.461333333 | 1.661464586 |
| Q27488 | pas-2 D1054.2 | 0.283 | 0 | 0.271 | 0.184666667 | 0.476 | 0 | 0.445 | 0.307 | 1.662454874 |
| P34574 | chc-1 T20G5.1 | 0.274 | 0.351 | 0.293 | 0.306 | 0.518 | 0.451 | 0.558 | 0.509 | 1.663398693 |
| Q10121 | C23G10.2 | 0.292 | 0.283 | 0.266 | 0.280333333 | 0.463 | 0.458 | 0.478 | 0.466333333 | 1.663495838 |
| O44411 | nog-1 T07A9.9 | 0 | 0.289 | 0 | 0.096333333 | 0 | 0.481 | 0 | 0.160333333 | 1.664359862 |
| P48154 | rps-1 F56F3.5 | 0.278 | 0.279 | 0.28 | 0.279 | 0.467 | 0.465 | 0.462 | 0.464666667 | 1.665471924 |
| O45525 | CELE_F45H10.2 F45H10.2 | 0.266 | 0.28 | 0.283 | 0.276333333 | 0.455 | 0.446 | 0.48 | 0.460333333 | 1.665862485 |
| Q20865 | CELE_F56C9.7 F56C9.7 | 0.305 | 0.317 | 0.296 | 0.306 | 0.479 | 0.615 | 0.436 | 0.51 | 1.666666667 |
| Q17770 | pdi-2 C07A12.4 | 0.284 | 0.273 | 0.286 | 0.281 | 0.471 | 0.469 | 0.465 | 0.468333333 | 1.666666667 |
| G5EBI0 | 4D656 CELE_Y54G2A.18 Y54G2A.18 | 0.299 | 0.274 | 0.258 | 0.277 | 0.481 | 0.469 | 0.435 | 0.461666667 | 1.666666667 |
| P91133 | ubr-1 C32E8.11 | 0.224 | 0 | 0 | 0.074666667 | 0.374 | 0 | 0 | 0.124666667 | 1.669642857 |
| Q23050 | clik-1 CELE_T25F10.6 T25F10.6 | 0.275 | 0.261 | 0.285 | 0.273666667 | 0.454 | 0.464 | 0.453 | 0.457 | 1.669914738 |
| O18687 | CELE_F26E4.6 F26E4.6 | 0.276 | 0 | 0 | 0.092 | 0.461 | 0 | 0 | 0.153666667 | 1.670289855 |
| O16259 | sti-1 R09E12.3 | 0.277 | 0.293 | 0.285 | 0.285 | 0.482 | 0.487 | 0.462 | 0.477 | 1.673684211 |
| G5EC98 | ctps-1 CELE_W06H3.3 W06H3.3 | 0 | 0.274 | 0.276 | 0.183333333 | 0 | 0.463 | 0.458 | 0.307 | 1.674545455 |
| P43509 | cpr-5 W07B8.5 | 0.301 | 0.276 | 0.244 | 0.273666667 | 0.392 | 0.458 | 0.525 | 0.458333333 | 1.674786845 |
| P46562 | alh-9 F01F1.6 | 0.277 | 0.276 | 0.276 | 0.276333333 | 0.454 | 0.471 | 0.464 | 0.463 | 1.675512666 |
| Q9TZQ3 | pgl-1 ZK381.4 | 0.263 | 0.28 | 0.297 | 0.28 | 0.471 | 0.469 | 0.468 | 0.469333333 | 1.676190476 |
| Q18066 | dim-1 C18A11.7/C18A11.8 | 0.279 | 0.291 | 0.285 | 0.285 | 0.473 | 0.489 | 0.473 | 0.478333333 | 1.678362573 |

| | | | | | | | | | | |
|---|---|---|---|---|---|---|---|---|---|---|
| O17953 | dld-1 LLC1.3 | 0.283 | 0.285 | 0.278 | 0.282 | 0.459 | 0.487 | 0.475 | 0.473666667 | 1.679669031 |
| P50305 | sams-3 C06E7.1 | 0.283 | 0.28 | 0.277 | 0.28 | 0.471 | 0.468 | 0.472 | 0.470333333 | 1.679761905 |
| Q18115 | rpn-2 C23G10.4 | 0.274 | 0.287 | 0.324 | 0.295 | 0.537 | 0.476 | 0.474 | 0.495666667 | 1.680225989 |
| O02056 | rpl-4 B0041.4 | 0.301 | 0.277 | 0.261 | 0.279666667 | 0.472 | 0.468 | 0.47 | 0.47 | 1.68057211 |
| O18236 | nuo-3 CELE_Y57G11C.12 Y57G11C.12 | 0.283 | 0.287 | 0.271 | 0.280333333 | 0.459 | 0.481 | 0.477 | 0.472333333 | 1.68489893 |
| Q9BL19 | rpl-17 Y48G8AL.8 | 0.275 | 0.258 | 0.271 | 0.268 | 0.438 | 0.453 | 0.464 | 0.451666667 | 1.685323383 |
| Q20219 | irg-7 drd-2 CELE_F40F4.6 F40F4.6 | 0.271 | 0.275 | 0.29 | 0.278666667 | 0.462 | 0.471 | 0.476 | 0.469666667 | 1.685406699 |
| Q9XW16 | pfn-1 Y18D10A.20 | 0.299 | 0.302 | 0.294 | 0.298333333 | 0.468 | 0.407 | 0.635 | 0.503333333 | 1.687150838 |
| Q22053 | fib-1 T01C3.7 | 0.261 | 0.284 | 0.28 | 0.275 | 0.505 | 0.447 | 0.44 | 0.464 | 1.687272727 |
| Q09607 | gst-36 R07B1.4 | 0.292 | 0 | 0.291 | 0.194333333 | 0.484 | 0 | 0.5 | 0.328 | 1.687821612 |
| O45934 | CELE_Y43F4B.5 Y43F4B.5 | 0.293 | 0 | 0 | 0.097666667 | 0.495 | 0 | 0 | 0.165 | 1.689419795 |
| Q22054 | rps-16 T01C3.6 | 0.287 | 0.295 | 0.27 | 0.284 | 0.457 | 0.481 | 0.502 | 0.48 | 1.690140845 |
| Q93235 | nkb-1 C17E4.9 | 0.293 | 0.277 | 0.348 | 0.306 | 0.459 | 0.469 | 0.626 | 0.518 | 1.692810458 |
| G5EF37 | pat-10 tnc-1 CELE_F54C1.7 F54C1.7 | 0.28 | 0.446 | 0.287 | 0.337666667 | 0.534 | 0.609 | 0.573 | 0.572 | 1.693978282 |
| O76371 | rpt-5 CELE_F56H1.4 F56H1.4 | 0.395 | 0.283 | 0.291 | 0.323 | 0.605 | 0.457 | 0.58 | 0.547333333 | 1.694530444 |
| P54688 | bcat-1 eca-39 eca39 K02A4.1 | 0.29 | 0.261 | 0.299 | 0.283333333 | 0.458 | 0.471 | 0.515 | 0.481333333 | 1.698823529 |
| O01925 | mboa-6 R155.1 | 0 | 0.297 | 0 | 0.099 | 0 | 0.505 | 0 | 0.168333333 | 1.7003367 |
| Q11067 | tag-320 B0403.4 | 0.288 | 0.278 | 0.272 | 0.279333333 | 0.467 | 0.485 | 0.474 | 0.475333333 | 1.701670644 |
| P90921 | asg-1 K07A12.3 | 0 | 0.255 | 0 | 0.085 | 0 | 0.434 | 0 | 0.144666667 | 1.701960784 |
| Q9N3D0 | CELE_Y54E10BR.5 Y54E10BR.5 | 0 | 0 | 0.27 | 0.09 | 0 | 0 | 0.46 | 0.153333333 | 1.703703704 |
| Q21276 | K07C5.4 | 0.288 | 0.26 | 0.277 | 0.275 | 0.455 | 0.491 | 0.46 | 0.468666667 | 1.704242424 |
| Q9N4X8 | gst-10 Y45G12C.2 | 0.295 | 0.312 | 0.266 | 0.291 | 0.502 | 0.471 | 0.515 | 0.496 | 1.704467354 |
| P48158 | rpl-23 B0336.10 | 0.315 | 0.257 | 0.253 | 0.275 | 0.458 | 0.445 | 0.504 | 0.469 | 1.705454545 |
| Q19102 | ard-1 CELE_F01G4.2 F01G4.2 | 0.282 | 0.289 | 0.253 | 0.274666667 | 0.512 | 0.466 | 0.43 | 0.469333333 | 1.708737864 |
| Q22067 | got-1.2 T01C8.5 | 0 | 0.278 | 0.316 | 0.198 | 0 | 0.518 | 0.497 | 0.338333333 | 1.708754209 |
| P91249 | col-20 CELE_F11G11.11 F11G11.11 | 0.261 | 0 | 0.279 | 0.18 | 0.457 | 0 | 0.466 | 0.307666667 | 1.709259259 |
| Q86LS4 | dao-2 CELE_M03A1.7 M03A1.7 | 0 | 0.27 | 0.254 | 0.174666667 | 0 | 0.405 | 0.492 | 0.299 | 1.711832061 |
| Q18359 | C33A12.1 | 0.248 | 0.284 | 0.243 | 0.258333333 | 0.445 | 0.427 | 0.456 | 0.442666667 | 1.713548387 |
| P34528 | K12H4.7 | 0.295 | 0.293 | 0.228 | 0.272 | 0.493 | 0.395 | 0.511 | 0.466333333 | 1.714460784 |
| Q86MI3 | CELE_Y71H10B.1 Y71H10B.1 | 0.292 | 0 | 0.306 | 0.199333333 | 0.504 | 0 | 0.523 | 0.342333333 | 1.717391304 |
| Q9N3T5 | spg-7 CELE_Y47G6A.10 Y47G6A.10 | 0.296 | 0.239 | 0 | 0.178333333 | 0.452 | 0.468 | 0 | 0.306666667 | 1.719626168 |
| Q18786 | snr-4 C52E4.3 | 0.276 | 0 | 0 | 0.092 | 0.475 | 0 | 0 | 0.158333333 | 1.721014493 |
| P41988 | cct-1 tcp-1 T05C12.7 | 0.287 | 0.269 | 0.254 | 0.27 | 0.47 | 0.454 | 0.473 | 0.465666667 | 1.724691358 |
| Q17405 | AC3.5 | 0 | 0.296 | 0 | 0.098666667 | 0 | 0.511 | 0 | 0.170333333 | 1.726351351 |

| | | | | | | | | | | |
|---|---|---|---|---|---|---|---|---|---|---|
| Q19973 | cec-5 CELE_F32E10.6 F32E10.6 | 0.257 | 0 | 0 | 0.085666667 | 0.444 | 0 | 0 | 0.148 | 1.727626459 |
| P34629 | lap-1 ZK353.6 | 0.218 | 0 | 0 | 0.072666667 | 0.377 | 0 | 0 | 0.125666667 | 1.729357798 |
| O16785 | pat-6 T21D12.4 | 0 | 0 | 0.263 | 0.087666667 | 0 | 0 | 0.455 | 0.151666667 | 1.730038023 |
| Q9XUB7 | far-6 CELE_W02A2.2 W02A2.2 | 0.256 | 0.257 | 0.295 | 0.269333333 | 0.463 | 0.454 | 0.481 | 0.466 | 1.73019802 |
| Q9XVF7 | rpl-8 rpl-2 B0250.1 | 0.275 | 0.27 | 0.281 | 0.275333333 | 0.486 | 0.46 | 0.484 | 0.476666667 | 1.731234867 |
| G5EBK3 | erm-1 C01G8.5 CELE_C01G8.5 | 0.226 | 0.32 | 0.301 | 0.282333333 | 0.532 | 0.489 | 0.446 | 0.489 | 1.731995277 |
| H2FLL1 | lea-1 CELE_K08H10.1 K08H10.1 | 0.295 | 0.283 | 0.268 | 0.282 | 0.502 | 0.469 | 0.495 | 0.488666667 | 1.73286052 |
| Q6A8K1 | act-4 CELE_M03F4.2 M03F4.2 | 0.231 | 0.29 | 0.257 | 0.259333333 | 0.463 | 0.428 | 0.458 | 0.449666667 | 1.733933162 |
| Q93791 | nid-1 CELE_F54F3.1 F54F3.1 | 0.237 | 0 | 0 | 0.079 | 0.411 | 0 | 0 | 0.137 | 1.734177215 |
| O44887 | inx-13 inx CELE_Y8G1A.2 Y8G1A.2 | 0.241 | 0 | 0 | 0.080333333 | 0.418 | 0 | 0 | 0.139333333 | 1.734439834 |
| Q9U1W1 | tfg-1 CELE_Y63D3A.5 Y63D3A.5 | 0.256 | 0.271 | 0.295 | 0.274 | 0.496 | 0.472 | 0.458 | 0.475333333 | 1.734793187 |
| G5ECC3 | rme-1 CELE_W06H8.1 W06H8.1 | 0 | 0.361 | 0 | 0.120333333 | 0 | 0.627 | 0 | 0.209 | 1.736842105 |
| O18178 | pptr-1 W08G11.4 | 0.297 | 0 | 0.349 | 0.215333333 | 0.549 | 0 | 0.573 | 0.374 | 1.736842105 |
| Q09249 | C16C10.3 | 0 | 0 | 0.271 | 0.090333333 | 0 | 0 | 0.471 | 0.157 | 1.73800738 |
| O02328 | eif-3.C T23D8.4 | 0.237 | 0.298 | 0.283 | 0.272666667 | 0.5 | 0.435 | 0.487 | 0.474 | 1.738386308 |
| P62784 | his-1 T10C6.14; his-5 F45F2.3; his-10 ZK131.4; his-14 ZK131.8; his-18 K06C4.10; his-26 ZK131.1; his-28 K06C4.2; his-31 F17E9.12; his-37 C50F4.7; his-38 K03A1.6; his-46 B0035.9; his-50 F07B7.9; his-56 F54E12.3; his-60 F55G1.11; his-64 F22B3.1; his-67 T23D8.5 | 0.286 | 0.262 | 0.276 | 0.274666667 | 0.48 | 0.48 | 0.476 | 0.478666667 | 1.742718447 |
| Q9N350 | CELE_Y55F3BR.6 Y55F3BR.6 | 0.241 | 0.315 | 0.251 | 0.269 | 0.483 | 0.452 | 0.473 | 0.469333333 | 1.744733581 |
| G5EFL5 | alp-1 CELE_T11B7.4 T11B7.4 | 0 | 0.273 | 0.351 | 0.208 | 0 | 0.567 | 0.522 | 0.363 | 1.745192308 |
| Q966I8 | pbs-1 CELE_K08D12.1 K08D12.1 | 0.251 | 0.243 | 0.306 | 0.266666667 | 0.49 | 0.456 | 0.451 | 0.465666667 | 1.74625 |
| Q10943 | arf-1.2 arf-1 B0336.2 | 0.285 | 0.28 | 0.261 | 0.275333333 | 0.473 | 0.475 | 0.496 | 0.481333333 | 1.748184019 |
| Q9N4Y8 | nuo-5 CELE_Y45G12B.1 Y45G12B.1 | 0.288 | 0.216 | 0.323 | 0.275666667 | 0.474 | 0.528 | 0.444 | 0.482 | 1.748488513 |
| Q95Y72 | dss-1 Y119D3B.15 | 0 | 0 | 0.298 | 0.099333333 | 0 | 0 | 0.522 | 0.174 | 1.751677852 |
| O16520 | erfa-1 etf-1 T05H4.6 | 0 | 0 | 0.363 | 0.121 | 0 | 0 | 0.637 | 0.212333333 | 1.754820937 |
| P46550 | cct-6 F01F1.8 | 0.273 | 0.272 | 0.266 | 0.270333333 | 0.482 | 0.463 | 0.48 | 0.475 | 1.757090012 |
| P18334 | kin-3 B0205.7 | 0.256 | 0.259 | 0.282 | 0.265666667 | 0.479 | 0.473 | 0.449 | 0.467 | 1.757841907 |
| P52872 | dad-1 F57B10.10 | 0 | 0.275 | 0 | 0.091666667 | 0 | 0.484 | 0 | 0.161333333 | 1.76 |
| Q20641 | nmy-1 CELE_F52B10.1 F52B10.1 | 0.282 | 0 | 0.267 | 0.183 | 0.453 | 0 | 0.514 | 0.322333333 | 1.761384335 |
| Q21888 | CELE_R102.2 R102.2 | 0 | 0.237 | 0 | 0.079 | 0 | 0.418 | 0 | 0.139333333 | 1.76371308 |
| Q20034 | CELE_F35D11.4 F35D11.4 | 0.228 | 0 | 0.312 | 0.18 | 0.477 | 0 | 0.476 | 0.317666667 | 1.764814815 |

| | | | | | | | | | | |
|---|---|---|---|---|---|---|---|---|---|---|
| Q20751 | iff-2 F54C9.1 | 0.249 | 0.284 | 0.289 | 0.274 | 0.537 | 0.47 | 0.446 | 0.484333333 | 1.767639903 |
| P47209 | cct-5 C07G2.3 | 0.257 | 0.301 | 0.282 | 0.28 | 0.508 | 0.477 | 0.502 | 0.495666667 | 1.770238095 |
| Q27511 | htz-1 R08C7.3 | 0 | 0.296 | 0 | 0.098666667 | 0 | 0.524 | 0 | 0.174666667 | 1.77027027 |
| O45181 | CELE_K07H8.10 K07H8.10 | 0.214 | 0.278 | 0.292 | 0.261333333 | 0.519 | 0.43 | 0.44 | 0.463 | 1.771683673 |
| O62106 | eif-6 C47B2.5 | 0.269 | 0.219 | 0 | 0.162666667 | 0.396 | 0.47 | 0 | 0.288666667 | 1.774590164 |
| Q9XWJ5 | CELE_Y51H1A.3 Y51H1A.3 | 0.26 | 0 | 0.266 | 0.175333333 | 0.473 | 0 | 0.461 | 0.311333333 | 1.775665399 |
| G4SF79 | CELE_F25E2.2 F25E2.2 | 0 | 0 | 0.264 | 0.088 | 0 | 0 | 0.469 | 0.156333333 | 1.776515152 |
| P34652 | cnx-1 ZK632.6 | 0.27 | 0.242 | 0.392 | 0.301333333 | 0.543 | 0.458 | 0.608 | 0.536333333 | 1.779867257 |
| Q9BMU4 | atln-1 4D561 CELE_Y54G2A.2 Y54G2A.2 | 0 | 0.359 | 0 | 0.119666667 | 0 | 0.641 | 0 | 0.213666667 | 1.78551532 |
| Q1XFY9 | rps-24 CELE_T07A9.11 T07A9.11 | 0.256 | 0.298 | 0.293 | 0.282333333 | 0.574 | 0.473 | 0.473 | 0.506666667 | 1.794569067 |
| A3QMC5 | rpl-34 C42C1.14 CELE_C42C1.14 | 0.254 | 0.24 | 0.281 | 0.258333333 | 0.469 | 0.473 | 0.451 | 0.464333333 | 1.797419355 |
| Q09456 | col-80 C09G5.5 | 0.264 | 0.23 | 0 | 0.164666667 | 0.438 | 0.45 | 0 | 0.296 | 1.79757085 |
| Q9U3Q6 | ugt-22 C08F11.8 CELE_C08F11.8 | 0 | 0 | 0.24 | 0.08 | 0 | 0 | 0.432 | 0.144 | 1.8 |
| Q22253 | rpn-9 CELE_T06D8.8 T06D8.8 | 0.268 | 0 | 0 | 0.089333333 | 0.483 | 0 | 0 | 0.161 | 1.802238806 |
| G5EDI2 | ret-1 CELE_W06A7.3 W06A7.3 | 0.273 | 0.249 | 0.29 | 0.270666667 | 0.466 | 0.502 | 0.496 | 0.488 | 1.802955665 |
| P17140 | let-2 clb-1 F01G12.5 | 0.236 | 0 | 0.259 | 0.165 | 0.445 | 0 | 0.448 | 0.297666667 | 1.804040404 |
| Q11117 | lmp-1 C03B1.12 | 0 | 0 | 0.286 | 0.095333333 | 0 | 0 | 0.516 | 0.172 | 1.804195804 |
| P52821 | rps-25 K02B2.5 | 0.285 | 0.248 | 0.253 | 0.262 | 0.469 | 0.473 | 0.477 | 0.473 | 1.805343511 |
| Q06561 | unc-52 ZC101.2 | 0.24 | 0.287 | 0.237 | 0.254666667 | 0.489 | 0.456 | 0.435 | 0.46 | 1.806282723 |
| O61742 | rpn-10 B0205.3 | 0 | 0 | 0.268 | 0.089333333 | 0 | 0 | 0.486 | 0.162 | 1.813432836 |
| Q9XUV0 | pbs-5 CELE_K05C4.1 K05C4.1 | 0.232 | 0.285 | 0.292 | 0.269666667 | 0.546 | 0.503 | 0.419 | 0.489333333 | 1.814585909 |
| G5EDQ2 | hmg-12 hmg-I-beta CELE_Y17G7A.1 Y17G7A.1 | 0.253 | 0.249 | 0.283 | 0.261666667 | 0.5 | 0.459 | 0.471 | 0.476666667 | 1.821656051 |
| P91017 | hpo-32 C01G8.6 CELE_C01G8.6 | 0.307 | 0 | 0.324 | 0.210333333 | 0.475 | 0 | 0.676 | 0.383666667 | 1.824088748 |
| Q09510 | mlc-4 C56G7.1 | 0.251 | 0 | 0 | 0.083666667 | 0.458 | 0 | 0 | 0.152666667 | 1.824701195 |
| Q8TA47 | CELE_Y51F10.7 Y51F10.7 | 0 | 0 | 0.354 | 0.118 | 0 | 0 | 0.646 | 0.215333333 | 1.824858757 |
| P53703 | cchl-1 T06D8.6 | 0.219 | 0 | 0 | 0.073 | 0.4 | 0 | 0 | 0.133333333 | 1.826484018 |
| Q9N362 | CELE_Y55F3AM.13 Y55F3AM.13 | 0.286 | 0.278 | 0 | 0.188 | 0.569 | 0.462 | 0 | 0.343666667 | 1.828014184 |
| O44782 | vpr-1 CELE_F33D11.11 F33D11.11 | 0 | 0.203 | 0 | 0.067666667 | 0 | 0.372 | 0 | 0.124 | 1.832512315 |
| Q19437 | upb-1 CELE_F13H8.7 F13H8.7 | 0 | 0.205 | 0.268 | 0.157666667 | 0 | 0.451 | 0.416 | 0.289 | 1.832980973 |
| Q19162 | rpl-11.2 F07D10.1 | 0.279 | 0.247 | 0.258 | 0.261333333 | 0.49 | 0.488 | 0.465 | 0.481 | 1.840561224 |
| Q23058 | ssp-9 E03H12.10; ssp-11 T28H11.6 | 0.25 | 0.213 | 0.278 | 0.247 | 0.446 | 0.477 | 0.442 | 0.455 | 1.842105263 |
| Q95Y29 | CELE_Y34B4A.6 Y34B4A.6 | 0.284 | 0.255 | 0.213 | 0.250666667 | 0.44 | 0.469 | 0.478 | 0.462333333 | 1.844414894 |
| Q8MQC6 | cpr-6 C25B8.3 CELE_C25B8.3 | 0.267 | 0.267 | 0.273 | 0.269 | 0.522 | 0.459 | 0.509 | 0.496666667 | 1.846344486 |

| | | | | | | | | | |
|---|---|---|---|---|---|---|---|---|---|
| Q9TXP0 | rps-27 F56E10.4 | 0.279 | 0.247 | 0.258 | 0.261333333 | 0.486 | 0.49 | 0.472 | 0.482666667 | 1.846938776 |
| Q9XWD1 | acs-5 CELE_Y76A2B.3 Y76A2B.3 | 0.263 | 0.251 | 0 | 0.171333333 | 0.483 | 0.467 | 0 | 0.316666667 | 1.848249027 |
| O45502 | dnj-12 CELE_F39B2.10 F39B2.10 | 0.306 | 0 | 0.292 | 0.199333333 | 0.595 | 0 | 0.513 | 0.369333333 | 1.852842809 |
| Q21086 | nst-1 K01C8.9 | 0 | 0.205 | 0 | 0.068333333 | 0 | 0.38 | 0 | 0.126666667 | 1.853658537 |
| Q2PJ97 | CELE_F52E1.14 F52E1.14 | 0.263 | 0.271 | 0.21 | 0.248 | 0.463 | 0.433 | 0.488 | 0.461333333 | 1.860215054 |
| Q18947 | ule-3 CELE_D1054.11 D1054.11 | 0 | 0 | 0.165 | 0.055 | 0 | 0 | 0.307 | 0.102333333 | 1.860606061 |
| P36609 | ncs-2 F10G8.5 | 0.289 | 0.32 | 0.276 | 0.295 | 0.563 | 0.493 | 0.592 | 0.549333333 | 1.862146893 |
| Q6AHR3 | CELE_R151.2 R151.2 | 0.188 | 0.309 | 0.255 | 0.250666667 | 0.488 | 0.461 | 0.452 | 0.467 | 1.863031915 |
| Q09657 | CELE_ZK1320.9 ZK1320.9 | 0 | 0.252 | 0 | 0.084 | 0 | 0.47 | 0 | 0.156666667 | 1.865079365 |
| Q967F1 | eif-1 CELE_T27F7.3 T27F7.3 | 0.263 | 0 | 0.25 | 0.171 | 0.527 | 0 | 0.43 | 0.319 | 1.865497076 |
| P45971 | ostb-1 T09A5.11 | 0 | 0.263 | 0 | 0.087666667 | 0 | 0.492 | 0 | 0.164 | 1.870722433 |
| Q21966 | asp-4 CELE_R12H7.2 R12H7.2 | 0.287 | 0.227 | 0.27 | 0.261333333 | 0.484 | 0.501 | 0.487 | 0.490666667 | 1.87755102 |
| Q9XUY5 | nkb-3 F55F3.3 | 0.247 | 0.282 | 0.308 | 0.279 | 0.535 | 0.48 | 0.558 | 0.524333333 | 1.879330944 |
| Q9BKQ9 | cisd-3.2 CELE_Y67D2.3 Y67D2.3 | 0.325 | 0.307 | 0.235 | 0.289 | 0.675 | 0.479 | 0.478 | 0.544 | 1.882352941 |
| Q564Q1 | gale-1 C47B2.6 CELE_C47B2.6 | 0.244 | 0 | 0.227 | 0.157 | 0.441 | 0 | 0.454 | 0.298333333 | 1.900212314 |
| O17570 | rpl-38 C06B8.8 | 0.177 | 0.264 | 0.362 | 0.267666667 | 0.56 | 0.473 | 0.496 | 0.509666667 | 1.904109589 |
| Q8MXI1 | ucr-11 CELE_F57B10.14 F57B10.14 | 0.267 | 0.211 | 0.278 | 0.252 | 0.488 | 0.488 | 0.467 | 0.481 | 1.908730159 |
| Q9XWL1 | ttr-17 CELE_Y5F2A.2 Y5F2A.2 | 0.547 | 0 | 0.236 | 0.261 | 0.441 | 0.613 | 0.441 | 0.498333333 | 1.909323116 |
| O01685 | C32F10.8 CELE_C32F10.8 | 0.255 | 0 | 0.248 | 0.167666667 | 0.491 | 0 | 0.474 | 0.321666667 | 1.918489066 |
| P55216 | cth-2 ZK1127.10 | 0.252 | 0.324 | 0.244 | 0.273333333 | 0.507 | 0.455 | 0.617 | 0.526333333 | 1.925609756 |
| C6KRN1 | sao-1 R10D12.14 | 0.26 | 0.265 | 0.261 | 0.262 | 0.497 | 0.486 | 0.534 | 0.505666667 | 1.930025445 |
| Q23571 | dbt-1 ZK669.4 | 0.248 | 0.246 | 0.227 | 0.240333333 | 0.486 | 0.44 | 0.466 | 0.464 | 1.930651872 |
| Q21568 | M28.5 | 0.228 | 0.235 | 0.261 | 0.241333333 | 0.479 | 0.456 | 0.465 | 0.466666667 | 1.933701657 |
| O17954 | CELE_LLC1.2 LLC1.2 | 0 | 0.224 | 0 | 0.074666667 | 0 | 0.434 | 0 | 0.144666667 | 1.9375 |
| O45106 | ech-5 CELE_F56B3.5 F56B3.5 | 0 | 0.27 | 0.256 | 0.175333333 | 0 | 0.449 | 0.571 | 0.34 | 1.939163498 |
| Q17348 | snr-1 Y116A8C.42 | 0.241 | 0.25 | 0.241 | 0.244 | 0.489 | 0.476 | 0.456 | 0.473666667 | 1.941256831 |
| Q65ZJ7 | C32D5.8 CELE_C32D5.8 | 0.258 | 0 | 0.244 | 0.167333333 | 0.464 | 0 | 0.512 | 0.325333333 | 1.944223108 |
| Q23635 | CELE_ZK84.1 ZK84.1 | 0 | 0 | 0.231 | 0.077 | 0 | 0 | 0.45 | 0.15 | 1.948051948 |
| Q9U2K8 | abce-1 CELE_Y39E4B.1 Y39E4B.1 | 0.299 | 0.243 | 0.229 | 0.257 | 0.48 | 0.542 | 0.481 | 0.501 | 1.949416342 |
| Q6AW03 | CELE_Y37E3.17 Y37E3.17 | 0.317 | 0.235 | 0.329 | 0.293666667 | 0.67 | 0.397 | 0.655 | 0.574 | 1.954597049 |
| Q9U238 | trap-4 CELE_Y56A3A.21 Y56A3A.21 | 0.248 | 0.241 | 0.262 | 0.250333333 | 0.506 | 0.494 | 0.479 | 0.493 | 1.969374168 |
| P19626 | mlc-2 C36E6.5 | 0.243 | 0.255 | 0.27 | 0.256 | 0.523 | 0.512 | 0.493 | 0.509333333 | 1.989583333 |
| P34714 | ost-1 sparc C44B12.2 | 0.275 | 0.232 | 0 | 0.169 | 0.53 | 0.485 | 0 | 0.338333333 | 2.001972387 |

| | | | | | | | | | | |
|---|---|---|---|---|---|---|---|---|---|---|
| G5EDY2 | wars-1 wrs-1 CELE_Y80D3A.1 Y80D3A.1 | 0.176 | 0.199 | 0.198 | 0.191 | 0.389 | 0.395 | 0.378 | 0.387333333 | 2.027923211 |
| O61792 | rpn-8 CELE_R12E2.3 R12E2.3 | 0 | 0.27 | 0.295 | 0.188333333 | 0 | 0.641 | 0.513 | 0.384666667 | 2.042477876 |
| O16521 | hpo-19 CELE_T05H4.5 T05H4.5 | 0.262 | 0 | 0 | 0.087333333 | 0.541 | 0 | 0 | 0.180333333 | 2.064885496 |
| P90789 | D2030.4 | 0 | 0.206 | 0.213 | 0.139666667 | 0 | 0.435 | 0.457 | 0.297333333 | 2.128878282 |
| A8WHP8 | plin-1 mdt-28 CELE_W01A8.1 W01A8.1 | 0 | 0.231 | 0 | 0.077 | 0 | 0.493 | 0 | 0.164333333 | 2.134199134 |
| Q20589 | CELE_F49C12.12 F49C12.12 | 0 | 0.21 | 0 | 0.07 | 0 | 0.451 | 0 | 0.150333333 | 2.147619048 |
| O18180 | mrpl-12 CELE_W09D10.3 W09D10.3 | 0.274 | 0.273 | 0.178 | 0.241666667 | 0.543 | 0.491 | 0.527 | 0.520333333 | 2.153103448 |
| Q09236 | copd-1 C13B9.3 | 0.28 | 0.239 | 0 | 0.173 | 0.72 | 0.398 | 0 | 0.372666667 | 2.154142582 |
| Q03206 | ced-10 rac-1 C09G12.8 | 0.256 | 0.224 | 0 | 0.16 | 0.518 | 0.524 | 0 | 0.347333333 | 2.170833333 |
| O44781 | CELE_F33D11.10 F33D11.10 | 0 | 0.334 | 0.289 | 0.207666667 | 0.598 | 0.382 | 0.42 | 0.466666667 | 2.247191011 |
| Q9XWV2 | Y37D8A.2 | 0 | 0 | 0.209 | 0.069666667 | 0 | 0 | 0.473 | 0.157666667 | 2.263157895 |
| Q22135 | CELE_T04A8.6 T04A8.6 | 0.198 | 0 | 0 | 0.066 | 0.459 | 0 | 0 | 0.153 | 2.318181818 |
| B7FAR9 | CELE_Y43F8B.1 Y43F8B.1 | 0 | 0.223 | 0.171 | 0.131333333 | 0 | 0.418 | 0.5 | 0.306 | 2.329949239 |
| O61709 | CELE_R119.3 R119.3 | 0 | 0.212 | 0 | 0.070666667 | 0 | 0.496 | 0 | 0.165333333 | 2.339622642 |
| Q9BL46 | CELE_Y71H2AM.11 Y71H2AM.11 | 0.255 | 0.255 | 0.186 | 0.232 | 0.521 | 0.521 | 0.592 | 0.544666667 | 2.347701149 |
| P52715 | F13D12.6 | 0.267 | 0.277 | 0 | 0.181333333 | 0.562 | 0.723 | 0 | 0.428333333 | 2.362132353 |
| G5EFV4 | hmg-1.1 CELE_Y48B6A.14 Y48B6A.14 | 0.23 | 0 | 0 | 0.076666667 | 0.549 | 0 | 0 | 0.183 | 2.386956522 |
| Q95YA9 | cas-1 CELE_F41G4.2 F41G4.2 | 0 | 0.212 | 0 | 0.070666667 | 0 | 0.507 | 0 | 0.169 | 2.391509434 |
| Q69ZI6 | CELE_T12D8.10 T12D8.10 | 0.249 | 0 | 0.223 | 0.157333333 | 0.539 | 0 | 0.591 | 0.376666667 | 2.394067797 |
| Q9XTU6 | snr-6 Y49E10.15 | 0.192 | 0 | 0 | 0.064 | 0.462 | 0 | 0 | 0.154 | 2.40625 |
| P34255 | B0303.3 | 0.305 | 0 | 0.306 | 0.203666667 | 0.514 | 0.592 | 0.411 | 0.505666667 | 2.482815057 |
| Q8I711 | gars-1 CELE_T10F2.1 T10F2.1 | 0.234 | 0 | 0 | 0.078 | 0.593 | 0 | 0 | 0.197666667 | 2.534188034 |
| O16368 | rpt-2 F29G9.5 | 0.348 | 0 | 0.229 | 0.192333333 | 0.431 | 0.545 | 0.488 | 0.488 | 2.537261698 |
| G2HK03 | pqn-87 CELE_Y57A10A.18 Y57A10A.18 | 0.414 | 0 | 0 | 0.138 | 0.405 | 0 | 0.646 | 0.350333333 | 2.538647343 |
| Q09545 | sdhb-1 tag-55 F42A8.2 | 0.33 | 0.329 | 0 | 0.219666667 | 0.57 | 0.41 | 0.799 | 0.593 | 2.699544765 |
| G5EG85 | unc-70 bgs-1 CELE_K11C4.3 K11C4.3 | 0 | 0.414 | 0 | 0.138 | 0.828 | 0.334 | 0 | 0.387333333 | 2.806763285 |
| O17352 | aagr-4 CELE_F52D1.1 F52D1.1 | 0 | 0 | 0.305 | 0.101666667 | 0.364 | 0 | 0.496 | 0.286666667 | 2.819672131 |
| O62415 | lys-1 CELE_Y22F5A.4 Y22F5A.4 | 0 | 0.16 | 0.226 | 0.128666667 | 0 | 0.622 | 0.49 | 0.370666667 | 2.880829016 |
| Q9N4L8 | lpd-5 CELE_ZK973.10 ZK973.10 | 0.241 | 0.282 | 0 | 0.174333333 | 0.454 | 0.426 | 0.641 | 0.507 | 2.908221797 |
| O02252 | pcp-3 CELE_F23B2.11 F23B2.11 | 0.124 | 0 | 0 | 0.041333333 | 0.364 | 0 | 0 | 0.121333333 | 2.935483871 |
| Q22719 | CELE_T24B8.3 T24B8.3 | 0 | 0.328 | 0.302 | 0.21 | 0.71 | 0.576 | 0.602 | 0.629333333 | 2.996825397 |

| | | | | | | | | | | |
|---|---|---|---|---|---|---|---|---|---|---|
| A7DTF0 | mrg-1 CELE_Y37D8A.9 Y37D8A.9 | 0 | 0.248 | 0 | 0.082666667 | 0 | 0.752 | 0 | 0.250666667 | 3.032258065 |
| Q23487 | CELE_ZK418.9 ZK418.9 | 0 | 0.246 | 0 | 0.082 | 0 | 0.754 | 0 | 0.251333333 | 3.06504065 |
| H2L0M0 | pod-2 CELE_W09B6.1 W09B6.1 | 0 | 0.276 | 0 | 0.092 | 0 | 0.851 | 0 | 0.283666667 | 3.083333333 |
| Q10661 | dpy-30 ZK863.6 | 0.145 | 0 | 0 | 0.048333333 | 0.452 | 0 | 0 | 0.150666667 | 3.117241379 |
| C7FZU5 | C41G7.9 CELE_C41G7.9 | 0 | 0 | 0.327 | 0.109 | 0 | 0.617 | 0.42 | 0.345666667 | 3.171253823 |
| P91250 | col-73 CELE_F11G11.12 F11G11.12 | 0 | 0.205 | 0.199 | 0.134666667 | 0.43 | 0.42 | 0.441 | 0.430333333 | 3.195544554 |
| Q22170 | ile-2 CELE_T04G9.3 T04G9.3 | 0 | 0.383 | 0 | 0.127666667 | 0.68 | 0.617 | 0 | 0.432333333 | 3.386422977 |
| G5EGH7 | lon-8 Y59A8B.20 | 0.167 | 0 | 0 | 0.055666667 | 0.606 | 0 | 0 | 0.202 | 3.628742515 |
| G5EE74 | nud-1 CELE_F53A2.4 F53A2.4 | 0 | 0 | 0.312 | 0.104 | 0.774 | 0 | 0.396 | 0.39 | 3.75 |
| O45864 | CELE_T27E9.2 T27E9.2 | 0 | 0 | 0.263 | 0.087666667 | 0.609 | 0 | 0.379 | 0.329333333 | 3.756653992 |
| Q9XWM1 | oig-2 CELE_Y38F1A.9 Y38F1A.9 | 0.21 | 0 | 0.193 | 0.134333333 | 0.435 | 0.59 | 0.496 | 0.507 | 3.774193548 |
| Q21313 | epi-1 K08C7.3 | 0.247 | 0.207 | 0 | 0.151333333 | 0.463 | 0.668 | 0.604 | 0.578333333 | 3.821585903 |
| Q23089 | xpo-1 CELE_ZK742.1 ZK742.1 | 0 | 0 | 0.116 | 0.038666667 | 0 | 0 | 0.448 | 0.149333333 | 3.862068966 |
| O62432 | dpy-4 CELE_Y41E3.2 Y41E3.2 | 0.067 | 0.157 | 0.131 | 0.118333333 | 0.502 | 0.468 | 0.442 | 0.470666667 | 3.977464789 |
| O17287 | tomm-22 W10D9.5 | 0 | 0.135 | 0 | 0.045 | 0 | 0.548 | 0 | 0.182666667 | 4.059259259 |
| Q09289 | C56G2.7 | 0 | 0.242 | 0 | 0.080666667 | 0 | 0.512 | 0.486 | 0.332666667 | 4.123966942 |
| Q9XTT4 | ani-1 Y49E10.19 | 0 | 0.07 | 0 | 0.023333333 | 0 | 0.299 | 0 | 0.099666667 | 4.271428571 |
| H2KYI5 | C06G3.5 CELE_C06G3.5 | 0.24 | 0 | 0 | 0.08 | 0.482 | 0 | 0.558 | 0.346666667 | 4.333333333 |
| H2L076 | ketn-1 CELE_F54E2.3 F54E2.3 | 0 | 0 | 0.249 | 0.083 | 0 | 0.594 | 0.513 | 0.369 | 4.445783133 |
| Q93896 | mct-3 CELE_M03B6.2 M03B6.2 | 0 | 0.315 | 0 | 0.105 | 1 | 0.42 | 0 | 0.473333333 | 4.507936508 |
| P34183 | hrs-1 syh-1 T11G6.1 | 0.393 | 0 | 0 | 0.131 | 0.486 | 0.63 | 0.683 | 0.599666667 | 4.577608142 |
| Q8T3G5 | egl-30 CELE_M01D7.7 M01D7.7 | 0 | 0.121 | 0 | 0.040333333 | 0 | 0.555 | 0 | 0.185 | 4.58677686 |
| Q9BL07 | Y54F10AM.8 | 0 | 0.172 | 0 | 0.057333333 | 0 | 0.509 | 0.432 | 0.313666667 | 5.470930233 |
| Q93618 | F27D4.4 | 0 | 0.297 | 0 | 0.099 | 0.673 | 0.446 | 0.63 | 0.583 | 5.888888889 |
| O02153 | grd-14 CELE_T01B10.2 T01B10.2 | 0 | 0 | 0.132 | 0.044 | 0.45 | 0 | 0.4 | 0.283333333 | 6.439393939 |
| Q22192 | ttr-14 CELE_T05A10.3 T05A10.3 | 0 | 0 | 0.157 | 0.052333333 | 0.684 | 0 | 0.503 | 0.395666667 | 7.560509554 |
| Q94360 | nduf-7 W10D5.2 | 0.076 | 0 | 0 | 0.025333333 | 0.572 | 0 | 0.611 | 0.394333333 | 15.56578947 |
